# Analytical strategies to include the X-chromosome in variance heterogeneity analyses: evidence for trait-specific polygenic variance structure

**DOI:** 10.1101/306654

**Authors:** Wei Q. Deng, Shihong Mao, Anette Kalnapenkis, Tõnu Esko, Reedik Mägi, Guillaume Paré, Lei Sun

## Abstract

Genotype-stratified variance of a quantitative trait could differ in the presence of gene-gene or gene-environment interactions. Genetic markers associated with phenotypic variance are thus considered promising candidates for follow-up interaction or joint location-scale analyses. However, as in studies of main effects, the X-chromosome is routinely excluded from ‘whole-genome’ scans due to analytical challenges. Specifically, as males carry only one copy of the X-chromosome, the inherent sex-genotype dependency could bias the trait-genotype association, through sexual dimorphism in quantitative traits with sex-specific means or variances. Here we investigate phenotypic variance heterogeneity associated with X-chromosome SNPs and propose valid and powerful strategies. Among those, a generalized Levene’s test has adequate power and remains robust to sexual dimorphism. An alternative approach is sex-stratified analysis but at the cost of slightly reduced power and modeling flexibility. We applied both methods to an Estonian study of gene expression quantitative trait loci (eQTL; *n*=841), and two complex trait studies of height, hip and waist circumferences, and body mass index from multi-ethnic study of atherosclerosis (MESA; *n*=2,073) and UK Biobank (UKB; *n*=327,393). Consistent with previous eQTL findings on mean, we found some but no conclusive evidence for *cis* regulators being enriched for variance association. SNP rs2681646 is associated with variance of waist circumference (*p*=9.5E-07) at X-chromosome-wide significance in UKB, with a suggestive female-specific effect in MESA (*p*=0.048). Collectively, an enrichment analysis using permutated UKB (*p*<1/10) and MESA (*p*<1/100) datasets, suggests a possible polygenic structure for the variance of human height.

## Introduction

Several recent reports have examined autosomal genetic loci contributing to phenotypic variance (as opposed to mean) for a wide range of complex traits (Pare, Cook, Ridker, & Chasman, 2010; Shungin et al., 2017; Yang et al., 2012), and corresponding methodology development remains an active area of research (Aschard, Zaitlen, Tamimi, Lindstrom, & Kraft, 2013; Cao, Wei, Bailey, Kauwe, & Maxwell, 2014; Deng, Asma, & Pare, 2014; Deng & Pare, 2011; Hulse & Cai, 2013; Soave et al., 2015; Soave & Sun, 2017; Struchalin, Dehghan, Witteman, van Duijn, & Aulchenko, 2010; Sun, Elston, Morris, & Zhu, 2013). One possible reason for such phenotypic variance and SNP genotype association, or variance heterogeneity, is that genotype-stratified variances of a trait differ in the presence of gene-gene (*G*x*G*) or gene-environment (*G*x*E*) interactions; both referred to as *G*x*E* hereinafter. For example, rs1358030 (*SORCS1*) was shown to interact with treatment type affecting HbA1c levels in Type 1 Diabetes subjects (Paterson et al., 2010). And indeed, in a proof-of-principle study where the treatment information was intentionally masked, the SNP was then demonstrated to be associated with variance of HbA1c (Soave et al., 2015). Conversely, because direct *G*x*E* modeling may not be feasible in an initial whole-genome scan, the question was then raised as to whether SNPs having effects on the variance of a trait make good candidates for follow-up interaction testing (Shungin et al., 2017). For instance, rs7202116 (*FTO* as the nearest gene) was significantly associated with variance of body mass index (BMI) (Yang et al., 2012), and at the same locus, rs1121980 (*FTO)* showed evidence for a statistical interaction with physical activity influencing the mean of BMI (Ahmad et al., 2013; Kilpelainen et al., 2011); it is worth noting that un-modeled interaction induces variance heterogeneity, but the causes of variance heterogeneity are multifaceted (Cao et al., 2014; Dudbridge & Fletcher, 2014; Pare et al., 2010; Soave et al., 2015; Struchalin et al., 2010; Sun et al., 2013; Wood et al., 2014). In practice, although it is possible that an interacting SNP has a stronger effect on variance than on mean, as in the case of rs12753193 (*LEPR*) interacting with BMI in the prediction of CRP levels in the absence of detectable main effect (Pare et al., 2010), a more powerful approach to selecting association candidates is to jointly evaluate their mean and variance effects (Aschard, Hancock, London, & Kraft, 2010; Cao et al., 2014; Soave et al., 2015).

Despite enthusiasm to discover SNPs with variance effects and the availability of statistical tests, variance heterogeneity has not been formally explored for SNPs on the X-chromosome (XCHR). As in the conventional ‘genome-wide’ (mean) association studies (Wise, Gyi, & Manolio, 2013), the reluctance to include XCHR is due to analytical challenges (Konig, Loley, Erdmann, & Ziegler, 2014; Wise et al., 2013). They range from technical difficulties in genotype calling to statistical complexities in imputation and association (e.g. model uncertainty involving random or skewed X-inactivation (Carrel & Willard, 2005; Ross et al., 2005; Tukiainen et al., 2017; Wang, Yu, & Shete, 2014) and sex as a potential confounder). Solutions to overcome some of these challenges had been provided, but all in the context of genetic association analysis of main effects (Chen, Craiu, Strug, & Sun, 2019; Chen, Craiu, & Sun, 2018; D. Clayton, 2008; D. G. Clayton, 2009; Hickey & Bahlo, 2011; Wang et al., 2014; Özbek et al., 2018).

Here we focus on understanding the impact of the inherent sex-genotype dependency on variance heterogeneity association analysis, and when the trait of interest has sex-specific mean or variance values for males and females. In practice, sexual dimorphism is consistently observed. For example, based on the UK Biobank (UKB) (Sudlow et al., 2015) and Multi-Ethnic Study of Atherosclerosis (Bild et al., 2002) (MESA) data, height displays a sex-specific difference in mean, hip circumference differs in variance, while body mass index (BMI) and waist circumference contrast in both mean and variance between males and females (Fig 1). These empirical patterns of sexual dimorphism vary according to the underlying physiology of the trait, which might or might not be related to genes. Thus, association analyses of phenotypic mean or variance with XCHR SNPs could be biased if these potential sex-specific main or variance effects were not appropriately accounted for.

**Figure 1.**
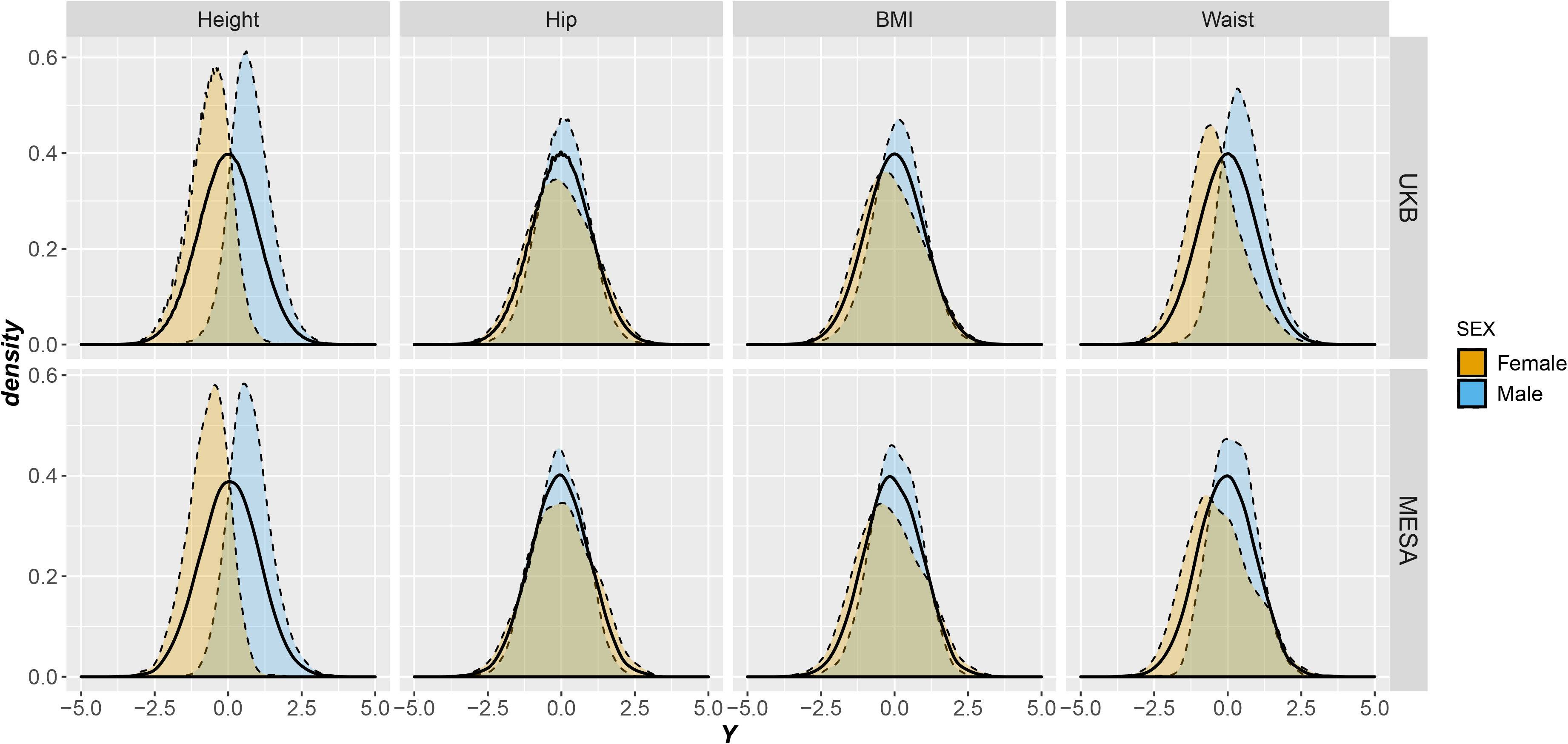
Empirical examples of sexual dimorphism: quantitative trait distribution stratified by sex. Phenotype data from UK Biobank (top row) and the Multi-Ethnic Study of Atherosclerosis (bottom row) were used to illustrate the possible types of sexual dimorphism as characterized by a location shift in mean of height, a scale difference in the variance of hip circumference, and changes in both mean and variance of waist circumference and BMI. Each trait (*Y*) was inversely transformed so the overall distribution (solid curve) is normal with mean 0 and variance 1. Areas under the sex-stratified distributions (dashed curves) are colored by blue for male and orange for female, respectively.

For an autosomal SNP, evaluating differences in phenotypic variance across the three genotype groups can be readily achieved by the classical Levene’s test for variance heterogeneity (Levene, 1960). SNPs with significant variance association *p*-values are then selected as likely candidates for follow-up interaction studies. However, the same strategy to prioritize SNPs on XCHR can be problematic, because sex-specific mean and variance differences could create spurious variance heterogeneity unrelated to the putative *G*x*E* interactions of interest. Thus, the correct formulation of variance test is dependent on a proper formulation of sex effect with respect to both mean and variance.

In this paper, we explicitly model the possible sources of confounding related to sex, and propose two general testing strategies that strike a balance between power and robustness against various model uncertainties. Using extensive simulations, we demonstrate the danger of directly applying autosomal methods to the XCHR that would otherwise be suitable for testing variance heterogeneity, and we conclude that special consideration for sex-genotype dependence must be made for the XCHR to maintain correct type I error rates. Application studies include identifying SNPs associated with variances of height, BMI, hip and waist circumference using the UK Biobank (UKB) and multi-ethnic study of atherosclerosis (MESA) data, as well as detecting loci associated with variance of expression quantitative traits using data from the Estonian Genome Center at the University of Tartu (EGCUT) cohort (Leitsalu et al., 2015; Metspalu, 2002; Westra et al., 2013).

## Methods

### Variance of a quantitative trait by genotype in the presence of genetic interactions

Of interest is a quantitative trait *Y*, assumed to be (approximately) normally distributed or had been inversely transformed to resemble a normal distribution. Without loss of generality, first consider the following linear model for the ‘*true’* association relationship between *Y* and an autosomal SNP,

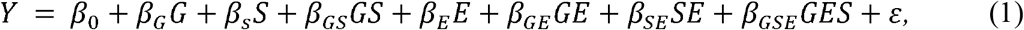

where *G* denotes the SNP genotype coded additively (Hill, Goddard, & Visscher, 2008) with respect to the number of the minor allele 0, 1 and 2 for *bb*, *Bb* and *BB* as in convention, *S* is the male sex indicator variable (e.g. *S* = 0 for females and *S* = 1 for males), *E* ∼ *N*(0, 1) is a standardized continuous covariate following the classical *G*-*E* independence assumption (Lindstrom, Yen, Spiegelman, & Kraft, 2009), and the error term *ε* ∼ *N*(0, 1) is independent of *G*, *S* and *E*. The minor allele frequency (MAF) of *G* is assumed to be the same for male and females; sex-specific MAF affects the naïve methods and we will return to this point in the Discussion section.

Under these assumptions, it is possible to identify autosomal SNPs potentially involved in *G*x*E* or high-order interactions, without having to measure *E* directly, through detecting phenotypic variance associated with *G* via the *working* model of *Y* ∼ *G*. Note that the analytical context here is that direct *G*x*E* (or *G*x*G*) modeling may not be possible (e.g. *E* may not be known or measured precisely) or desirable (e.g. due to computational or multiple hypothesis testing concerns for whole-genome *G*x*G* scans). To see the rationale behind the working model, with the additional assumption of conditional independence between *E* and *S* conditional on *G*, one can show that the conditional variance of *Y* on *G* is,

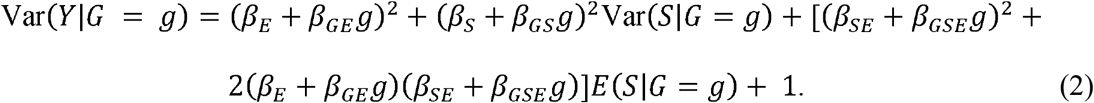

Since *S* is independent of *G* for an autosomal SNP, Pr(*S*|*G* = *g*) is constant across *g* = 0, 1 and 2, so are E*(S|G=g)* and Var*(S|G=g)*. Thus, if *β*_*GS*_= *β*_*GE*_= *β*_*GSE*_= 0, expression (2) can be reduced to a constant with respect to *G*:

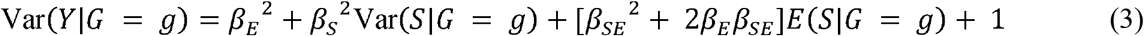

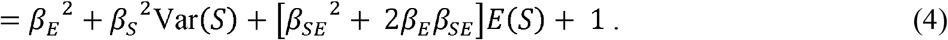

Conversely, variation in Var(*Y*|*G*=*g*) across *G* suggests that at least some of the (un-modeled) interaction terms involving *G* (i.e. *β*_*GS*_, *β*_*GE*_ and *β*_*GSE*_) are non-zero. This was precisely the motivation behind the original idea of using Levene’s test to identify variance heterogeneity induced by the underlying but un-modeled *G*x*E* interaction (Pare et al., 2010).

### X-chromosome (XCHR) specific challenges for variance tests

The same approach to draw similar conclusions for XCHR SNPs, however, is questionable, because Pr(*S*|*G* = *g*) is no longer constant in *G* and expression (3) cannot be further reduced to (4). For example, Clayton’s approach (Clayton, 2008) suggests coding the *bb*, *Bb* and *BB* genotypes in females as 0, 1 and 2 and the *b* and *B* genotypes as 0 and 2 in males, the *G=1* group contains only females. Similarly, without considerations for the X-chromosome, using the usual autosomal coding of 0, 1 and 2 in females and 0 and 1 in males, the *G*=*2* group then contains only females. Thus, omitting the sex indicator *S* from the covariates can bias the conclusion through sexual dimorphism as seen in Fig 1.

Consider the simplest case of no interaction effects at all (*β*_*GS*_ = *β*_*GE*_ = *β*_*SE*_ = *β*_*GSE*_ = 0) nor environmental main effect (*β*_*E*_ = 0), but there is a sex main effect (*β*_*S*_ ≠ 0, i.e. the sex-stratified phenotypic means differ between males and females), then expression (3) is reduced to

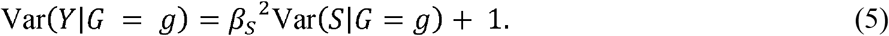

Thus, in the absence of any interactions that involve *G*, there is a spurious phenotypic variance heterogeneity across levels of *G* through a non-zero sex main effect (*β*_*S*_), or through a sex-environment interaction effect (*β*_*SE*_) if present as in equation (3). Severity of the confounding depends on the discrepancy between the two sex-stratified trait distributions (real data in Fig 1 and conceptual data in Fig 2A-D), as well as on the strength of correlation between sex and the observed genotype, which in turn depends on the MAF and proportions of males and females in a sample (details in S1 Text).

**Figure 2.**
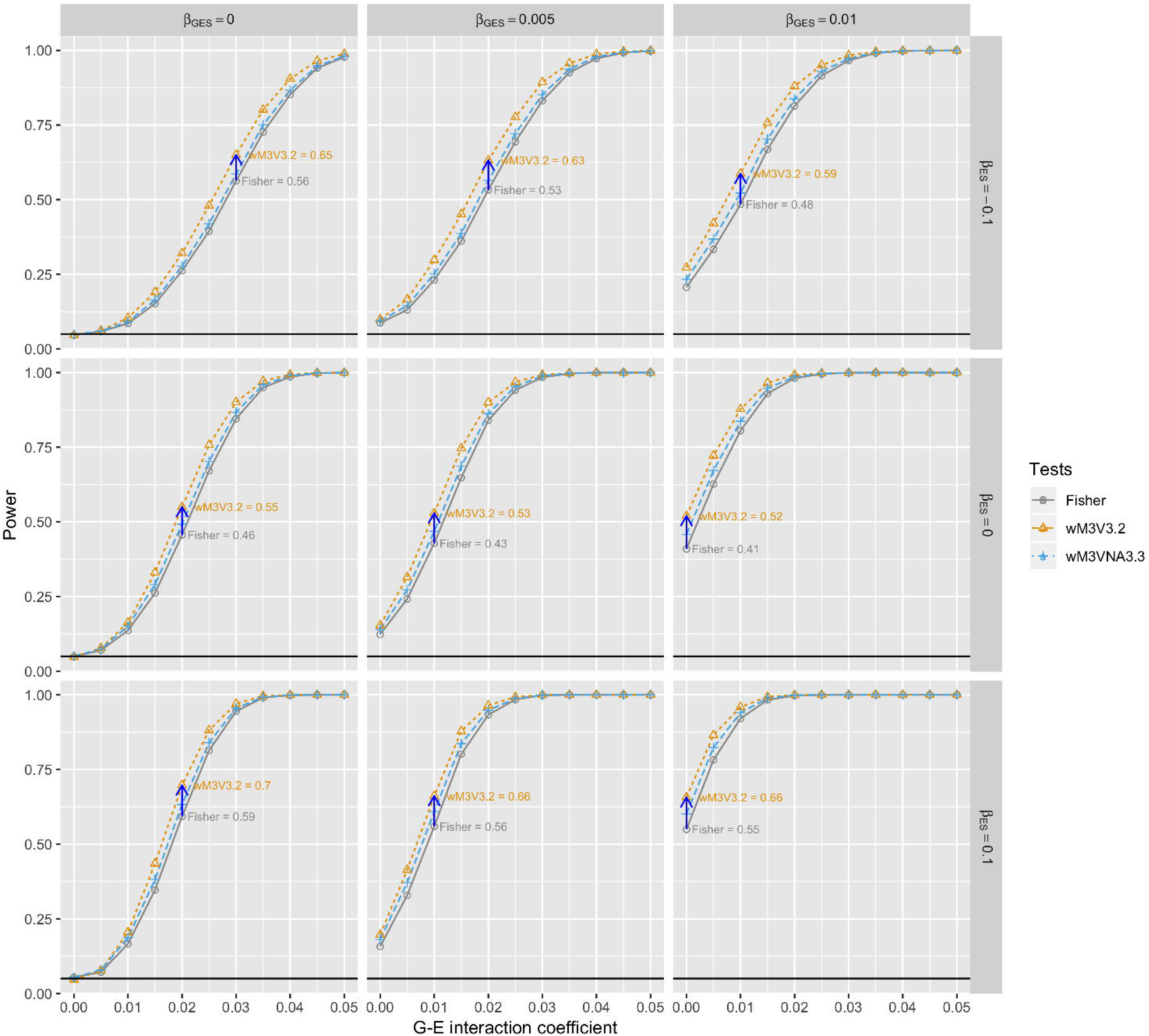
Defining null and alternative hypotheses for X-chromosome variance heterogeneity test allowing for sexual dimorphism. Upper panel: Figures A-D showcase the different types of conceptually *null* distributions, where the variance of a quantitative trait does not vary across the different genotype groups, but is subjected to a possible sex-specific difference in either mean (B), variance (C) or both (D). The black curve is for the overall distribution, and without loss of generality, the orange curve is for female and the blue curve is for male the same as in Fig 1. Lower panel: Figures E-H represent the respective *alternative* distributions. The different genotype groups are marked by different line types and visible only under the alternative conditions when there is phenotypic variance heterogeneity among the genotype groups.

To avoid spurious variance heterogeneity signals, alternative approaches are needed to quantify variance differences induced by *G*x*E* or higher order interactions involving *G*. To this end, it is important to appropriately define the null hypothesis of variance homogeneity that corresponds to an absence of phenotypic variance associated with genotype while allowing for variance (and mean) to differ between males and females (Fig 2A-D).

We also note that the different coding schemes (i.e. 0-1 or 0-2 in males) are not meant to equate or model the biological effect of X-inactivation (or the absence of it), as the relationship between allele dosage in tissues and phenotypic mean at an organism level depends on a number of biological factors (Deng, Berletch, Nguyen, & Disteche, 2014). Rather, they are used to build association tests that are analytical appropriate for different biological scenarios. Further, it has been noted that, for association analysis of phenotypic mean, the use of an additively coded genotype alone might not be sufficient for the XCHR (Chen et al., 2019; Özbek et al., 2018). Thus, we will be considering models that account for both sex main and genotype-sex interaction effects.

### X-chromosome (XCHR) variance heterogeneity tests

Here we consider various analytical strategies to assess phenotypic variance associated with genotypes of XCHR SNPs, including naïve methods that directly apply the original Levene’s test to different genotype groups, and an alternative approach that utilizes a generalized Levene’s test derived from a two-stage regression framework (Gastwirth, Gel, & Miao, 2009; Levene, 1960; Soave & Sun, 2017). The recommend methods have been implemented as an open-source R program (Web resources).

#### Naïve methods: apply Levene’s test to three or five genotype groups

The original Levene’s test for variance heterogeneity treats an autosomal genotype *G* as a categorical variable (Gastwirth et al., 2009; Levene, 1960) and examines any variance difference in trait *Y* amongst the three possible genotype groups. A direct application to XCHR, however, is problematic. Because sex *S* is inherently correlated with *G* of a XCHR SNP, so any potential correlation between *S* and *Y* (e.g. as observed in human height) would create the classic case of confounding. Consider the null situation where *G* is *not* associated with the variance of *Y* as in the top panel of Fig 2. Assume the 0-2 coding of *G* in males was used (same conclusion for the 0-1 coding in males), the *Bb* group contains only females and its variance would be the same as 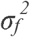, reflected by variance of the orange curve in the figure. In contrast, the other two groups (*bb+b* and *BB+B*) contain both males and females, and their respective variance values, involving both the orange and blue curves, depend on sex-specific means (*μ*_*m*_ and *μ*_*f*_) and variances (*σ*_*m*_^*2*^ and 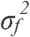), as well the proportion of males in each group. Thus, in the presence of sexual dimorphism, either in mean (Fig 2B), variance (Fig 2C), or both (Fig 2D), there would be spurious variance heterogeneity resulting in increased false positive rates.

As an alternative, one may be tempted to treat each genotype and sex combination as one group, resulting in a total of 5 groups. Indeed, this five-group strategy does not induce spurious association in the presence of sex-specific mean effect (*μ*_*m*_ ≠ *μ*_*f*_ as in Fig 2B). However, it is not difficult to see that the problem remains when there is a sex-specific variance effect (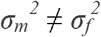 as in Fig 2C or 2D).

#### Fisher’s method: combine sex-stratified Levene’s test

Sex-stratified analysis provides a practical strategy whereby variance heterogeneity is assessed separately in males (Levene’s test for two groups) and females (Levene’s test for three groups). Fisher’s method can then be used to combine the two independent *p*-values (Derkach, Lawless, & Sun, 2013). Though a sex-stratified analysis does not allow direct *G*x*S* modeling, it is robust to various forms of sexual dimorphism as seen in Fig 2B-D, and it does not require considerations for dosage compensation related to random X-inactivation.

#### Model-based generalized Levene’s test: account for sex-specific mean and variance effects via two-stage regression models

Since we defined the null hypothesis in terms of phenotypic variance heterogeneity induced by (un-modeled) *G*x*E* interactions while allowing for sexual dimorphism (Fig 2B-D), a preferred method should explicitly account for the effect of sex on the phenotype of interest. We consider the generalized Levene’s test in a flexible two-stage regression framework. In essence, stage one regresses *Y* on *G* and obtains the absolute residual *d* (i.e. the *absolute* value of residuals given by the difference between the observed and fitted *Y* values). Stage two regresses *d* on *G* again and tests the slope using the ANOVA *F*-test. This approach has been shown to be equivalent to testing variance heterogeneity in *Y* associated with *G*, because the expectation of *d* linearly depends on variance of *Y* (Gastwirth et al., 2009; Soave & Sun, 2017). It is possible to regress *d*^*2*^ or other function forms of the absolute residual, however, *d* approximately follows a folded-normal distribution and is more robust to model assumptions (Gastwirth et al., 2009; Levene, 1960).

The generalized Levene’s test has been used to study autosomal SNPs with more complex data structures including genotype group uncertainty (e.g. imputed SNPs) or sample dependency (e.g. correlated family members) (Soave & Sun, 2017). For XCHR analysis, the implementation requires additional care because it is not immediately clear whether *S* (or *G*x*S*) should be included in both stages. For a comprehensive evaluation, we consider all combinations of the following two-stage models:

**Stage One: Mean models,**

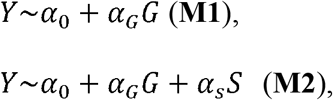

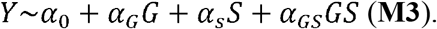
**Stage Two: Variance models,**

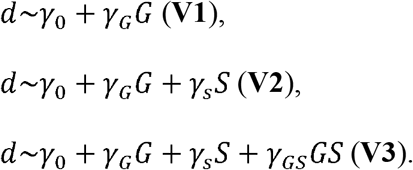

Note that a non-additive variance model (VNA) in stage two may be considered:

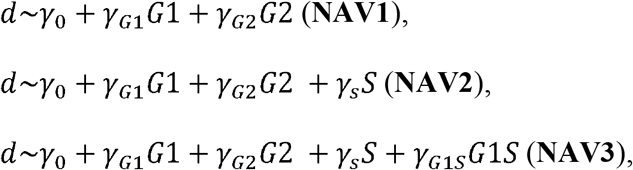

where *G*1 and *G*2 are indicator variables, respectively, for the *Bb* and *BB+B* groups under the 0-2 coding scheme in males, or alternatively for the *Bb+B* and *BB* groups assuming a 0-1 coding in males.

The models in stage one are only used to calculate residuals, using either the traditional ordinary least squares (OLS) or the recommended least absolute deviations (LAD); LAD is more robust to data with asymmetric distributions or low genotype counts in a specific group (Chen et al., 2019; Hines & Hines, 2000; Soave & Sun, 2017). The goal of this stage is to remove any *Mean* effects associated with the covariates included in the model (i.e. *G*, *S* or *G*x*S*); thus models in stage 1 are denoted as M1, M2 or M3.

Test for *Variance* heterogeneity is achieved in stage two (V1, V2 or V3), by testing *H*_*o*_: *γ*_*G*_ = *0* or *H*_*o*_: *γ*_*G*_ = *γ*_*GS*_ = *0* via the standard regression *F*-test, where the model is fitted using OLS for independent samples or generalized least squares for dependent samples.

Note that in both stage one and two, the homoscedasticity assumption is violated whenever the residual variance is not constant across the stratified predictors (i.e. whenever *β*_*SE*_ ≠ 0, see S1 Text). In this case, the estimated regression coefficients are still unbiased, but the inference will be inflated due to underestimated variance of the estimates. Since our main objective in the first stage is not inference, but rather estimating an accurate location shift for each subgroup, the violation does not invalidate our procedure to remove any mean effect. However, in stage two, the inference on *γ*_*G*_ and *γ*_*GS*_ would be affected if the homoscedastic residuals from stage one were not properly adjusted. A simple fix to this problem is to use a weighted response *d*_*w*_ = ***1***_*(S=0)*_*d/s*_*f*_ + ***1***_*(S=1)*_ *d/s*_*m*_, where *s*_*f*_ and *s*_*m*_ denote the sample standard deviations of *Y* in females and males, respectively (S1 Text).

The model-based regression approach includes a total of 24 strategies, with 18 M+V two-stage models, V3 and VNA3 also allow a two and three degrees of freedom (d.f.) test for each stage one model, respectively (summarized in S1 Table). Based on the earlier discussion, it is expected that mean modeling strategies omitting *S* (i.e. M1) would be sensitive to sex-specific mean effect (e.g. Fig 2B or 2D). Meanwhile, variance testing strategies omitting *S* (i.e. response without weights) are anticipated to be sensitive to sex-specific variance effect (e.g. Fig 2C or 2D). For completeness of our empirical validation, we first examined the three genotype group-based naïve strategies and all 24 model-based strategies in simulation studies, as well as Fisher’s method, and then focused on the more robust ones in applications.

### Simulation studies

Note that although *G* could be coded 0-2 or 0-1 in males, these two strategies are generally highly correlated leading to similar association results (Chen et al., 2018). Further, it has been shown recently that when *G*x*S* interaction is included in the mean model the two coding schemes are equivalent in terms of association analysis of phenotypic mean (Chen et al., 2019). Thus, for a more focused study here the genotype for males was coded 0-1 and simulated with the same minor allele frequency as in females.

A joint mean and variance test can be more powerful than testing for variance heterogeneity alone, but the power of the joint test depends on the individual components (Soave et al., 2015). Therefore, here we focus on comparing the different variance-testing strategies as outlined above, recommending the most robust yet powerful method that is also suitable for the joint location-scale test.

#### Simulations for evaluating type I error control - design I based on model (1)

A sample of 5,000 females and 5,000 males were simulated, and the MAF was fixed at 0.2; other sample sizes and MAFs led to qualitatively similar results. The genotype-phenotype relationship was generated according to model (1), where the environmental variable *E* ∼ *N*(0, 1) was used in generating observed phenotypic values but assumed not being available for the actual association analysis.

The null scenarios were defined by the absence of interaction effects for *G*x*E* and *G*x*E*x*S*, so the quantitative trait for each null scenario was generated assuming *β*_*GE*_ = *β*_*GES*_ = 0 in model (1). A SNP could have a *G* main effect, but it does not affect the phenotypic variance of interest, which is induced by un-modeled *β*_*GE*_ and *β*_*GES*_ in the working model, so *β*_*G*_ = 0 without loss of generality. Note that the naïve variance methods could also pick up a non-zero *G*x*S* interaction effect if *β*_*GS*_ ≠ 0, but *β*_*GS*_ itself in fact can be directly tested as gender information is routinely collected (or reliably inferred from the available genotype data). Thus, *β*_*GS*_ is not related to the variance heterogeneity of interest here and was set to be zero, *β*_*GS*_= 0. For the remaining parameters, *β*_*0*_ = 0, *β*_*E*_= 0 or 0.5, *β*_*S*_= 0 or 0.5, and *β*_*SE*_= 0, −0.25 or 0.25, giving a total of 12 scenarios.

The different scenarios roughly fall into four categories, corresponding to the four conceptual sex-stratified distributions as shown in Fig 2A-D. For example, sexual dimorphism was introduced via *β*_*S*_ and *β*_*SE*_, where a none-zero *β*_*S*_ allows for *sex*-specific mean effect (Fig 2B and 2D) while a non-zero *β*_*SE*_ allows for *sex*-specific variance effect (Fig 2C and 2D). Note that both *β*_*S*_ and *β*_*SE*_ are independent of the *genotype*-specific variance effect to be identified, which is absent in the null cases. The number of simulated replicates was 10,000 so that estimates of the empirical type I error rates within ±0.5% of the nominal rate of 5% were considered satisfactory.

#### Simulations for evaluating type I error control – design II based on sex-stratified mean and variance

The null scenarios based on model (1) may not fully capture the extremes of sexual dimorphism, thus we further simulated trait values directly according to sex-specific distributions using means *μ*_*m*_ and *μ*_*f*_) and variances (*σ*_*m*_^*2*^ and 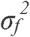) that mimic the values observed in inverse-normally transformed BMI, height, hip and waist circumference from MESA (S2 Table). The simulated traits, generated independent of any genotypes, were then tested for variance association with genotypes XCHR SNPs from the MESA dataset after LD pruning based on a window size of 50, a step size of 10 and a variance inflation factor of 3 among females using PLINK (Purcell et al., 2007) and filtering by a minimum count of 30 observations in the five sex-genotype stratified groups, resulting in a sample of 2,073 individuals and 2,100 SNPs.

#### Simulations for evaluating power

Only strategies with satisfactory type I error control were considered for power evaluation. We focused on model-based design I where the power directly depends on the size of *G*x*E* and *G*x*E*x*S* interaction effects and has a clearer genetic interpretation than design II.

A sample of 60,000 females and 50,000 males were simulated to mimic the typical sizes observed in large genome-wide studies, and the MAF was fixed at 0.2. Under model (1), *β*_*GE*_ was varied from 0 to 0.05 with a 0.005 incremental increase, and combined with a possible three-way interaction *β*_*GES*_ of 0, 0.005, or 0.01. Other parameter values were *β*_*0*_ = *β*_*G*_ = *β*_*GS*_ = 0, *β*_*G*_ = 0.2, *β*_*E*_ = 0.5, and *β*_*SE*_ = 0, −0.1 or 0.1.

### Applications

Robust variance testing strategies for X-chromosome SNPs that also had reasonable power performance were then applied to real data. Only reportedly unrelated and ethnically Caucasian individuals were included, and diabetic individuals were excluded based on electronic medical records in the UK Biobank (Sudlow et al., 2015), and based on blood glucose level greater than 7 mmol/L in MESA (Bild et al., 2002). All quantitative traits were quantile-normally transformed (Pare et al., 2010; Shungin et al., 2017); see the Discussion section for a discussion on applying quantile transformation to the original data. The significance level for discovery was set at a nominal level of 5% with Bonferroni correction for the total number of XCHR SNPs examined.

#### The UK Biobank (UKB) data

The available genotyped XCHR SNPs were filtered based on whether they were in pseudo autosomal region, a minimal sample count of 30 across the five sex-genotype groups, and genotype missing rate lower than 0.01. In total, 13,621 XCHR SNPs on the Caucasian samples (178,743 females and 148,620 males) were analyzed, and the XCHR-wide significance level was 3.7E-06.

#### The Multi-Ethnic Study of Atherosclerosis (MESA) data

The genotype data in MESA, available from dbGap (Study accession: phs000209.v10.p2), were filtered similarly as the UKB data. In total, 12,205 XCHR SNPs on the Caucasian sample (1,003 females and 1,070 males) were analyzed. We did not perform a multi-ethnic analysis with all ethnicities combined. Instead, we focused on the Caucasian subset and used it to corroborate findings from the UKB data.

#### Estonian Genome Center at the University of Tartu (EGCUT) cohort

We sought to discover XCHR SNPs influencing the variance of expression traits, as variability of gene expression has been suggested to be associated with genetic variants on autosomes (Hulse & Cai, 2013). The recommended strategies were applied to a sample of 413 male and 421 female Estonians across 648 gene expression traits that had gone through standard quality control procedures and further inversely normal transformed. After filtering using the same criteria as the UKB data, 4,034 XCHR SNPs were analyzed for variance association with each of the 648 gene expression traits, resulting in a total number of 2,614,032 tests and a global significance level of 1.9E-08.

## RESULTS

### Simulation studies

As expected, the naïve Levene’s test, with either a three-level genotype factor *G* or a five-level genotype and sex factor *G*-*S*, resulted in grossly inflated empirical type I error rates in almost all scenarios except for when *β*_*S*,_ and *β*_*SE*_ were all set to zero, or equivalently, in the absence of any sexual dimorphism (Table 1).

**Table 1.** Empirical type I error rates of variance heterogeneity tests under simulation design I. A quantitative trait was simulated according to simulation design I based on linear regression model (1) with coefficient values specified above such that Condition 1 captures the null scenario of no sexual dimorphism, i.e. no sex-specific mean nor variance differences as depicted in Fig 2A; Condition 2 corresponds to the conceptual null scenario in Fig 2B with the presence of sex-specific means via a non-zero *β*_*S*_; Conditions 3A and 3B correspond to Fig 2C, representing a sex-specific variance difference through a non-zero *β*_*SE*_, the *S*x*E* interaction effect, where the environmental effect *β*_*E*_ takes a value of either 0 (Condition 3A) or 0.5 (Condition 3B). Similarly, Conditions 4A and 4B correspond to Fig 2D with sexual dimorphism in both means and variances, with the absence and presence of environmental effect *β*_*E*_, respectively. The total sample size was 10,000 with 5,000 females and 5,000 males, and the MAF was 0.2. The nominal α-level was set to 0.05 and the empirical type I error rates were calculated based on 10,000 simulated replicates. Those empirical type I error rates exceeding 5%±0.5% were in bold. Testing strategies that showed satisfactory type I error controls were underlined, and details of the testing strategies are provided in the text and summarized in S1 Table.

|  | Condition |  | Condition |  | Condition |  | Condition |  | Condition |  | Condition |  |
| --- | --- | --- | --- | --- | --- | --- | --- | --- | --- | --- | --- | --- |
|  | 1 |  | 2 |  | 3A |  | 3B |  | 4A |  | 4B |  |
| $\beta_E$ | 0 | 0.5 | 0 | 0.5 | 0 | 0 | 0.5 | 0.5 | 0 | 0 | 0.5 | 0.5 |
| $\beta_S$ | 0 | 0 | 0.5 | 0.5 | 0 | 0 | 0 | 0 | 0.5 | 0.5 | 0.5 | 0.5 |
| $\beta_{SE}$ | 0 | 0 | 0 | 0 | 0.5 | -0.5 | 0.5 | -0.5 | 0.5 | -0.5 | 0.5 | -0.5 |
| Lev3 | 0.0453 | 0.0445 | 0.0621 | 0.0905 | 0.7747 | 0.8271 | 0.8184 | 0.2584 | 0.8092 | 0.7868 | 0.7489 | 0.1494 |
| Lev5 | 0.0240 | 0.0235 | 0.0230 | 0.1033 | 1 | 1 | 1 | 1 | 1 | 1 | 1 | 1 |
| Female | 0.0484 | 0.0512 | 0.0534 | 0.0508 | 0.0505 | 0.0524 | 0.0495 | 0.0515 | 0.0521 | 0.0464 | 0.0489 | 0.0497 |
| Male | 0.0495 | 0.0494 | 0.0514 | 0.0481 | 0.0506 | 0.0471 | 0.0503 | 0.0509 | 0.0507 | 0.0513 | 0.0524 | 0.0485 |
| Fisher | 0.0501 | 0.0492 | 0.0521 | 0.0543 | 0.0504 | 0.0503 | 0.0518 | 0.0509 | 0.0501 | 0.0498 | 0.0491 | 0.0483 |
| wM3V3.2 | 0.048 | 0.0486 | 0.0508 | 0.051 | 0.0506 | 0.0487 | 0.0505 | 0.0519 | 0.0481 | 0.0509 | 0.0478 | 0.050 |
| wM3VNA3.3 | 0.050 | 0.0499 | 0.0502 | 0.0528 | 0.0516 | 0.0515 | 0.0516 | 0.0499 | 0.0487 | 0.0505 | 0.0488 | 0.0493 |

For generalized two-stage Levene’s tests, appropriate choices of the mean model in stage one and variance test in stage two should explicitly account for any effects of sex to avoid inflating the phenotype-genotype association test statistics. Thus, as expected, any strategies involving M1 had inflated type I error rates, where the degrees of departure from the nominal α-level varied according to sizes of the unadjusted sex mean or variance effects (S3 Table). The remaining strategies also have reasonably controlled type I error rates when considering design II where sexual dimorphism was more extreme (S4 Table). The sex-stratified approach, as expected, gave correct empirical type I error rates in females and males separately, and subsequently in the combined sample via Fisher’s methods, under both design I (Table 1) and design II (S4 Table).

The reason for performance similarity between M2 and M3 is because the model was generated (and correctly modeled) under the 0-1 coding and *β*_*G*_ = 0. Interestingly, when there is a strong genotypic main effect (*β*_*G*_ ≠ 0), an increased variance in the female heterozygote *Bb* group could be observed as a result of unknown X-inactivation (Ma, Hoffman, & Keinan, 2015). Indeed, additional simulation studies confirmed that variance heterogeneity *p*-values given by models based on M2 were influenced by increased variance due to inconsistent coding choices used when simulating and analyzing the data. Though the model-based wM3V2 and wM3V3 still maintained corrected type I error rates, the only strategies remained consistent irrespective of coding choices are wM3V3.2, wM3VNA3.3, and Fisher’s method (S5 Table). This is consistent with the recent results that including the *G*x*S* interaction in models studying phenotypic mean difference between genotype groups can analytically overcome the X-inactivation uncertainty (Chen et al, 2019).

In terms of statistical power among testing strategies with reasonable control of type I error rate and invariant to coding choices, wM3V3.2 and wM3VNA3.3 have better power than Fisher’s method in all scenarios (Figure 3 and S1 Figure). Unsurprisingly, wM3V3.2 has the best performance when the underlying variance effect is additive, most notably when the effect is small (Figure 3). When only non-additive variance effect is present, wM3VNA3.3 has the best performance, but wM3V3.2 quickly becomes competitive as the additive effects start to deviate from zero (S1 Figure). Thus, we recommend the model-based wM3V3.2 and wM3VNA3.3, which were then applied to the three application datasets along with the complementary sex-stratified Fisher’s method.

**Figure 3.**
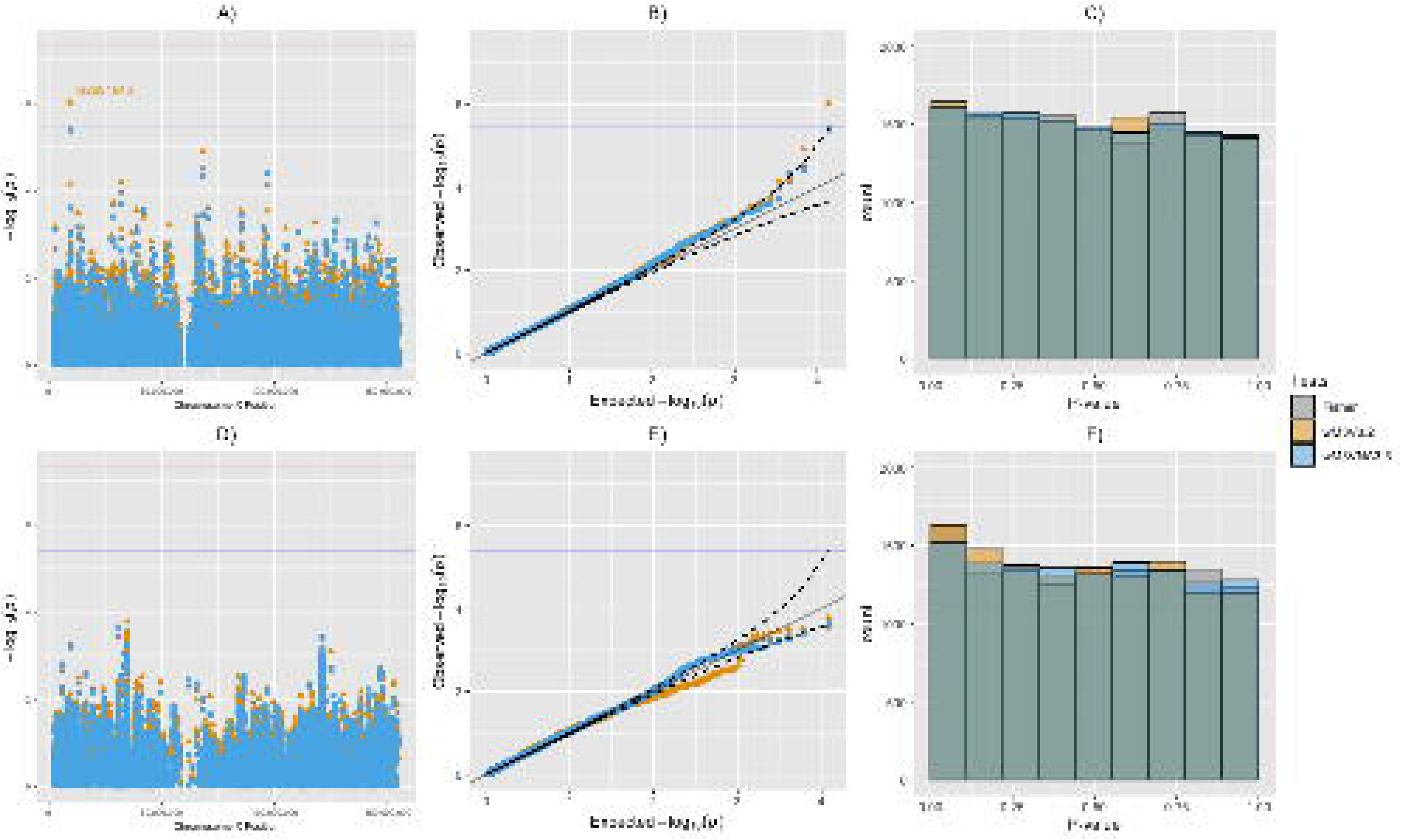
Statistical power to detect variance heterogeneity induced by gene-environment interactions using generalized Levene’s tests and Fisher’s method. The total sample size was 110,000 with 60,000 females and 50,000 males, and the MAF was 0.2 in both females and males. The type I error rate was set to 0.05 and the power were calculated based on 10,000 simulated replicates. The genotype-phenotype relationship was generated according to model (1), where the null parameter values are set to *β*_*0*_ = *β*_*G*_ = *β*_*GS*_, *β*_*S*_ = 0.2, *β*_*E*_ = 0.5, and *β*_*SE*_ = 0, −0.1 or 0.1. A total of 9 scenarios were considered to assess the power of variance heterogeneity tests induced by either the gene-environment interaction with *β*_*GE*_ from 0 to 0.05 with 0.005 incremental increases, combined with a three-way interaction between genotype, (unobserved) environmental covariate, and sex with *β*_*GES*_ = 0, 0.005 or 0.01. The maximum difference between wM3V3.2 and Fisher’s method is shown in the plots below indicated by the lengths of blue arrows under each scenario.

### Applications

As expected, for traits with sexual dimorphism, the Lev3 and Lev5 strategies produced *p*-values of varying levels of departure from the reference uniform distribution (S2-5 Figures). Using the proposed wM3V3.2, rs2681646 (*TBLIX*) was X-chromosome-wide significant (*p* < 3.7E-06) for waist circumference (*p* = 9.45E-07; Figure 4 and S6 Table) in UKB. The same SNP is marginally significant in females (Levene’s test *p* = 0.048) using the MESA data. We searched for SNPs within ±5Kb of rs2681646 at a minimal linkage disequilibrium (LD) of *r*^*2*^ > 0.7, and found 3 out of the 5 SNPs with wM3V3.2 test *p* < 0.05 in the MESA data (S6 Figure). In addition, one of the SNPs with the lowest *p*-value of testing variance heterogeneity in height, rs1474563, though not X-chromosome wide significant (wM3V3.2 *p* = 5.87E-06; S6 Table), has been shown to have a strong marginal effect on human height with a reported marginal association *p* = 3.0E-06 (Gudbjartsson et al., 2008). A joint location-scale approach (*p* = 4.54E-10) would have selected it as a good candidate for direct gene-environment interaction testing, if data on the environmental variable had been collected.

**Figure 4.**
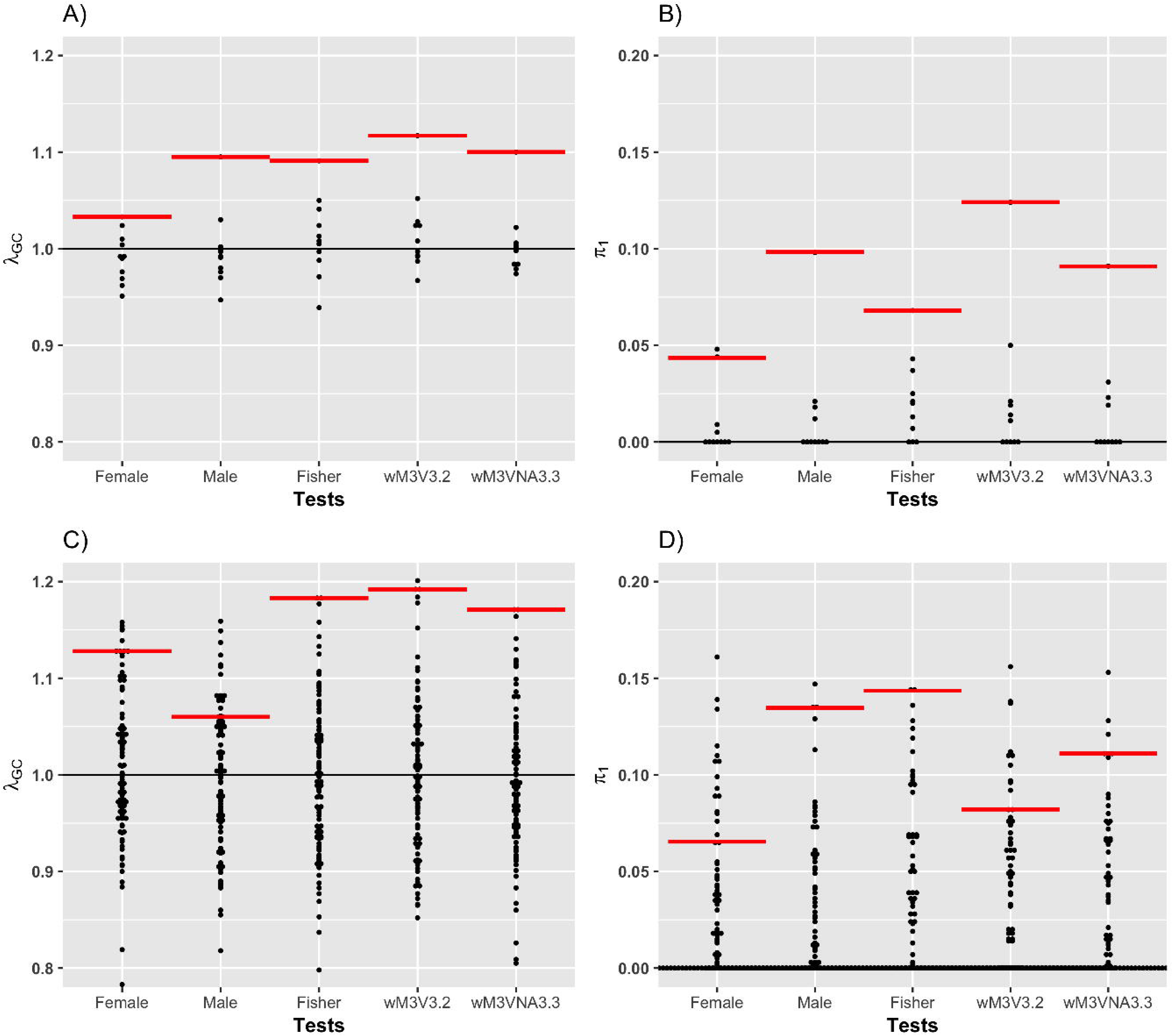
XCHR-wide variance heterogeneity test results for waist circumference using the UK Biobank (upper panel) and MESA (lower panel) data. For each XCHR SNP passing quality controls, the variance heterogeneity *p*-value was calculated using the sex-stratified Fisher’s method (grey color), the model-based strategies wM3V3.2 (orange color) and wM3VNA3.3 (blue color). Manhattan plots (A and D), quantile-quantile plots (B and E), and histograms (C and F) of the *p*-values using data from the UK Biobank are shown on the top row, and on the bottom row for the Multi-Ethnic Study of Atherosclerosis. In Fig 3A, SNP rs2661646 (wM3V3.2 test *p* = 9.5E-07) was annotated for passing the XCHR-wide significance at 3.7E-06 in the UK Biobank data.

It is established in the literature that SNPs associated with phenotypic means are more likely to interact with environmental factors (Shungin et al., 2017), thus we performed a literature search of all X-chromosome SNPs associated with height, waist and hip circumference, waist-hip-ratio, BMI, and type 2 diabetes, at a suggestive significance of *p* < 1E-03. We found 32 out of 40 SNPs to be available or in LD (*r*^*2*^>0.8) in the UK Biobank data (S7 Table). Among the 32 SNPs, 8 and 4 with significant variance heterogeneity *p*-values (< 0.05/32 = 1.6E-03), respectively, based on wM3V3.2 and wM3VNA3.3, and 5 based on Fisher’s method; the 4 and 5 SNPs are all part of the 8 SNPs identified by the wM3V3.2 test (S7 Table). These results strengthen the motivation of using variance heterogeneity to prioritize SNPs likely to be involved in *G*x*E* interactions.

Although there were no additional X-chromosome-wide significant SNPs in UKB (S7-9 Figures; S6 Tables), the overall distributions of the *p*-values suggest enrichment of variance-associated variants for some of the traits. For example, the estimated genomic lambda *λ*_*GC*_ based on the wM3V3.2 variance test for height was 1.12 and it was 1.10 based on Fisher’s method (S8 Table). For waist circumference, the estimated *λ*_*GC*_ based on wM3VNA3.3 was 1.08 and 1.07 based on Fisher’s method. Strikingly, *λ*_*GC*_ for hip circumference was sex-specific with a clearly more prominent *λ*_*GC*_ = 1.05 in females than *λ*_*GC*_ = 1.01 in males. The proportion of truly associated SNPs, π_1_ (estimated using methods described in (Storey & Tibshirani, 2003)), also suggested enrichment in height and waist circumference (S8 Table).

To benchmark the observation of *λ*_*GC*_=1.12 and π_1_=12.4% for height in UKB, and *λ*_*GC*_=1.19 and π_1_=10.9% in MESA, we performed a permutation-based analysis (Figure 5). For the UKB data, we permutated the individual phenotypic data, within the two sex strata, independently, 10 times, out of consideration for the heavy computation involved (139 hours on an Intel(R) Xeon(R) E5-2697 v3 @ 2.60GHz machine with >500Gb memory running 5 cores simultaneously). For each permutated null dataset, we applied the wM3V3.2 test (and wM3VNA3.3 and Fisher’s method) and calculated the corresponding *λ*_*GC*_ and π_1_ values (Figures 5-A and 5-B). We then applied the same analysis to the MESA data but increased the permutation replicates to 100 because of the smaller sample size (Figures 5-C and 5-D).

**Figure 5.**
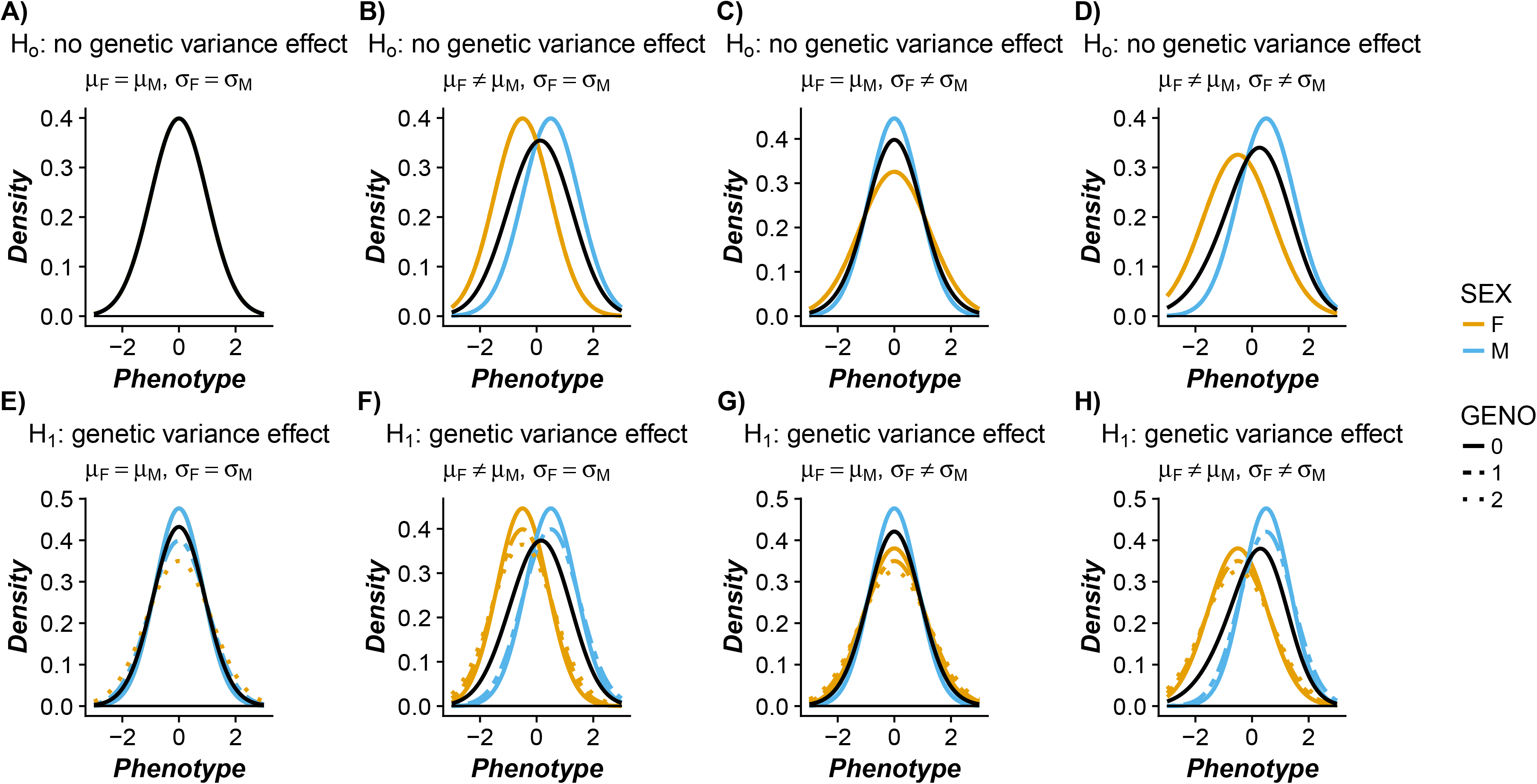
Enrichment analyses of polygenic variance structure for human height using the UKB and MESA data. The permutation study was done using the UK Biobank genetic data released in May 2017 (Figures A and B) and the Multi-Ethnic Study of Atherosclerosis (Figures C and D). The sample size used in the current analysis is *n* = 327,393 for UKB and *n* = 2,073 for MESA. The permuted dataset was obtained by sampling each quantitative trait without replacement within each sex, independently, 10 times for UKB and 100 times for MESA. The genomic control lambda *λ*_*GC*_ (Figures A and C) and proportion of truly associated SNPs *π*_*1*_ (Figures B and D) were computed in each of the permuted datasets and shown as a black dot under each test. The red line represents the estimate for the originally observed data, and the black horizontal line represents the reference line at *λ*_*GC*_ = 1 or *π*_*1*_ = 0.

The permutation-based null *λ*_*GC*_ values based on the wM3V3.2 test, as expected, centered around 1 in both the UKB and MESA permuted datasets (Figure 5). For the UKB sample, none of the 10 permuted values under the null of no variance heterogeneity was bigger than the observed *λ*_*GC*_=1.12, while for the MESA sample, only one out of the 100 replicates was bigger than the observed *λ*_*GC*_=1.19; results are similar for π_1_ estimates and for the wM3VNA3.3 and Fisher’s methods (Figure 5). Collectively, these results suggest a potential polygenic variance structure for human height.

For the eQTL analysis, we observed various forms of sexual dimorphism in expression traits. In total, 182 out of the 648 expression traits had *p* < 0.05 based on either a *t*-test of equality of means or an *F*-test for equality of variance between the two sexes (S10 Figure). Among the eQTLs, the top five variance-associated SNPs belonged to three genes, *NGFRAP1, TSC22D3*, and *ZMYM3* (S11-13 Figures), but no SNPs passed the strict Bonferroni correction at *p* < 1.9E-08. There was no apparent enrichment of association globally over all SNP-expression 2,614,032 (= 648 × 4,034) *p*-values (S14 Figure). However, upon further investigation based on stratifying SNPs and gene expression pairs according to whether they were *cis* or *trans* acting (using a physical distance of 5Mbps from the start and the end of the gene for each expression trait), we found that the estimated proportion of truly associated SNP-expression pairs appear to be slightly higher for SNPs in *cis*, as compared to those in *trans* (S15 Figure); this result is consistent with earlier results in testing mean differences in gene expression (Huang, Rangrej, Paterson, & Sun, 2007). The estimated *λ*_*GC*_ were 1.013, 1.000, and 0.998 for *cis*-acting pairs using, respectively, wM3V3.2, wM3VNA3.3, and Fisher’s methods, while for SNP-expression pairs in *trans* the estimates were 0.993, 0.993, and 0.983. Meanwhile, the estimated π_1_ were 0.020, 0.017, and 0.010 for *cis*-acting pairs using, respectively, wM3V3.2, wM3VNA3.3, and Fisher’s methods, while for SNP-expression pairs in *trans* the estimates were 0, 0, and 0.002, suggesting some enrichment, but additional studies are needed to establish convincing evidence for enrichment of variance-associated eQTLs.

## DISCUSSION

This work was motivated by the recent call to include X-chromosome (XCHR) in ‘whole-genome’ scans (Wise et al., 2013), as well as the recent development of identifying autosomal SNPs associated with phenotypic variance (Deng et al., 2014; Shungin et al., 2017; Soave & Sun, 2017; Struchalin et al., 2010; Yang et al., 2012). To pave the way for future XCHR-wide study of variance heterogeneity and subsequent joint location-scale test (Aschard et al., 2013; Cao et al., 2014; Soave et al., 2015), we examined a catalogue of analytical strategies and recommended two robust and powerful approaches. We emphasize the importance of recognizing sex as an inherent confounder in analyzing XCHR variants that contribute to phenotypic variance heterogeneity, particularly for traits displaying sexual dimorphism with either sex-specific means or variances, or both; this also holds for the traditional association analysis of XCHR variants studying their effects on phenotypic mean (Konig et al., 2014).

Between the three strategies that are robust to sexual dimorphism, Fisher’s method to combine sex-specific Levene’s *p*-values is intuitive, but it comes at the cost of power, as well as modeling flexibility. For example, adjusting for effects of other covariates that may differ between males and females. Through exploiting the recently proposed generalized Levene’s test based on a two-stage regression approach, the model-based wM3V3.2 and wM3VNA3.3 test have better power and can directly account for sex main effect as well as an unobserved *S*x*E* interaction effect. The model-based regression testing strategy can also adjust for other covariates such as principal components (Price et al., 2006). In conclusion, we recommend in practice to apply both wM3V3.2 and wM3VNA3.3 to identify additive and non-additive signals, which could be leveraged to prioritize SNPs for subsequent interaction or joint location-scale analyses.

The naïve strategies that directly test for variance heterogeneity across either the classical three genotype groups or the sex-stratified five groups are inadequate with grossed inflated type I error rates in the presence of any sexual dimorphism (Table 1). Under the non-additive coding scheme, Lev3 is equivalent to M1VNA1 in which the main effect of sex is not account for in stage one; while the setup of Lev5 is equivalent to the model-based M3VNA3, but erroneously testing *γ*_*G*1_ = *γ*_*G*2_ = *γ*_*S*_ = *γ*_*G*1*S*_ = 0 capturing the variance heterogeneity due to sex *γ*_*S*_. These observations clearly revealed the source of bias inherent in the naïve methods. Indeed, the naïve approach would not be suitable for complex traits with any sexual dimorphism, which was the case for all the four complex traits studied here (S2-5 Figures). For example, the Levene’s test three-group *p*-value for variance of waist circumference is less than 1.0E-100 for all SNPs studied in UKB, while the same test produces *p*-values less than 1.0E-10 in MESA. This provides strong evidence that the naïve Levene’s method cannot be reliably applied, and *post-hoc* adjustment is difficult. Further, any false positive findings in the discovery data would be also falsely replicated in the replication data as long as the pattern of sexual dimorphism is consistent.

The additive coding for *G* in stage one is believed to sufficiently capture the genetic main effect (Hill et al., 2008), while it may not be the case for analysis of variance. The ambiguous genotype grouping under unknown X-inactivation status adds another layer of complexity for non-additive variance models. In fact, the choice of reference allele coding matters for the XCHR when the mean association model does not include the sex main effect as shown recently (Chen et al., 2019). Further, a variance difference in the female homozygote group could be observed as a result of a strong marginal effect coupled with unknown X-inactivation (Ma et al., 2015).

Additional simulation results under model (1) with a non-zero genetic main effect suggested inflated type I error rate derived from M2 when the underlying X-inactivation status was not accounted for (S5 Table). Though in applications, the genetic main effect would have to be extremely large for the M2 model-based variance test resulting in different conclusions from M3. Interesting, it has been shown that the 0-2 and 0-1 codings lead to identical (mean) association results if the *G*x*S* interaction term is included in the model and being tested (Chen et al., 2019), which explains the consistent performance of wM3V3.2 and wM3VNA3.3 irrespective of coding choices. Further, skewed X-inactivation can be analytically represented by an over-dominant term (Chen et al., 2019), thus we also considered a non-additive variance model. It is of future interest to study the effects when considering a genotypic model for the stage 1 mean models as well.

In practice, inverse-normal quantitle-based transformation is often applied to the original phenotypic data to ensure normality but at the cost of statistical power. Although Levene’s test itself is robust to certain types of non-normal distribution, particularly when data are non-normal but symmetric (Soave & Sun, 2017), there are arguments both for and against the transformation with respect to the interpretation of variance heterogeneity (Struchalin et al., 2010; Sun et al., 2013). For example, a significant variance heterogeneity could be the result of “a mean-variance relationship induced by an inappropriate measurement scale for the phenotype” (Soave et al., 2015). This was empirically observed and cautioned by Pare et al. (Pare et al., 2010) and Yang et al. (Yang et al., 2012) in the analyses of C-reactive protein level and body mass index, respectively, where both traits had been inverse-normally transformed. On the same note, the dependence between mean and variance created by an inappropriately chosen scale could also induce false positives in a subsequent joint location-scale test if the correlation between the individual tests were not accounted for appropriately (Soave et al., 2015).

Analyses of imputed SNPs warrant some considerations. For example, real data show that thirds of the imputed SNPs were lost on chromosome X because of being monomorphic” and “16% of the imputed SNPs were available after QC on chromosome X” (Konig et al., 2014). For SNPs passing standard quality control, three approaches can be used. One is the ‘hard call’ approach using discrete values for genotypes with the highest posterior genotype probabilities. Another would be the ‘dosage’ approach assuming an additive model. And the third is to incorporate the genotype probabilities into the model. The proposed method uses the generalized Levene’s variance testing framework and is amenable to all three approaches, because the method relies on regression where the genotype predictor(s) can be discrete, dosage or probabilities. However, similar to mean association tests (Acar & Sun, 2013), genotype uncertainty decreases power regardless of the specific variance testing approaches.

Since variance testing requires larger sample size than mean testing, detecting individual variance signals that are significant at the XCHR-wide or genome-wide level requires studies of very large size that might only be viable through meta-analysis. Meta-analyses of variance heterogeneity (Deng et al., 2014) for XCHR variants can be conducted in parallel to that of a single study incorporating the analytical strategies proposed for autosomal variants (Deng et al., 2014). Note that Levene’s test statistic is asymptotically χ^2^ (with degrees of freedom equal to the number of groups subtracted by 1) distributed without an apparent ‘direction of effect’, so the traditional meta-analysis that combines the weighted (directional) *Z*-values for testing mean effect is not immediately applicable here. A non-directional meta-analysis of Levene’s test has been proposed in the context of autosomal SNPs (Deng, Asma, & Pare, 2014), but the model-based approach proposed here is more applicable for meta-analysis of XCHR variants to combine the regression coefficients from stage two jointly (Manning et al., 2011).

Similar to a polygenic model proposed for association studies of main effects, it is possible that a large proportion of genetic variants, though not individually detectable, could collectively contribute to variance heterogeneity in certain complex traits (International Schizophrenia et al., 2009; Yang et al., 2011). Though a polygenic structure for the *mean* of human height is well-known, the observed *λ*_*GC*_ and π_1_ values as compared to those computed from permutations suggest enrichment and point to a possible XCHR polygenic inheritance model for the *variance* of human height, which means that height could be potentially enriched for gene-environment interactions. Some have suggested that X-linked genes contribute to the sex-specific architecture of complex traits (Weiss, Pan, Abney, & Ober, 2006), yet the amount of contribution from XCHR SNPs involved in possible *G*x*E* or higher-order interactions is unclear. Results from this study call for new developments of the broad-sense heritability estimation methods that can incorporate variance loci, as well as quantify their contributions to sex-specific heritability.

## Acknowledgements

The authors thank Professor Radu V. Craiu and Mr. Bo Chen for helpful discussions and two external reviewers for their constructive suggestions. The authors would also like to thank Silva Kasela for her help on optimizing the eQTL analysis. We are thankful to all the participants of the Multi-Ethnic Study of Atherosclerosis, UK Biobank, Estonian Genome Center at the University of Tartu cohort. This research was funded by the Canadian Institutes of Health Research (CIHR, MOP-310732) and the Natural Sciences and Engineering Research Council of Canada (NSERC, RGPIN-04934 and RGPAS-522594) to LS. WQD is supported by NSERC Alexander Graham Bell Canada Graduate Scholarship and Ontario Graduate Scholarship.

## Web Resources

The proposed method has been implemented and is currently hosted on github as an open-source and user-friendly R package (https://github.com/WeiAkaneDeng/Xvarhet). Note that as the proposed tests are regression-based, they could be easily implemented in any other standard statistical software language preferred by the readers.

## Supporting Information

Supporting information includes 15 figures, 8 tables, and theoretical derivations for the proposed method.

## Notes

#### Summary of Updates

Section on Methods and Results updated to reflect the correction imposed in the variance heterogeneity tests; Figure 3 updated to show more clearly the advantage of our method; Figure 5 was updated using the entire UKB dataset instead of the previous release. The supplemental files are reduced and include additional simulation results.

