## Supplementary material for "Analytical strategies to include the X-chromosome in variance heterogeneity analyses: evidence for trait-specific polygenic variance structure": S1 Text; S1 Table

**S1 Figure. Statistical power to detect variance heterogeneity under non-additive genetic models using generalized linear tests and Fisher's method.** The total sample size was 2,000 with 1,000 females and 1,000 males, and the MAF was 0.2 in both females and males. The type I error rate was set to 0.05 and the power were calculated based on 10,000 simulated replicates. The pattern of sex-stratified distribution considered including those with no difference, a difference in mean, variance or both. The genotype-phenotype relationship was generated under the alternative hypothesis to assess the power of variance heterogeneity tests with the effect of one reference allele on standard deviation of quantitative trait from 0 to 0.1 with 0.01 incremental increases, and combined with the possible non-additive effect of increased variance in female heterozygote group at $\delta_1 = 0$, 0.005 or 0.01.

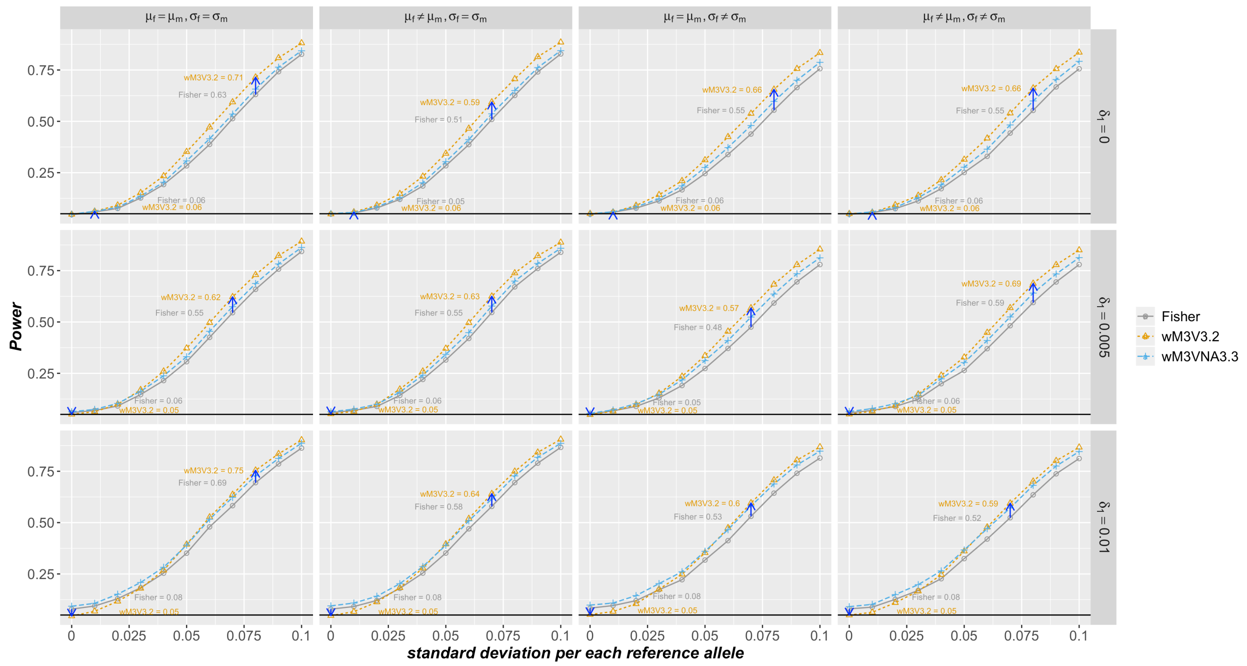

**S2 Figure. Quantile-quantile plots and histograms of variance heterogeneity p-values using Naïve Strategies for height.** For each XCHR SNP, the variance heterogeneity *p*-value was calculated using Levene's test without modifications stratified by either the genotype (Lev3) or the combinations of genotype and sex (Lev5). Quantile-quantile plots (A and C), and histograms (B and D) of the *p*-values were produced using data from the Multi-Ethnic Study of Atherosclerosis are shown on the top row, and on the bottom row for UK Biobank.

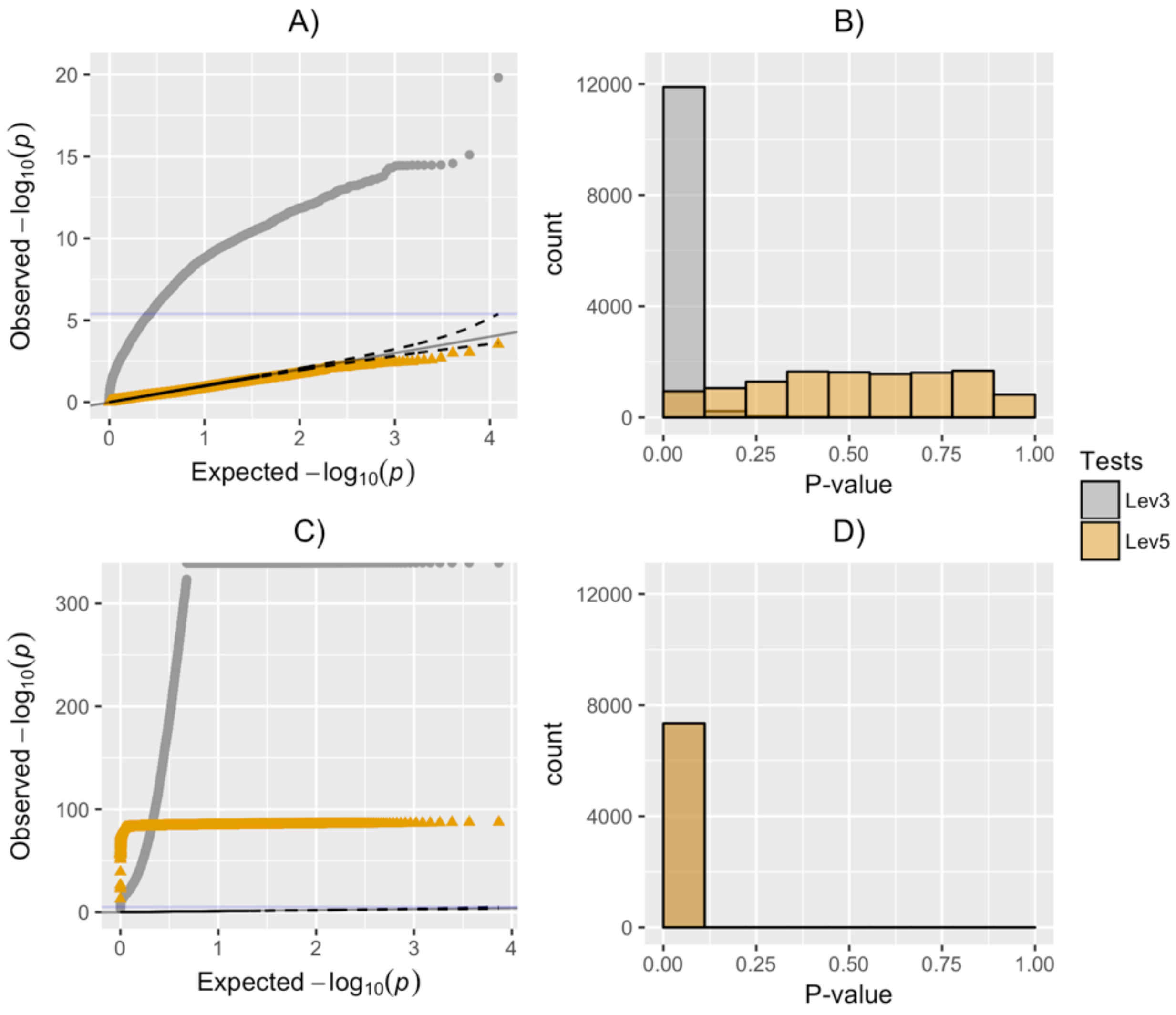

**S3 Figure. Quantile-quantile plots and histograms of variance heterogeneity p-values using Naïve Strategies for hip circumference.** For each XCHR SNP, the variance heterogeneity *p*-value was calculated using Levene's test without modifications stratified by either the genotype (Lev3) or the combinations of genotype and sex (Lev5). Quantile-quantile plots (A and C), and histograms (B and D) of the *p*-values were produced using data from the Multi-Ethnic Study of Atherosclerosis are shown on the top row, and on the bottom row for UK Biobank.

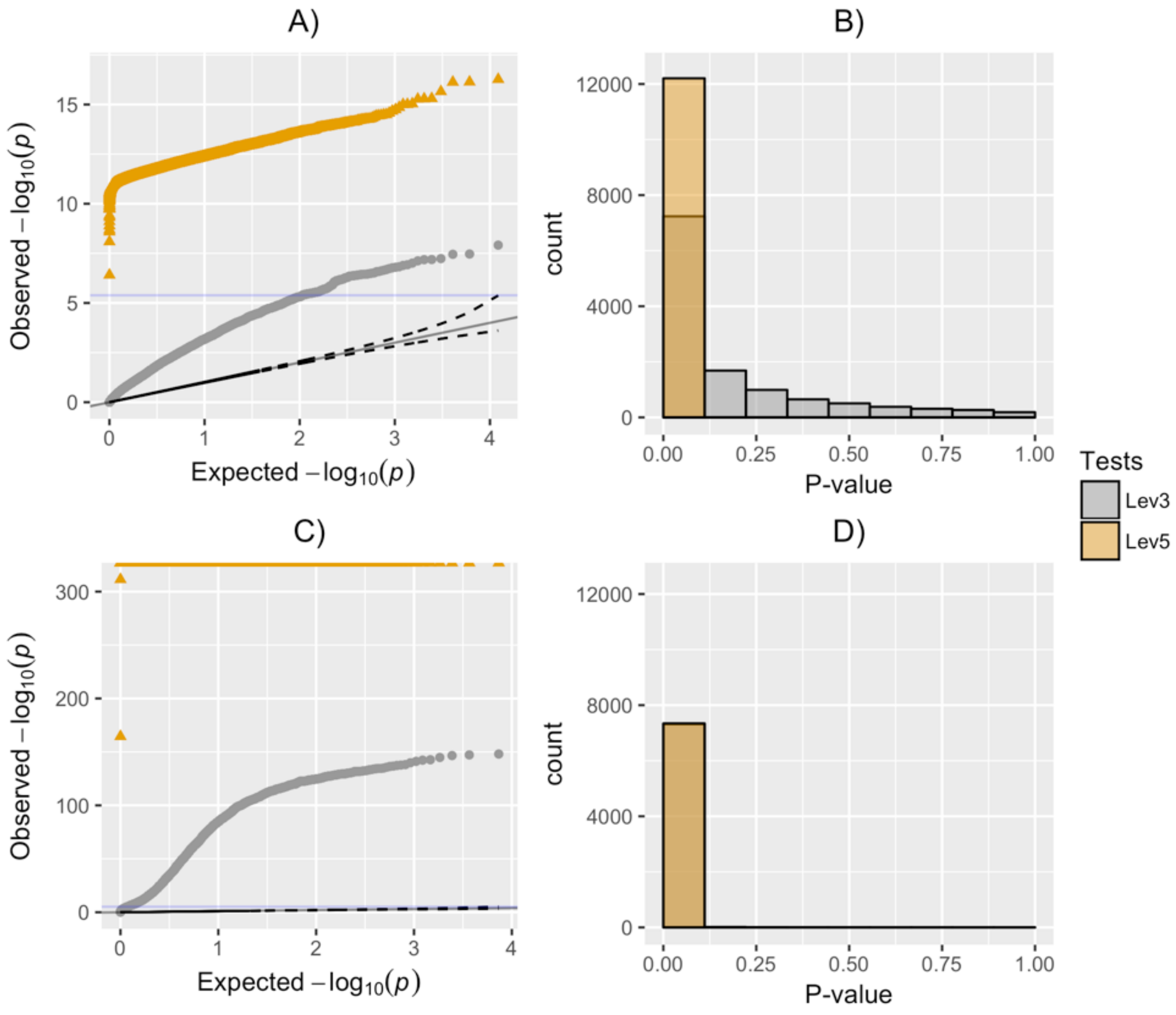

**S4 Figure. Quantile-quantile plots and histograms of variance heterogeneity p-values using Naïve Strategies for body mass index.** For each XCHR SNP, the variance heterogeneity *p*-value was calculated using Levene's test without modifications stratified by either the genotype (Lev3) or the combinations of genotype and sex (Lev5). Quantile-quantile plots (A and C), and histograms (B and D) of the *p*-values were produced using data from the Multi-Ethnic Study of Atherosclerosis are shown on the top row, and on the bottom row for UK Biobank.

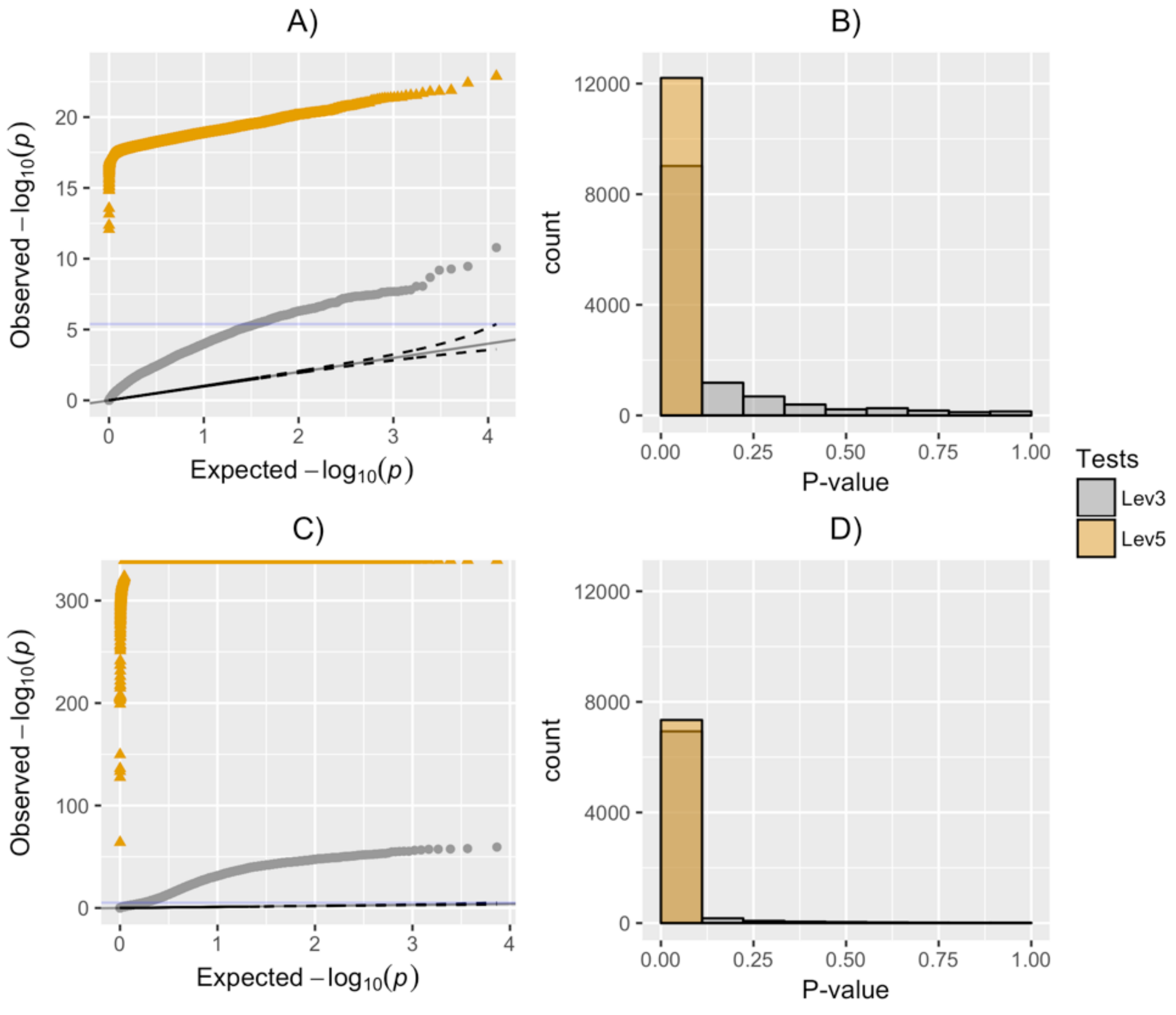

**S5 Figure. Quantile-quantile plots and histograms of variance heterogeneity p-values using Naïve Strategies for waist circumference.** For each XCHR SNP, the variance heterogeneity *p*-value was calculated using Levene's test without modifications stratified by either the genotype (Lev3) or the combinations of genotype and sex (Lev5). Quantile-quantile plots (A and C), and histograms (B and D) of the *p*-values were produced using data from the Multi-Ethnic Study of Atherosclerosis are shown on the top row, and on the bottom row for UK Biobank.

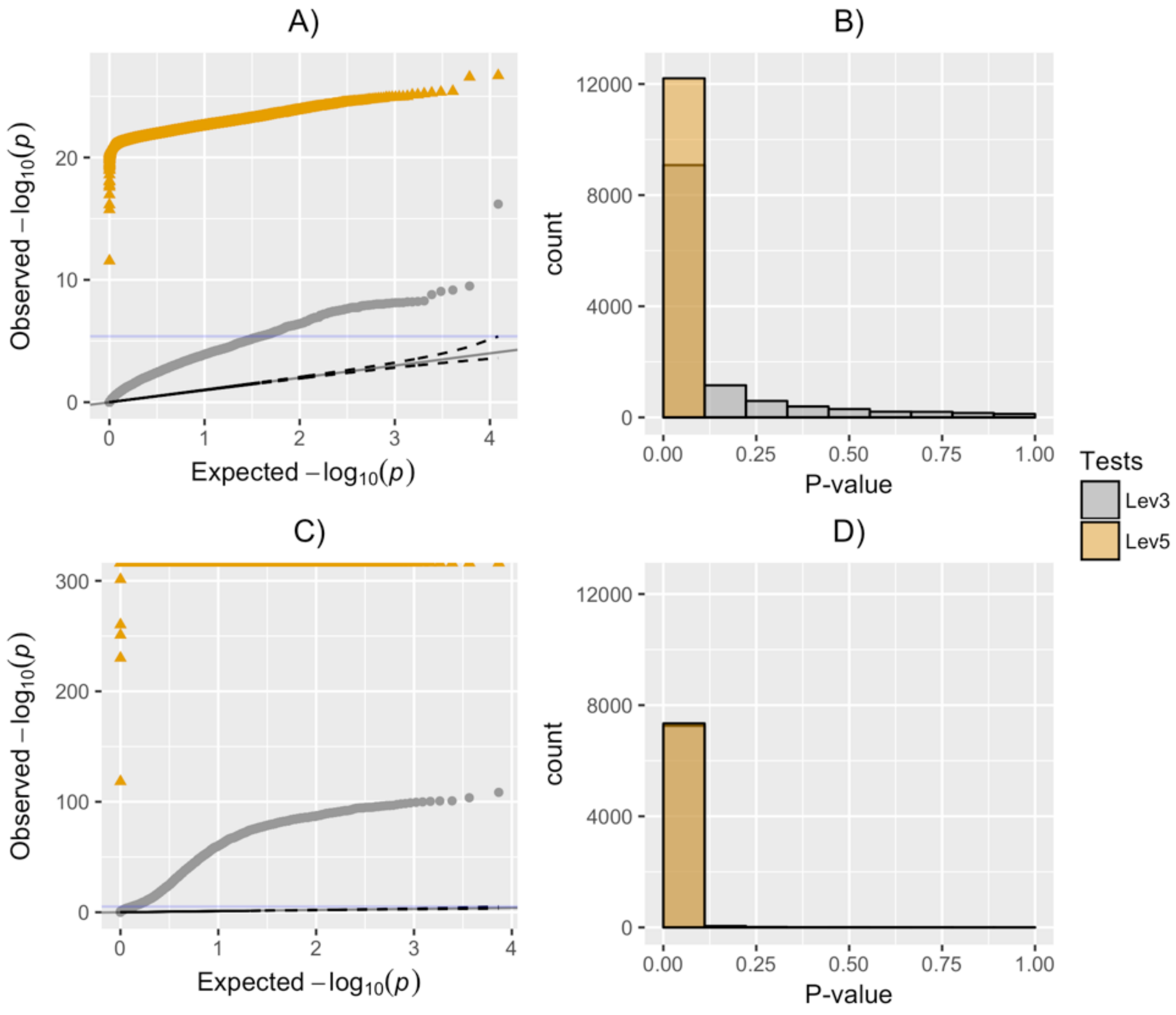

**S6 Figure. Locus zoom plots of rs2681646 variance heterogeneity replication results for waist circumference using MESA data.**

The variance heterogeneity wM3V3.2 *p*-values of nearby SNPs (+/-500KB) for waist circumference in the MESA dataset were plotted.

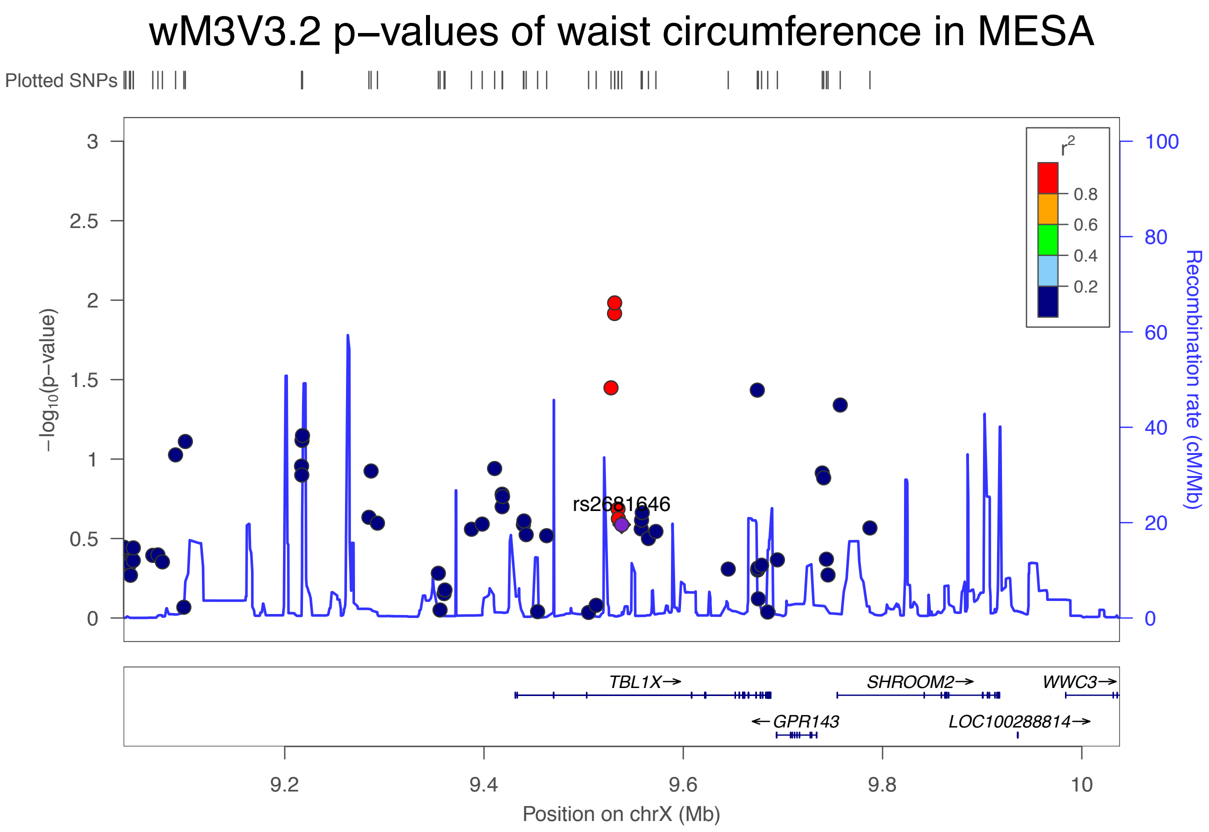

**S7 Figure. XCHR-wide variance heterogeneity test results for height using the UK Biobank (upper panel) and MESA (lower panel) data.**

For each XCHR SNP, the variance heterogeneity *p*-value was calculated using a sex-stratified Fisher's method or the model-based wM3V3.2 and wM3VNA3.3 strategies. Manhattan plots (A and D), quantile-quantile plots (B and E), and histograms of the *p*-values using data from the UK Biobank are shown on the top row, and on the bottom row using data from Multi-Ethnic Study of Atherosclerosis.

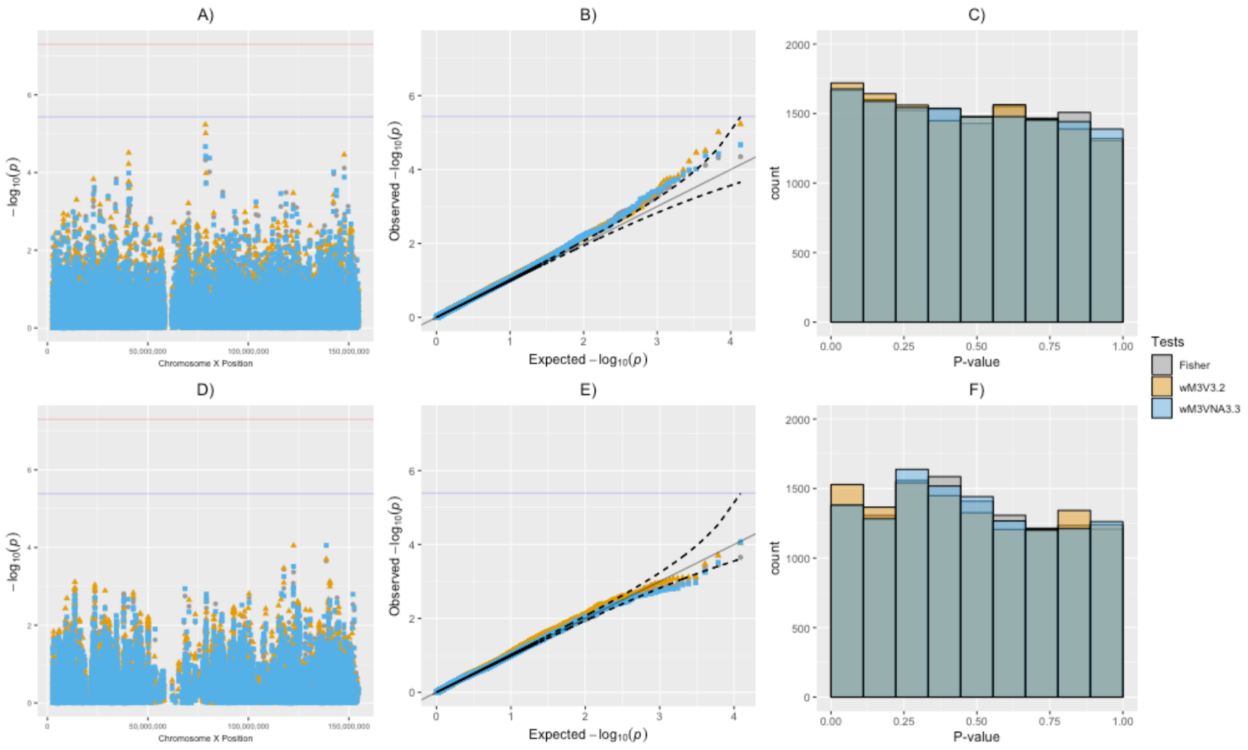

**S8 Figure. XCHR-wide variance heterogeneity test results for hip circumference using the UK Biobank (upper panel) and MESA (lower panel) data.**

For each XCHR SNP, the variance heterogeneity *p*-value was calculated using a sex-stratified Fisher's method or the model-based wM3V3.2 and wM3VNA3.3 strategies. Manhattan plots (A and D), quantile-quantile plots (B and E), and histograms of the *p*-values using data from the UK Biobank are shown on the top row, and on the bottom row using data from Multi-Ethnic Study of Atherosclerosis.

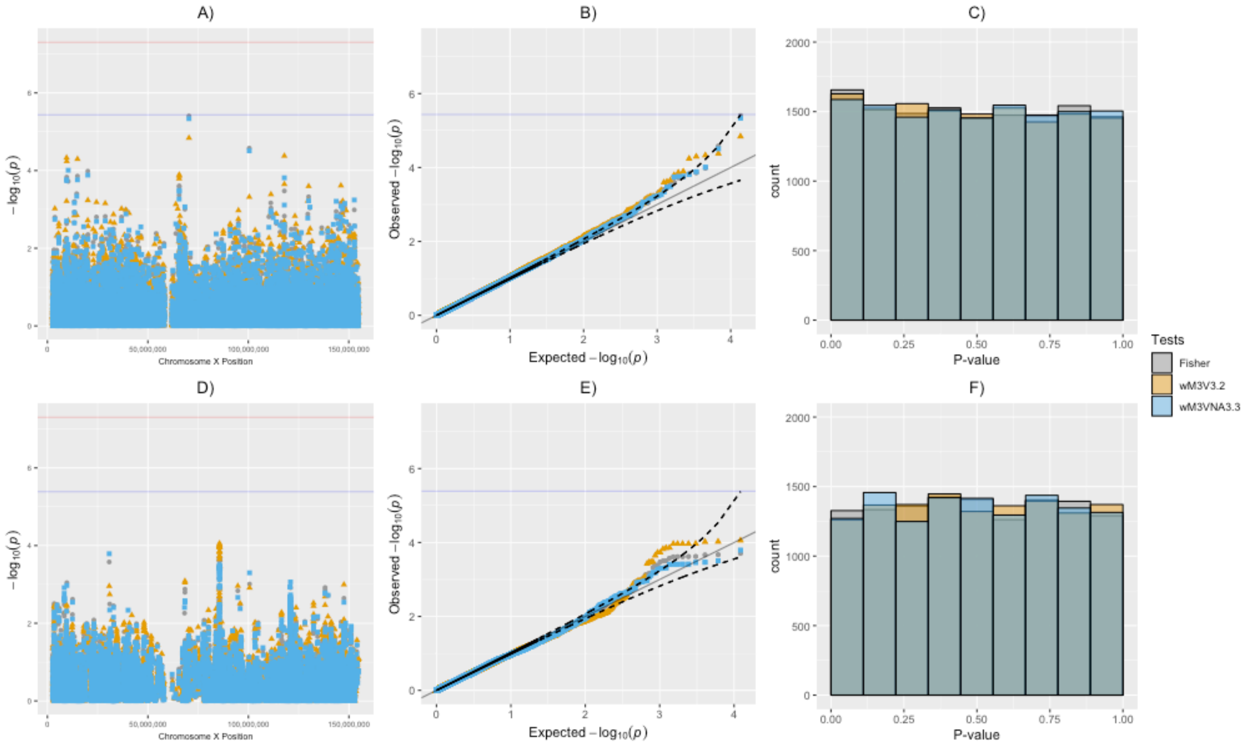

**S9 Figure. XCHR-wide variance heterogeneity test results for body mass index (BMI) using the UK Biobank (upper panel) and MESA (lower panel) data.**

For each XCHR SNP, the variance heterogeneity *p*-value was calculated using a sex-stratified Fisher's method or the model-based wM3V3.2 and wM3VNA3.3 strategies. Manhattan plots (A and D), quantile-quantile plots (B and E), and histograms of the *p*-values using data from the UK Biobank are shown on the top row, and on the bottom row using data from Multi-Ethnic Study of Atherosclerosis.

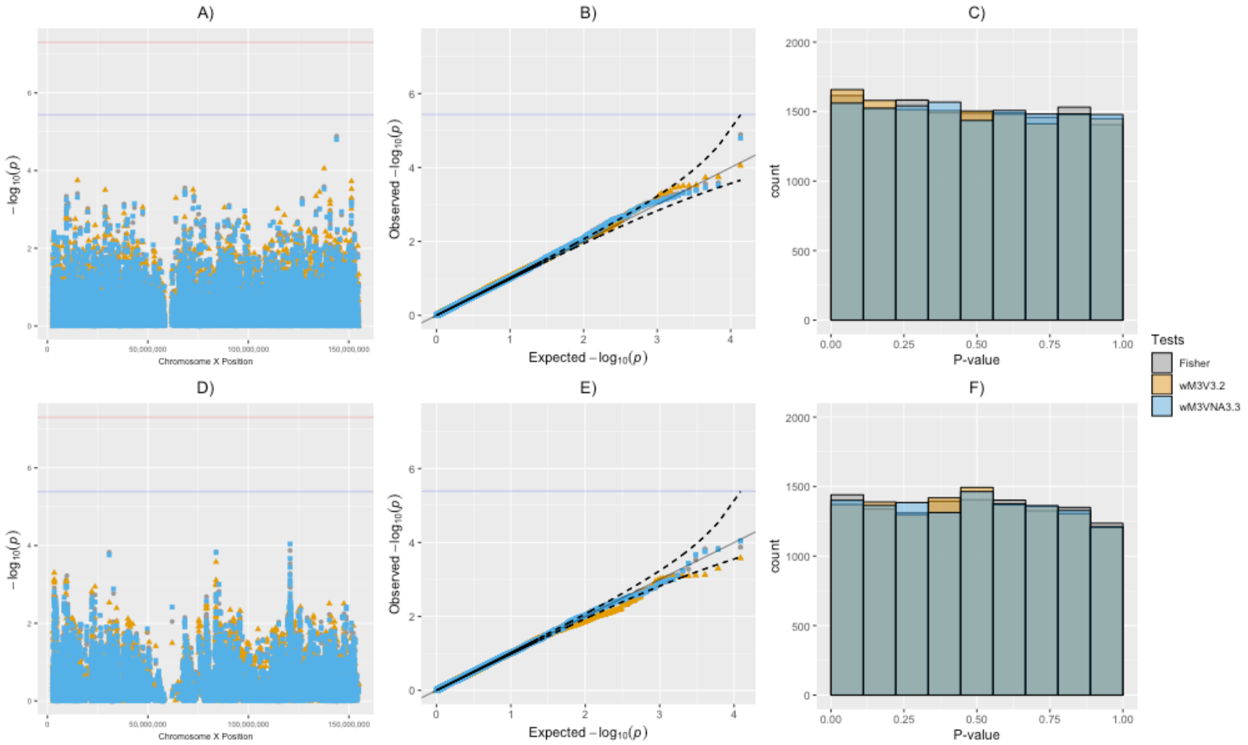

**S10 Figure. A summary of sex-stratified differences in gene expression traits.** For each of the 648 gene expression traits, we computed the sex-stratified mean and variance values, and conducted a *t*-test for equality of means and an *F*-test for equality of variance between the two sexes. The scatterplots of male and female mean and variance values are shown in A) and B), the *p*-values for inequality of mean or variance were summarized in a histogram to show there was an excessive proportion of expression traits with either mean or variance difference between sexes.

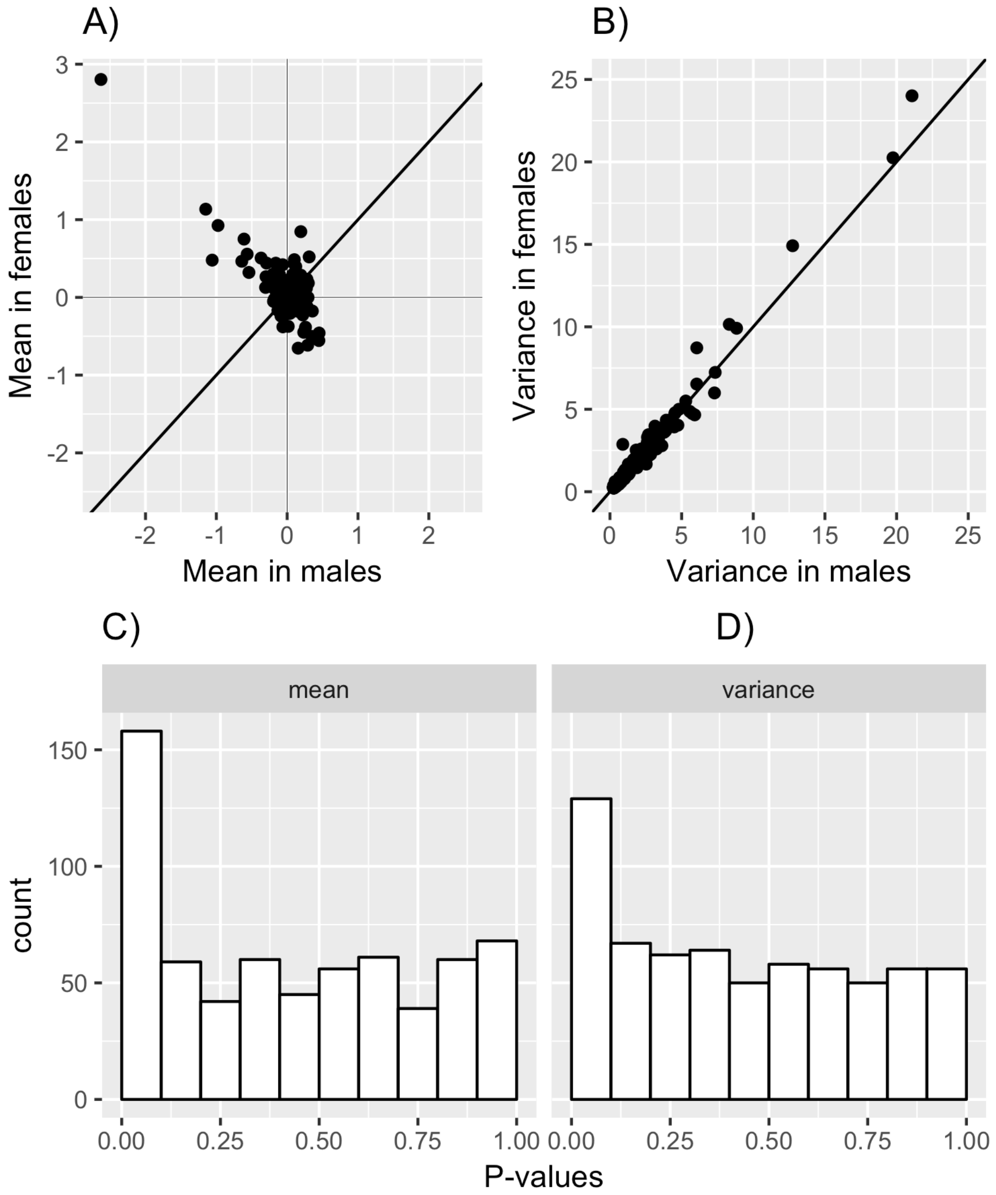

**S11 Figure. XCHR-wide variance heterogeneity test results for gene expression trait at the *NGFRAP1* locus**

For each XCHR SNP, the variance heterogeneity *p*-value was calculated using a sex-stratified Fisher's method or the model-based wM3V3.2 and wM3VNA3.3, and summarized in a Manhattan plot (A), quantile-quantile plot (B), and a histogram (C). SNPs rs222378 and rs3747313 at the *MTMR1* locus was annotated for passing the suggestive significance at 1.3E-05.

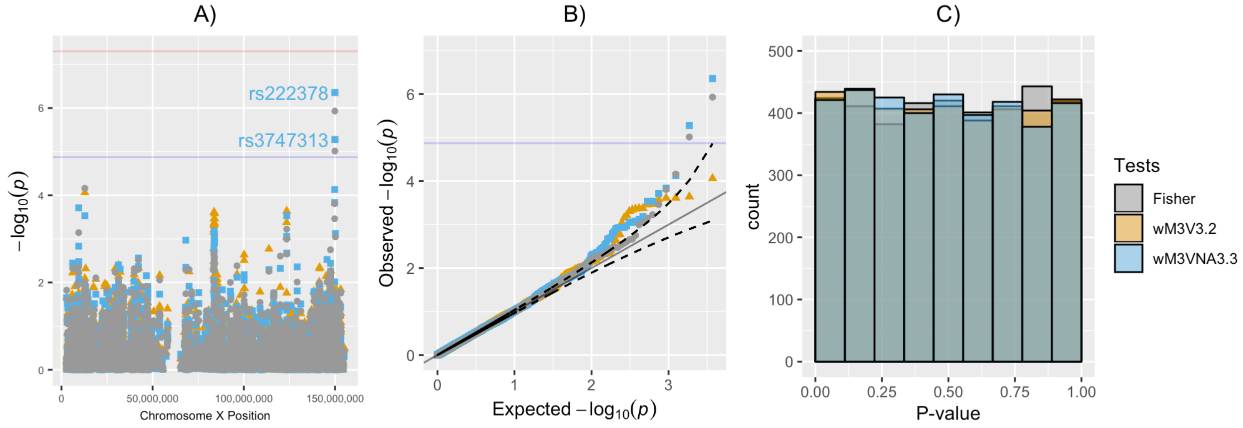

**S12 Figure. XCHR-wide variance heterogeneity test results for gene expression trait at the *ZMYM3* locus**

For each XCHR SNP, the variance heterogeneity *p*-value was calculated using a sex-stratified Fisher's method or the model-based wM3V3.2 and wM3VNA3.3, and summarized in a Manhattan plot (A), quantile-quantile plot (B), and a histogram (C). SNPs rs4830296 and rs4830297 at the *PLAC1* locus was annotated for passing the suggestive significance at 1.3E-05.

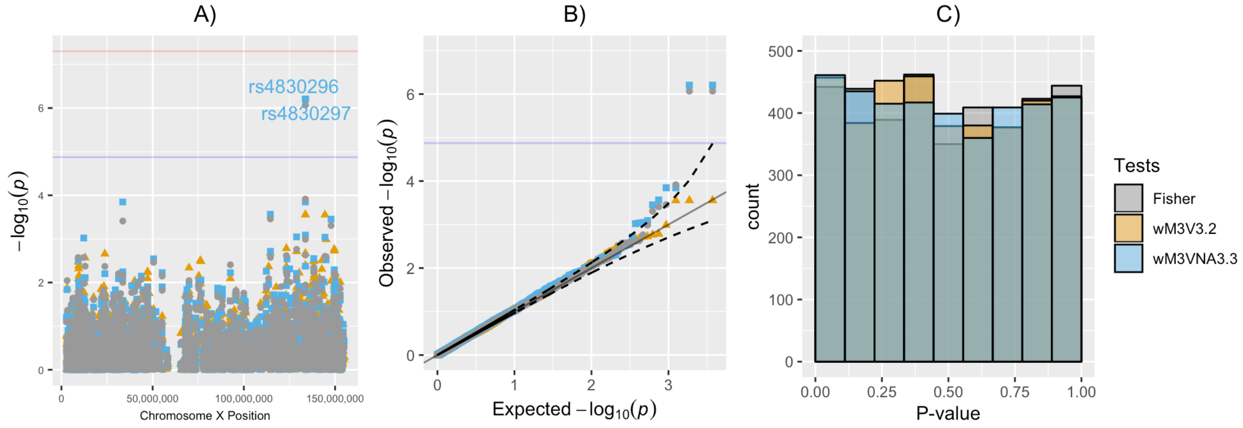

**S13 Figure. XCHR-wide variance heterogeneity test results for gene expression trait at the *TSC22D3* locus**

For each XCHR SNP, the variance heterogeneity $p$-value was calculated using a sex-stratified Fisher's method or the model-based wM3V3.2 and wM3VNA3.3, and summarized in a Manhattan plot (A), quantile-quantile plot (B), and a histogram (C). SNPs rs2040394 at the *PHEX* locus was annotated for passing the suggestive significance at 1.3E-05.

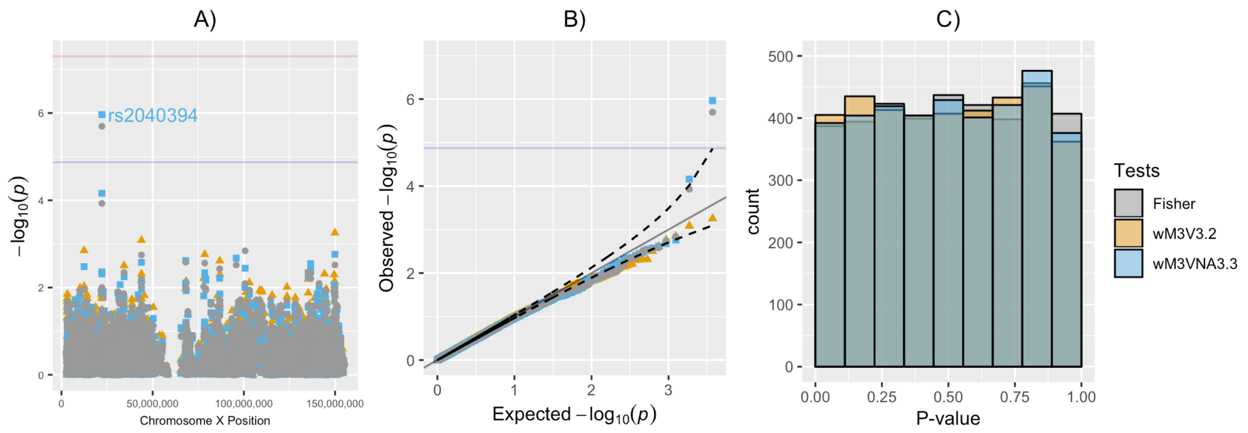

**S14 Figure. Distribution of variance heterogeneity *p*-values over 648 gene expression traits.** A total of 2,412,504 *p*-values of 3,723 SNPs for all 648 gene expression traits were used to generate the overall quantile-quantile plots and histograms for each variance heterogeneity test.

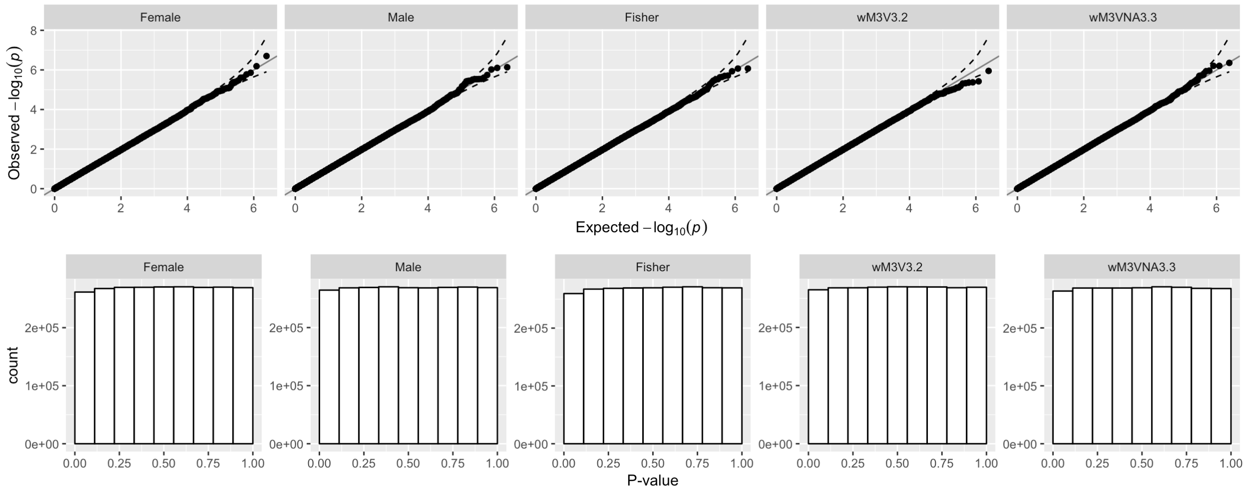

**S15 Figure. Estimated genomic control lambda and proportion of truly associated SNPs among *cis* and *trans* acting variants.** For each gene expression trait, the variance heterogeneity *p*-values from all SNPs that were *cis* were used to produce an estimate of genomic control lambda and the proportion of truly associated SNPs, and are shown as red dots under each test. The remaining *p*-values used to produce estimates of *trans*-acting SNPs and are shown as green dots under each test. We also calculated the estimates using all the *p*-values irrespective of SNPs in *cis* or *trans*, and are shown as blue dots under each test. The distribution of the estimates is shown for each variance heterogeneity testing strategies considered. The black line represents the reference line at $\lambda_{GC} = 1$ or $\pi_1 = 0$.

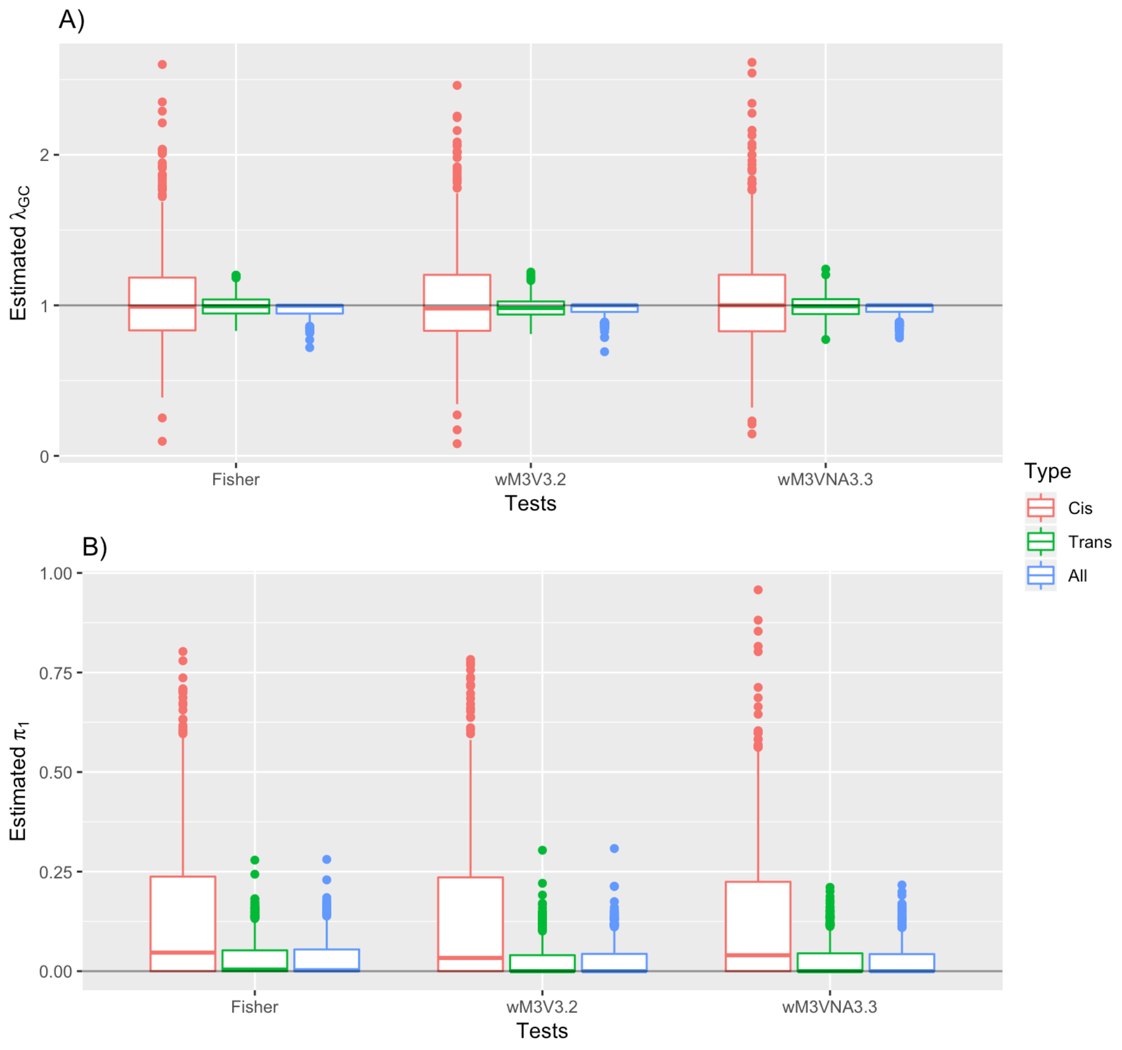

**S1 Table. A summary of variance heterogeneity testing strategies for XCHR SNPs.**

| *Approach* | *Shorthand of strategy* | *Description of the statistical model used* |
|---|---|---|
| *Levene's test (median)* | ***Lev3*** | Levene's test with grouping factor *G* |
| | ***Lev5*** | Levene's test with grouping factor *G*x*S* |
| | ***LevF*** | Levene's test with grouping factor *G* (coded non-additively) in female only |
| | ***LevM*** | Levene's test with grouping factor *G* (= 0, 1) in male only |
| | ***Fisher*** | Fisher's method to combine sex-specific Levene's test *p*-values: ***LevF*** and ***LevM*** |
| *Generalized Levene's test*<br><br>*(Testing for the additive variance effect per reference allele)* | ***wM1V1 –wM3V3*** | Model-based one-degree of freedom test for variance heterogeneity associated with *G* (coded additively by number of reference alleles 0, 1, 2)<br><br>$H_o$: $\gamma_G = 0$ |
| | ***wM1NAV1 –wM3NAV3*** | Model-based one-degree of freedom test for variance heterogeneity associated with *G* (coded additively by number of reference alleles 0, 1, 2)<br><br>$H_o$: $\gamma_{G1} = \gamma_{G2} = 0$ |
| | ***wM1V3.2***<br><br>***wM2V3.2***<br><br>***wM3V3.2*** | Model-based two-degree of freedom test for variance heterogeneity associated with both *G* (coded additively by number of reference alleles 0, 1, 2) and *GS:*<br><br>$H_o$: $\gamma_G = \gamma_{GS} = 0$ |
| | ***wM1VNA3.3***<br><br>***wM2VNA3.3***<br><br>***wM3VNA3.3*** | Model-based three-degree of freedom test for variance heterogeneity associated with non-additive *G* and its interaction with *S:*<br><br>$H_o$: $\gamma_{G1} = \gamma_{G2} = \gamma_{G1S} = 0$ |

There are 5 testing strategies using the original Levene's test, and a total of 24 model-based testing strategies derived from the combinations of mean and variance models using weighted residuals.

#### S2 Table. A summary of sex-stratified mean and variance for quantitative traits.

| | | Untransformed | | | | | | | | Inverse-normally transformed | | | | | | | |
|---|---|---|---|---|---|---|---|---|---|---|---|---|---|---|---|---|---|
| | | Height (cm) | | Hip (cm) | | BMI (kg/m$^2$) | | Waist (cm) | | Height | | Hip | | BMI | | Waist | |
| | | $\mu$ | $\sigma$ | $\mu$ | $\sigma$ | $\mu$ | $\sigma$ | $\mu$ | $\sigma$ | $\mu$ | $\sigma$ | $\mu$ | $\sigma$ | $\mu$ | $\sigma$ | $\mu$ | $\sigma$ |
| **UKB** | Male | 176.02 | 6.74 | 103.24 | 7.27 | 27.61 | 4.05 | 96.43 | 10.86 | 0.76 | 0.71 | 0.05 | 0.85 | 0.13 | 0.87 | 0.52 | 0.78 |
| | Female | 162.74 | 6.20 | 103.08 | 9.99 | 26.85 | 4.97 | 84.12 | 12.05 | -0.64 | 0.72 | -0.04 | 1.11 | -0.11 | 1.08 | -0.43 | 0.96 |
| **MESA** | Male | 176.29 | 6.94 | 104.91 | 8.41 | 27.84 | 4.08 | 100.83 | 11.57 | 0.73 | 0.71 | 0.18 | 0.77 | -0.10 | 0.85 | 0.04 | 0.82 |
| | Female | 162.24 | 6.37 | 106.70 | 11.74 | 27.29 | 5.58 | 94.55 | 15.65 | -0.68 | 0.70 | -0.29 | 1.09 | -0.01 | 1.09 | -0.16 | 1.10 |

Sex-stratified mean ($\mu$) and standard deviation ($\sigma$) for phenotype data from UK Biobank and the Multi-Ethnic Study of Atherosclerosis (MESA). The mean and standard deviation values for the untransformed traits were calculated in their original units, while values for inverse-normally transformed trait were calculated such that the overall distribution has mean 0 and variance 1.

#### S3 Table. Empirical type I error (T1E) rates of X-chromosome variance heterogeneity tests under Simulation Design I.

| | Condition 1 | | Condition 2 | | Condition 3-A | | Condition 3-B | | Condition 4-A | | Condition 4-B | |
|---|---|---|---|---|---|---|---|---|---|---|---|---|
| $\beta_E$ | 0 | 0.5 | 0 | 0.5 | 0 | 0 | 0.5 | 0.5 | 0 | 0 | 0.5 | 0.5 |
| $\beta_S$ | 0 | 0 | 0.5 | 0.5 | 0 | 0 | 0 | 0 | 0.5 | 0.5 | 0.5 | 0.5 |
| $\beta_{SE}$ | 0 | 0 | 0 | 0 | 0.5 | -0.5 | 0.5 | -0.5 | 0.5 | -0.5 | 0.5 | -0.5 |
| **Lev3** | 0.0453 | 0.0445 | **0.0621** | **0.0905** | **0.7747** | **0.8271** | **0.8184** | **0.2584** | **0.8092** | **0.7868** | **0.7489** | **0.1494** |
| **Lev5** | 0.0240 | 0.0235 | 0.0230 | **0.1033** | **1** | **1** | **1** | **1** | **1** | **1** | **1** | **1** |
| **Female** | 0.0484 | 0.0512 | 0.0534 | 0.0508 | 0.0505 | 0.0524 | 0.0495 | 0.0515 | 0.0521 | 0.0464 | 0.0489 | 0.0497 |
| **Male** | 0.0495 | 0.0494 | 0.0514 | 0.0481 | 0.0506 | 0.0471 | 0.0503 | 0.0509 | 0.0507 | 0.0513 | 0.0524 | 0.0485 |
| **Fisher** | 0.0501 | 0.0492 | 0.0521 | 0.0543 | 0.0504 | 0.0503 | 0.0518 | 0.0509 | 0.0501 | 0.0498 | 0.0491 | 0.0483 |
| **wM1V1** | 0.0444 | 0.0449 | **0.0588** | 0.0486 | 0.0450 | 0.0436 | 0.0446 | 0.0478 | 0.0433 | 0.0440 | 0.0461 | 0.0565 |
| **wM1V2** | 0.0486 | 0.0495 | **0.0619** | 0.0563 | 0.0500 | 0.0475 | 0.0501 | 0.0480 | 0.0487 | 0.0488 | 0.0510 | **0.0584** |
| **wM1V3** | 0.0493 | 0.0480 | **0.2198** | **0.1563** | 0.0479 | 0.047 | 0.0481 | 0.0532 | **0.1435** | **0.1458** | **0.1030** | **0.2163** |
| **wM1V3.2** | 0.0481 | 0.0487 | **0.2129** | **0.1432** | 0.0506 | 0.0494 | 0.0505 | 0.0525 | **0.1223** | **0.1251** | **0.0928** | **0.1976** |
| **wM1VNA1** | 0.0458 | 0.0457 | **0.066** | 0.0520 | 0.0472 | 0.0477 | 0.0471 | 0.0497 | 0.0508 | 0.0497 | 0.0481 | **0.0646** |
| **wM1VNA2** | 0.0485 | 0.0482 | **0.068** | **0.0575** | 0.0501 | 0.0503 | 0.0498 | 0.0506 | 0.0563 | 0.0520 | 0.0511 | **0.0659** |
| **wM1VNA3.2** | 0.0489 | 0.0479 | **0.1953** | **0.1344** | 0.0474 | 0.049 | 0.0521 | 0.0501 | **0.1185** | **0.1205** | **0.0924** | **0.1845** |
| **wM1VNA3.3** | 0.0502 | 0.0493 | **0.1846** | **0.1275** | 0.0511 | 0.0512 | 0.0519 | 0.0503 | **0.1066** | **0.1079** | **0.0806** | **0.1678** |
| **wM2V1** | 0.0447 | 0.0446 | 0.0457 | 0.0443 | 0.045 | 0.0434 | 0.0449 | 0.0478 | 0.0427 | 0.0471 | 0.0436 | 0.0455 |
| **wM2V2** | 0.0485 | 0.0496 | 0.0494 | 0.0487 | 0.0499 | 0.0474 | 0.0502 | 0.0483 | 0.0485 | 0.0492 | 0.0476 | 0.0501 |
| **wM2V3** | 0.0486 | 0.0483 | 0.0532 | 0.0481 | 0.0476 | 0.0469 | 0.0483 | 0.0529 | 0.0515 | 0.0505 | 0.0492 | 0.0482 |
| **wM2V3.2** | 0.0488 | 0.0486 | 0.0507 | 0.0504 | 0.0507 | 0.0495 | 0.0509 | 0.0527 | 0.0479 | 0.051 | 0.0481 | 0.0502 |
| **wM2VNA1** | 0.0460 | 0.0453 | 0.0483 | 0.0495 | 0.0473 | 0.0475 | 0.047 | 0.0495 | 0.0444 | 0.0448 | 0.0461 | 0.0517 |

| | | | | | | | | | | | | |
|---|---|---|---|---|---|---|---|---|---|---|---|---|
| **wM2VNA2** | 0.0489 | 0.0478 | 0.0513 | 0.0522 | 0.0501 | 0.0506 | 0.0501 | 0.0505 | 0.0472 | 0.0478 | 0.0502 | 0.0556 |
| **wM2VNA3.2** | 0.0490 | 0.0478 | 0.0533 | 0.0514 | 0.0478 | 0.049 | 0.0519 | 0.0503 | 0.0513 | 0.0501 | 0.0512 | 0.0494 |
| **wM2VNA3.3** | 0.0499 | 0.0493 | 0.0504 | 0.0528 | 0.0509 | 0.0513 | 0.0521 | 0.0503 | 0.0487 | 0.0504 | 0.0485 | 0.049 |
| **wM3V1** | **0.0442** | **0.0444** | 0.0464 | **0.0438** | **0.0445** | **0.0431** | **0.0447** | 0.0472 | **0.0426** | 0.0471 | **0.044** | 0.0459 |
| **wM3V2** | 0.0484 | 0.0498 | 0.0493 | 0.0485 | 0.0500 | 0.0477 | 0.0494 | 0.0478 | 0.0486 | 0.0500 | 0.0472 | 0.0503 |
| **wM3V3** | 0.0488 | 0.0478 | 0.0530 | 0.0478 | 0.0478 | 0.0467 | 0.0484 | 0.0533 | 0.0519 | 0.0502 | 0.0497 | 0.0484 |
| **wM3V3.2** | 0.0480 | 0.0486 | 0.0508 | 0.0510 | 0.0506 | 0.0487 | 0.0505 | 0.0519 | 0.0481 | 0.0509 | 0.0478 | 0.05 |
| **wM3VNA1** | 0.0460 | 0.0451 | 0.048 | 0.0505 | 0.0481 | 0.0477 | 0.0472 | 0.0497 | 0.0445 | 0.045 | 0.0462 | 0.0516 |
| **wM3VNA2** | 0.0490 | 0.0479 | 0.0510 | 0.0531 | 0.0510 | 0.0510 | 0.0494 | 0.0506 | 0.0471 | 0.0467 | 0.0501 | 0.0552 |
| **wM3VNA3.2** | 0.0496 | 0.0483 | 0.0538 | 0.0509 | 0.0489 | 0.0495 | 0.0519 | 0.05 | 0.0508 | 0.0496 | 0.051 | 0.0491 |
| **wM3VNA3.3** | 0.0500 | 0.0499 | 0.0502 | 0.0528 | 0.0516 | 0.0515 | 0.0516 | 0.0499 | 0.0487 | 0.0505 | 0.0488 | 0.0493 |

A quantitative trait was simulated according to simulation design I based on linear regression model (1) with coefficient values specified above such that CONDITION 1 captures the null scenario of no sexual dimorphism, i.e. no sex-specific mean nor variance differences as depicted in Fig 2A; CONDITION 2 corresponds to the conceptual null scenario in Fig 2B with the presence of sex-specific means via a non-zero $\beta_S$; CONDITIONS 3A and 3B correspond to Fig 2C, representing a sex-specific variance difference through a non-zero $\beta_{SE}$, the $S$x$E$ interaction effect, where the environmental effect $\beta_E$ takes a value of either 0 (CONDITION 3A) or 0.5 (CONDITION 3B). Similarly, CONDITIONS 4A and 4B correspond to Fig 2D with sexual dimorphism in both means and variances, with the absence and presence of environmental effect $\beta_E$, respectively. The total sample size was 10,000 with 5,000 females and 5,000 males, and the MAF was 0.2. The nominal T1E rate was set to 0.05 and the empirical T1E rates were calculated based on 10,000 simulated replicates. Those empirical T1E rates exceeding 5%±0.5% were in bold.

**S4 Table. Empirical type I error (T1E) rates of X-chromosome variance heterogeneity tests under Simulation Design II.**

| | *Condition 1* | | *Condition 2-1* | | *Condition 2-2* | | *Condition 2-3* | | *Condition 3* | | *Condition 4-1* | | *Condition 4-2* | | *Condition 4-3* | |
|---|---|---|---|---|---|---|---|---|---|---|---|---|---|---|---|---|
| $\mu_m$ | *-0.01* | *-0.1* | *0.04* | *-0.16* | *0.18* | *-0.29* | *0.73* | *-0.68* | *-0.1* | *-0.01* | *0.04* | *-0.16* | *0.18* | *-0.29* | *0.73* | *-0.68* |
| $\mu_f$ | *-0.1* | *-0.01* | *-0.16* | *0.04* | *-0.29* | *0.18* | *-0.68* | *0.73* | *-0.01* | *-0.1* | *-0.16* | *0.04* | *-0.29* | *0.18* | *-0.68* | *0.73* |
| $\sigma_m^2$ | *0.7* | *0.71* | *0.71* | *0.7* | *0.71* | *0.7* | *0.71* | *0.7* | *0.77* | *1.09* | *0.77* | *1.09* | *0.77* | *1.09* | *0.77* | *1.09* |
| $\sigma_f^2$ | *0.71* | *0.7* | *0.7* | *0.71* | *0.7* | *0.71* | *0.7* | *0.71* | *1.09* | *0.77* | *1.09* | *0.77* | *1.09* | *0.77* | *1.09* | *0.77* |
| **Lev3** | 0.0481 | **0.0776** | 0.0467 | 0.0490 | 0.0495 | 0.0419 | **0.6524** | **0.3848** | **0.6948** | **0.7986** | **0.7452** | **0.6843** | **0.6010** | **0.8071** | **0.2162** | **0.9286** |
| **Lev5** | 0.0538 | **0.0814** | **0.0324** | **0.0276** | **0.0195** | **0.0295** | **0.0276** | **0.0833** | **1.0000** | **1.0000** | **1.0000** | **1.0000** | **1.0000** | **1.0000** | **1.0000** | **1.0000** |
| **Female** | 0.0538 | 0.0519 | 0.0452 | 0.0562 | 0.0510 | 0.0524 | 0.0457 | 0.0452 | 0.0500 | 0.0586 | 0.0386 | 0.0471 | 0.0500 | 0.0424 | 0.0543 | 0.0476 |
| **Male** | 0.0419 | 0.0481 | 0.0600 | 0.0557 | 0.0529 | **0.0357** | 0.0490 | 0.0452 | 0.0500 | 0.0395 | 0.0567 | 0.0462 | 0.0495 | 0.0467 | 0.0590 | 0.0448 |
| **Fisher** | 0.0481 | 0.0519 | 0.0557 | 0.0562 | 0.0505 | 0.0390 | 0.0505 | 0.0500 | 0.0443 | 0.0571 | 0.0514 | 0.0452 | 0.0505 | 0.0414 | 0.0505 | 0.0476 |
| **wM1V1** | 0.0381 | 0.0405 | 0.0400 | 0.0490 | 0.0462 | 0.0500 | **0.5919** | **0.3829** | 0.0414 | 0.0433 | 0.0395 | 0.0371 | 0.0462 | 0.0486 | 0.0438 | **0.4786** |
| **wM1V2** | 0.0457 | 0.0457 | 0.0529 | 0.0571 | **0.0776** | **0.0762** | **0.8024** | **0.6162** | 0.0500 | 0.0524 | 0.0443 | 0.0448 | 0.0586 | 0.0600 | **0.0800** | **0.6381** |
| **wM1V3** | 0.0500 | 0.0490 | 0.0638 | **0.0738** | **0.2762** | **0.2824** | **0.9995** | **0.9990** | 0.0567 | 0.0405 | 0.0595 | 0.0481 | **0.2348** | **0.0933** | **0.9938** | **0.9814** |
| **wM1V3.2** | 0.0490 | 0.0500 | 0.0638 | **0.0671** | **0.2752** | **0.2910** | **0.9990** | **0.9986** | 0.0495 | 0.0557 | 0.0500 | 0.0529 | **0.2062** | **0.0938** | **0.9924** | **0.9933** |

| | | | | | | | | | | | | | | | | |
|---|---|---|---|---|---|---|---|---|---|---|---|---|---|---|---|---|
| **wM1VNA1** | 0.0386 | 0.0395 | 0.0471 | 0.0495 | 0.0567 | 0.0567 | **0.9257** | **0.8562** | 0.0405 | 0.0429 | 0.0443 | 0.0424 | 0.0533 | 0.0424 | **0.5795** | **0.5995** |
| **wM1VNA2** | 0.0443 | 0.0462 | 0.0557 | 0.0562 | **0.0848** | **0.0771** | **0.9629** | **0.9148** | 0.0505 | 0.0529 | 0.0481 | 0.0457 | **0.0719** | 0.0514 | **0.6319** | **0.7243** |
| **wM1VNA3.2** | 0.0486 | 0.0533 | **0.0652** | **0.0686** | **0.2724** | **0.2781** | **0.9995** | **0.9990** | 0.0505 | 0.0490 | 0.0433 | 0.0533 | **0.1995** | **0.0900** | **0.9924** | **0.9895** |
| **wM1VNA3.3** | 0.0505 | 0.0505 | **0.0643** | **0.0667** | **0.2419** | **0.2433** | **0.9990** | **0.9986** | 0.0486 | 0.0567 | 0.0490 | 0.0443 | **0.1733** | **0.0781** | **0.9924** | **0.9895** |
| **wM2V1** | 0.0371 | 0.0443 | 0.0429 | 0.0457 | 0.0471 | **0.0352** | 0.0467 | 0.0448 | 0.0405 | 0.0419 | 0.0424 | 0.0371 | 0.0414 | 0.0376 | 0.0405 | 0.0467 |
| **wM2V2** | 0.0438 | 0.0500 | 0.0500 | 0.0557 | 0.0562 | 0.0438 | 0.0524 | 0.0562 | 0.0505 | 0.0524 | 0.0481 | 0.0448 | 0.0448 | 0.0457 | 0.0452 | 0.0571 |
| **wM2V3** | 0.0519 | 0.0538 | 0.0595 | 0.0600 | 0.0543 | 0.0395 | 0.0429 | 0.0500 | 0.0548 | 0.0405 | 0.0610 | 0.0476 | 0.0538 | 0.0486 | 0.0538 | 0.0438 |
| **wM2V3.2** | 0.0476 | 0.0486 | 0.0548 | 0.0581 | 0.0500 | 0.0414 | 0.0457 | 0.0452 | 0.0476 | 0.0548 | 0.0505 | 0.0457 | 0.0514 | 0.0462 | 0.0471 | 0.0443 |
| **wM2VNA1** | 0.0400 | 0.0433 | 0.0467 | 0.0457 | 0.0448 | 0.0390 | 0.0486 | 0.0452 | 0.0386 | 0.0414 | 0.0429 | 0.0438 | 0.0419 | 0.0386 | 0.0457 | 0.0443 |
| **wM2VNA2** | 0.0471 | 0.0500 | 0.0548 | 0.0529 | 0.0505 | 0.0433 | 0.0505 | 0.0510 | 0.0471 | 0.0524 | 0.0490 | 0.0476 | 0.0471 | 0.0424 | 0.0533 | 0.0476 |
| **wM2VNA3.2** | 0.0481 | 0.0524 | 0.0600 | **0.0662** | 0.0533 | 0.0438 | 0.0467 | 0.0500 | 0.0486 | 0.0500 | 0.0462 | 0.0510 | 0.0467 | 0.0424 | 0.0571 | 0.0433 |
| **wM2VNA3.3** | 0.0471 | 0.0519 | 0.0538 | 0.0590 | 0.0495 | 0.0424 | 0.0476 | 0.0490 | 0.0505 | 0.0590 | 0.0481 | 0.0467 | 0.0505 | 0.0433 | 0.0510 | 0.0495 |
| **wM3V1** | 0.0367 | 0.0457 | 0.0462 | 0.0467 | 0.0467 | **0.0348** | 0.0457 | 0.0443 | 0.0395 | 0.0419 | 0.0414 | 0.0371 | 0.0410 | 0.0390 | 0.0400 | 0.0476 |
| **wM3V2** | 0.0457 | 0.0500 | 0.0533 | 0.0562 | 0.0548 | 0.0438 | 0.0529 | 0.0562 | 0.0481 | 0.0524 | 0.0481 | 0.0429 | 0.0452 | 0.0462 | 0.0433 | 0.0557 |
| **wM3V3** | 0.0538 | 0.0519 | 0.0576 | 0.0581 | 0.0548 | 0.0371 | 0.0405 | 0.0471 | 0.0543 | 0.0410 | 0.0595 | 0.0490 | 0.0548 | 0.0471 | 0.0533 | 0.0429 |
| **wM3V3.2** | 0.0471 | 0.0490 | 0.0543 | 0.0581 | 0.0505 | 0.0410 | 0.0462 | 0.0452 | 0.0462 | 0.0543 | 0.0490 | 0.0457 | 0.0524 | 0.0471 | 0.0462 | 0.0438 |
| **wM3VNA1** | 0.0405 | 0.0419 | 0.0481 | 0.0443 | 0.0429 | 0.0386 | 0.0476 | 0.0452 | 0.0381 | 0.0405 | 0.0433 | 0.0438 | 0.0429 | 0.0386 | 0.0457 | 0.0438 |
| **wM3VNA2** | 0.0467 | 0.0490 | 0.0571 | 0.0519 | 0.0510 | 0.0429 | 0.0500 | 0.0505 | 0.0443 | 0.0505 | 0.0495 | 0.0476 | 0.0467 | 0.0433 | 0.0533 | 0.0476 |
| **wM3VNA3.2** | 0.0481 | 0.0533 | 0.0605 | 0.0610 | 0.0533 | 0.0457 | 0.0462 | 0.0495 | 0.0500 | 0.0495 | 0.0452 | 0.0505 | 0.0467 | 0.0457 | 0.0576 | 0.0429 |
| **wM3VNA3.3** | 0.0486 | 0.0524 | 0.0529 | 0.0571 | 0.0505 | 0.0433 | 0.0495 | 0.0481 | 0.0476 | 0.0586 | 0.0471 | 0.0462 | 0.0476 | 0.0424 | 0.0505 | 0.0476 |

A quantitative trait was simulated according to simulation design II based on sex-stratified distributions with mean and variance values specified above to mimic the observed sex-stratified distributions of height and weight that best matches the Conditions introduced in Fig 1. Here Condition 1 captures the null scenario of no sexual dimorphism, i.e. no sex-specific mean nor variance differences as depicted in Fig 2A; Condition 2 corresponds to the conceptual null scenario in Fig 2B with the presence of sex-specific means with varying levels of differences; Condition 3 corresponds to Fig 2C, representing a sex-specific variance difference. Similarly, Condition 4 corresponds to Fig 2D with sexual dimorphism in both means and variances while allowing a varying degree of differences in specific means. The testing is based on real genotypes from the MESA data, with a sample of 1,003 females and 1,070 males. The genotype data of 12,206 SNPs in MESA were first LD pruned based on a window size of 50, a step size of 10 and a variance inflation factor of 3 among females using PLINK and then filtered by a minimum count of 30 observations in the five sex-genotype stratified groups ($n$ = 2,073, $m$ = 2,100). The nominal T1E rate was set to 0.05 and the empirical T1E rates were calculated based on 2,100 replicates. Those empirical T1E rates exceeding 5%±1.5% (3$SD$ = $3(0.05*0.95/2{,}100)^{0.5}$= 0.015) were in bold. Testing strategies that showed satisfactory T1E controls were underlined.

#### S5 Table. Empirical type I error (T1E) rates of X-chromosome variance heterogeneity tests under Simulation Design I with unknown X-inactivation.

| | Ignoring X-inactivation | | | | | | Model X-inactivation | | | | | |
|---|---|---|---|---|---|---|---|---|---|---|---|---|
| | Condition 1 | Condition 2 | Condition 3 | | Condition 4 | | Condition 1 | Condition 2 | Condition 3 | | Condition 4 | |
| $\beta_S$ | 0 | 0.5 | 0 | 0 | 0.5 | 0.5 | 0 | 0.5 | 0 | 0 | 0.5 | 0.5 |
| $\beta_{SE}$ | 0 | 0 | 0.1 | -0.1 | 0.1 | -0.1 | 0 | 0 | 0.1 | -0.1 | 0.1 | -0.1 |
| Female | 0.048 | 0.046 | 0.040 | 0.048 | 0.044 | 0.044 | 0.048 | 0.046 | 0.040 | 0.048 | 0.044 | 0.044 |
| Male | 0.053 | 0.055 | 0.060 | 0.049 | 0.057 | 0.053 | 0.053 | 0.055 | 0.060 | 0.049 | 0.057 | 0.053 |
| Fisher | 0.048 | 0.051 | 0.046 | 0.053 | 0.051 | 0.045 | 0.048 | 0.051 | 0.046 | 0.053 | 0.051 | 0.045 |
| wM1V1 | **1.000** | **1.000** | **1.000** | **1.000** | **1.000** | **1.000** | 0.036 | 0.037 | 0.033 | 0.043 | 0.032 | 0.040 |
| wM1V2 | **1.000** | **1.000** | **1.000** | **1.000** | **1.000** | **1.000** | 0.039 | 0.039 | 0.036 | 0.045 | 0.035 | 0.041 |
| wM1V3 | **1.000** | **1.000** | **1.000** | **1.000** | **1.000** | **1.000** | 0.047 | 0.051 | 0.051 | 0.042 | 0.052 | 0.046 |
| wM1V3.2 | **1.000** | **1.000** | **1.000** | **1.000** | **1.000** | **1.000** | 0.051 | 0.057 | 0.051 | 0.052 | 0.052 | 0.051 |
| wM1VNA1 | **1.000** | **1.000** | **1.000** | **1.000** | **1.000** | **1.000** | **1.000** | **1.000** | **1.000** | **1.000** | **1.000** | **1.000** |
| wM1VNA2 | **1.000** | **1.000** | **1.000** | **1.000** | **1.000** | **1.000** | 0.059 | 0.060 | 0.050 | 0.055 | 0.050 | 0.050 |
| wM1VNA3.2 | **1.000** | **1.000** | **1.000** | **1.000** | **1.000** | **1.000** | **0.067** | **0.068** | **0.065** | **0.065** | **0.065** | **0.065** |
| wM1VNA3.3 | **1.000** | **1.000** | **1.000** | **1.000** | **1.000** | **1.000** | 0.061 | **0.064** | 0.058 | 0.060 | 0.058 | 0.059 |
| wM2V1 | **1.000** | **1.000** | **1.000** | **1.000** | **1.000** | **1.000** | 0.036 | 0.039 | 0.034 | 0.042 | 0.034 | 0.041 |
| wM2V2 | **1.000** | **1.000** | **1.000** | **1.000** | **1.000** | **1.000** | 0.039 | 0.041 | 0.036 | 0.045 | 0.035 | 0.043 |
| wM2V3 | **1.000** | **1.000** | **1.000** | **1.000** | **1.000** | **1.000** | 0.047 | 0.047 | 0.051 | 0.042 | 0.051 | 0.040 |
| wM2V3.2 | **1.000** | **1.000** | **1.000** | **1.000** | **1.000** | **1.000** | 0.051 | 0.055 | 0.051 | 0.052 | 0.050 | 0.049 |
| wM2VNA1 | **1.000** | **1.000** | **1.000** | **1.000** | **1.000** | **1.000** | **1.000** | **1.000** | **1.000** | **1.000** | **1.000** | **1.000** |
| wM2VNA2 | **1.000** | **1.000** | **1.000** | **1.000** | **1.000** | **1.000** | 0.058 | 0.059 | 0.051 | 0.056 | 0.048 | 0.050 |
| wM2VNA3.2 | **1.000** | **1.000** | **1.000** | **1.000** | **1.000** | **1.000** | **0.067** | **0.071** | **0.065** | **0.065** | **0.065** | 0.061 |
| wM2VNA3.3 | **1.000** | **1.000** | **1.000** | **1.000** | **1.000** | **1.000** | 0.062 | 0.062 | 0.058 | 0.059 | 0.062 | 0.058 |
| wM3V1 | **0.962** | **0.961** | **0.984** | **0.943** | **0.980** | **0.944** | 0.036 | 0.040 | 0.034 | 0.041 | 0.034 | 0.041 |
| wM3V2 | 0.056 | 0.056 | 0.048 | 0.056 | 0.051 | 0.058 | 0.039 | 0.041 | 0.036 | 0.045 | 0.035 | 0.042 |
| wM3V3 | 0.038 | 0.038 | 0.043 | 0.035 | 0.041 | 0.035 | 0.047 | 0.048 | 0.051 | 0.041 | 0.051 | 0.040 |
| wM3V3.2 | 0.051 | 0.055 | 0.051 | 0.053 | 0.051 | 0.049 | 0.051 | 0.055 | 0.051 | 0.053 | 0.051 | 0.049 |
| wM3VNA1 | **0.936** | **0.935** | **0.964** | **0.914** | **0.963** | **0.917** | **1.000** | **1.000** | **1.000** | **1.000** | **1.000** | **1.000** |
| wM3VNA2 | **0.064** | **0.064** | 0.055 | **0.067** | **0.064** | **0.065** | 0.058 | 0.059 | 0.051 | 0.055 | 0.048 | 0.050 |
| wM3VNA3.2 | 0.057 | 0.056 | 0.050 | 0.055 | 0.059 | 0.049 | **0.068** | **0.071** | **0.064** | **0.065** | **0.066** | 0.060 |
| wM3VNA3.3 | 0.062 | 0.062 | 0.058 | 0.060 | 0.062 | 0.057 | 0.062 | 0.062 | 0.058 | 0.060 | 0.062 | 0.057 |

A quantitative trait was simulated according to simulation design I based on linear regression model (1) assuming X-inactivation coding with coding of 0, 1 and 2 for the *bb*, *Bb* and *BB* genotypes in females and 0 and 2 for the *b* and *B* genotypes in males, the *G=1* group contains only females. The environmental effect $\beta_E$ takes a value of 0.5 and the remaining effects involving gene-environment interactions $\beta_{GS} = \beta_{GE} = \beta_{GSE} = 0$. To create a large variance effect through X-inactivation, we assume the genetic main effect $\beta_E = 2$. We use coefficient values specified above such that CONDITION 1 captures the null scenario of no sexual dimorphism, i.e. no sex-specific mean nor variance differences as depicted in Fig 2A; CONDITION 2 corresponds to the conceptual null scenario in Fig 2B with the presence of sex-specific means via a non-zero $\beta_S$; CONDITION 3 corresponds to Fig 2C, representing a sex-specific variance difference through a non-zero $\beta_{SE}$, the *S*x*E* interaction effect. Similarly, CONDITION 4 corresponds to Fig 2D with sexual dimorphism in both means and variances through a non-zero $\beta_S$ and $\beta_{SE}$. The total sample size was 20,000 with 10,000 females and 10,000 males, and the MAF was 0.2. The nominal T1E rate was set to 0.05 and the empirical T1E rates were calculated based on 5,000 simulated replicates. Those empirical T1E rates exceeding 5%±1.5% were in bold.

#### S6 Table. SNPs with the top 5 lowest *p*-values by wM3V3.2, wM3V3.3, or Fisher's for variance heterogeneity using data from UK Biobank.

| *rait* | *SNP* | *Position** | *Gene* | *Variance heterogeneity test p-value* | | | | | *Sample size* | | | | |
|---|---|---|---|---|---|---|---|---|---|---|---|---|---|
| | | | | *Female* | *Male* | *Fisher* | *wM3V3.2* | *wM3VNA3.3* | *Female G/G* | *Female G/g* | *Female g/g* | *Male G* | *Male g* |
| | rs5912970 | 78798170 | | 6.11E-03 | 2.10E-04 | 1.87E-05 | 1.88E-06 | 2.98E-06 | 84777 | 74699 | 16323 | 102162 | 45828 |
| | rs1474563 | 78649193 | | 1.41E-02 | 2.51E-04 | 4.79E-05 | 5.87E-06 | 2.14E-05 | 59597 | 87367 | 31556 | 85996 | 62518 |
| | rs5912299 | 78790814 | | 4.29E-03 | 7.72E-04 | 4.52E-05 | 9.78E-06 | 3.77E-05 | 85177 | 75335 | 16829 | 102144 | 45596 |
| **eight** | rs5918011 | 40473966 | | 3.24E-05 | 5.37E-01 | 2.08E-04 | 3.09E-05 | 1.03E-04 | 146610 | 30153 | 1488 | 134668 | 13723 |
| | rs241127 | 147762753 | *AFF2* | 9.18E-01 | 6.38E-06 | 7.64E-05 | 3.49E-05 | 1.31E-04 | 111158 | 59343 | 7818 | 116740 | 31688 |
| | rs144487202 | 142505526 | | 3.69E-03 | 2.34E-03 | 1.09E-04 | 3.74E-04 | 9.63E-05 | 167291 | 11199 | 185 | 143782 | 4764 |
| | rs147258056 | 80715033 | | 1.43E-03 | 5.26E-03 | 9.60E-05 | 1.91E-02 | 4.22E-05 | 169777 | 8777 | 88 | 144576 | 3764 |
| | rs1803001 | 70459753 | *ZMYM3* | 1.32E-02 | 1.83E-05 | 3.92E-06 | 1.45E-05 | 4.61E-06 | 156703 | 21195 | 734 | 139105 | 9371 |
| | rs2430200 | 117941116 | | 2.97E-05 | 9.85E-01 | 3.35E-04 | 4.19E-05 | 1.54E-04 | 86833 | 75311 | 16183 | 103925 | 44482 |
| | rs2681646 | 9538206 | ***TBL1X*** | 1.12E-04 | 1.34E-01 | 1.81E-04 | 4.68E-05 | 1.74E-04 | 73268 | 82200 | 22695 | 95922 | 52461 |
| **MI** | rs73441359 | 14958821 | | 1.15E-02 | 9.71E-04 | 1.39E-04 | 5.05E-05 | 1.73E-04 | 116864 | 54805 | 6269 | 120220 | 28147 |
| | rs73188221 | 9613728 | ***TBL1X*** | 1.59E-03 | 7.50E-03 | 1.47E-04 | 5.76E-05 | 1.81E-04 | 171442 | 7143 | 61 | 145370 | 3092 |
| | rs145211237 | 100484902 | *DRP2* | 2.53E-04 | 7.37E-03 | 2.64E-05 | 4.79E-04 | 3.10E-05 | 112082 | 58762 | 7691 | 117597 | 30786 |
| | rs60849679 | 20116215 | *MAP7D2* | 9.27E-03 | 8.80E-04 | 1.04E-04 | 5.81E-04 | 1.31E-04 | 120387 | 52269 | 5599 | 121993 | 26320 |
| | rs5934902 | 10360284 | | 1.76E-05 | 8.91E-01 | 1.89E-04 | 1.95E-03 | 9.84E-05 | 145731 | 30978 | 1737 | 134189 | 14359 |
| | rs17284572 | 137714384 | *FGF13* | 2.10E-03 | 1.05E-02 | 2.59E-04 | 8.86E-05 | 3.00E-04 | 159917 | 18156 | 458 | 140526 | 8009 |
| | rs73441359 | 14958821 | | 2.07E-02 | 1.67E-03 | 3.90E-04 | 1.79E-04 | 4.88E-04 | 116864 | 54805 | 6269 | 120220 | 28147 |
| | rs145701879 | 151337092 | *GABRA3* | 3.78E-01 | 1.29E-04 | 5.31E-04 | 1.88E-04 | 6.53E-04 | 153874 | 22736 | 766 | 137999 | 10032 |
| | rs62610383 | 151379479 | *GABRA3* | 1.01E-01 | 5.89E-04 | 6.38E-04 | 2.99E-04 | 8.00E-04 | 144672 | 31501 | 1728 | 133828 | 14429 |
| **ip** | rs239875 | 72510820 | | 6.02E-04 | 1.57E-01 | 9.72E-04 | 3.12E-04 | 9.39E-04 | 152374 | 24702 | 1046 | 137193 | 11013 |
| | rs5980801 | 68444238 | | 7.64E-02 | 3.14E-04 | 2.79E-04 | 3.34E-04 | 3.46E-04 | 128829 | 45594 | 4023 | 126242 | 22153 |
| | rs1088599 | 72963950 | | 1.43E-04 | 3.95E-01 | 6.08E-04 | 4.37E-04 | 4.83E-04 | 157655 | 19896 | 641 | 139761 | 8732 |
| | rs191398726 | 143928888 | | 1.07E-05 | 8.10E-02 | 1.30E-05 | 4.48E-02 | 1.59E-05 | 152449 | 24969 | 1010 | 136118 | 11375 |
| | rs150558194 | 68442084 | | 3.64E-04 | 9.78E-02 | 4.00E-04 | 1.35E-01 | 4.13E-04 | 121793 | 51282 | 5290 | 122354 | 25943 |
| | rs2681646 | 9538206 | ***TBL1X*** | 4.70E-05 | 5.24E-03 | 3.99E-06 | 9.45E-07 | 4.11E-06 | 73268 | 82200 | 22695 | 95922 | 52461 |
| | rs73528831 | 68334031 | | 8.81E-04 | 2.51E-03 | 3.10E-05 | 1.22E-05 | 4.79E-05 | 171286 | 7327 | 66 | 145210 | 3160 |
| **aist** | rs1795588 | 32190697 | *DMD* | 1.14E-01 | 7.57E-05 | 1.09E-04 | 6.24E-05 | 1.97E-04 | 70524 | 83432 | 3229 | 128049 | 19815 |
| | rs2521413 | 9531042 | ***TBL1X*** | 1.59E-03 | 1.32E-02 | 2.46E-04 | 6.98E-05 | 2.56E-04 | 64232 | 85631 | 28426 | 89759 | 58595 |
| | rs10521946 | 28770685 | *IL1RAPL1* | 2.15E-01 | 8.11E-05 | 2.09E-04 | 1.84E-04 | 4.04E-04 | 113734 | 57608 | 6474 | 120096 | 28193 |
| | rs146858325 | 96825471 | *DIAPH2* | 1.91E-05 | 3.01E-01 | 7.52E-05 | 2.83E-04 | 4.03E-05 | 152496 | 24481 | 24458 | 93708 | 54751 |

* Position in base pair is based on genomic placement according to the Genome Reference Consortium Human Build 38 patch release 7.

#### S7 Table. *P*-values (< 0.05) by wM3V3.2, wM3V3.3, or Fisher's for variance heterogeneity of known suggestive SNPs associated with height, BMI, T2D using UK Biobank.

| | SNP | Proxy SNPs | Distance to SNP | $R^2$ | Gene | Associated Traits | Associated P-values | Female | Male | Fisher | wM3V3.2 | wM3VNA3.3 |
|---|---|---|---|---|---|---|---|---|---|---|---|---|
| **BMI** | rs5945326 | | 0 | 1.00 | DUSP9 | T2D | 7.00E-16 | 1.96E-01 | 6.32E-03 | 9.53E-03 | 4.39E-03 | 1.22E-02 |
| | rs5906035 | | 0 | 1.00 | MIR221 | Height | 3.15E-04 | 7.91E-02 | 1.01E-01 | 4.65E-02 | 2.79E-02 | 5.24E-02 |
| | rs12010175 | | 0 | 1.00 | FAM58A | T2D | 2.00E-09 | 2.49E-01 | 2.57E-03 | 5.35E-03 | 5.51E-03 | 6.90E-03 |
| | | rs137887354 | 5666 | 1.00 | | | | 1.56E-01 | 5.29E-03 | 6.67E-03 | 1.11E-02 | 8.60E-03 |
| | | rs3817718 | -2181 | 1.00 | | | | 2.80E-01 | 4.30E-03 | 9.30E-03 | 8.94E-03 | 1.20E-02 |
| | rs1316982 | | | | NR | BMI | 3.00E-09 | | | | | |
| | | rs2248846 | 16431 | 1.00 | | | | **7.25E-04** | 7.12E-01 | 4.43E-03 | 2.05E-03 | 2.71E-03 |
| | | rs2254498 | -31632 | 1.00 | | | | 1.10E-03 | 5.87E-01 | 5.41E-03 | 2.83E-03 | 3.65E-03 |
| | | rs2489879 | 43560 | 1.00 | | | | **6.70E-04** | 6.62E-01 | 3.87E-03 | 2.40E-03 | 2.45E-03 |
| | | rs2495623 | 10755 | 1.00 | | | | **9.55E-04** | 6.59E-01 | 5.27E-03 | 2.56E-03 | 3.37E-03 |
| | | rs4825600 | -50763 | 1.00 | | | | 3.84E-03 | 5.07E-01 | 1.41E-02 | 5.25E-03 | 1.05E-02 |
| | | rs759147 | 58790 | 0.85 | | | | **1.79E-04** | 8.46E-01 | **1.49E-03** | **2.24E-04** | **7.73E-04** |
| | rs1586315 | | | | EDA2R | T2D | 9.28E-04 | | | | | |
| | | rs5919159 | 1479 | 1.00 | | | | 7.07E-02 | 7.94E-03 | 4.76E-03 | 2.09E-03 | 6.24E-03 |
| | | rs5919120 | -109901 | 1.00 | | | | 1.00E-01 | 5.78E-03 | 4.91E-03 | 2.20E-03 | 6.52E-03 |
| | | rs1385699 | 16391 | 0.84 | | | | 4.60E-01 | 8.01E-03 | 2.43E-02 | 1.34E-02 | 3.24E-02 |
| | | rs5965182 | -201902 | 0.80 | | | | 1.80E-02 | 1.17E-03 | **2.47E-04** | **1.28E-04** | **3.24E-04** |
| | rs5920868 | | | | NOX1 | Height | 5.51E-05 | | | | | |
| | | rs17323346 | -7349 | 1.00 | | | | 3.58E-01 | 1.44E-02 | 3.24E-02 | 3.69E-02 | 4.21E-02 |
| | rs5906035 | | 0 | 1.00 | MIR221 | Height | 3.15E-04 | 3.64E-02 | 7.49E-01 | 1.26E-01 | 3.90E-02 | 7.76E-02 |
| | rs1190736 | | 0 | 1.00 | GPR101 | BMI | 1.00E-08 | 3.87E-01 | 4.97E-02 | 9.53E-02 | 5.78E-02 | 1.27E-01 |
| | rs1474563 | | 0 | 1.00 | ITM2A | Height | 3.00E-06 | 3.60E-03 | 1.76E-01 | 5.30E-03 | **1.54E-03** | 4.10E-03 |
| | rs5945326 | | 0 | 1.00 | DUSP9 | T2D | 7.00E-16 | 1.01E-01 | 1.92E-02 | 1.40E-02 | 6.74E-03 | 1.82E-02 |
| | rs12010175 | | 0 | 1.00 | FAM58A | T2D | 2.00E-09 | 1.12E-01 | 6.87E-03 | 6.28E-03 | 6.05E-03 | 7.97E-03 |
| | | rs137887354 | 5666 | 1.00 | | | | 6.61E-02 | 1.50E-02 | 7.83E-03 | 1.12E-02 | 9.13E-03 |
| | | rs3817718 | -2181 | 1.00 | | | | 1.34E-01 | 1.32E-02 | 1.30E-02 | 1.05E-02 | 1.60E-02 |
| **Waist** | rs1316982 | | | | NR | BMI | 3.00E-09 | | | | | |
| | | rs2248846 | 16431 | 1.00 | | | | 1.26E-02 | 1.37E-01 | 1.27E-02 | 4.07E-03 | 1.08E-02 |
| | | rs2254498 | -31632 | 1.00 | | | | 1.95E-02 | 1.41E-01 | 1.89E-02 | 5.85E-03 | 1.58E-02 |
| | | rs2489879 | 43560 | 1.00 | | | | 1.08E-02 | 1.36E-01 | 1.11E-02 | 3.50E-03 | 9.20E-03 |
| | | rs2495623 | 10755 | 1.00 | | | | 1.18E-02 | 1.22E-01 | 1.09E-02 | 3.70E-03 | 9.43E-03 |
| | | rs4825600 | -50763 | 1.00 | | | | 2.74E-02 | 8.42E-02 | 1.63E-02 | 5.49E-03 | 1.52E-02 |
| | | rs759147 | 58790 | 0.85 | | | | 2.12E-03 | 1.31E-01 | 2.54E-03 | **5.34E-04** | 1.73E-03 |
| | rs1586315 | | | | EDA2R | T2D | 9.28E-04 | | | | | |
| | | rs5919159 | 1479 | 1.00 | | | | 1.79E-01 | **7.05E-04** | **1.26E-03** | **7.44E-04** | 2.38E-03 |
| **Hip** | | rs5919120 | -109901 | 1.00 | | | | 2.84E-01 | **7.63E-04** | 2.05E-03 | **1.21E-03** | 3.63E-03 |
| | | rs1385699 | 16391 | 0.84 | | | | 4.09E-01 | 1.06E-03 | 3.77E-03 | 2.19E-03 | 6.56E-03 |
| | | rs5965182 | -201902 | 0.80 | | | | 8.65E-02 | **5.57E-04** | **5.27E-04** | **6.89E-04** | **9.06E-04** |
| | rs1914711 | | | | AGTR2 | T2D | 1.00E-09 | | | | | |

| | | | | | | | | | | | | |
|---|---|---|---|---|---|---|---|---|---|---|---|---|
| | | rs5950474 | 1777 | 0.97 | | | | 6.87E-01 | 4.98E-02 | 1.50E-01 | 1.17E-01 | 2.10E-01 |
| | rs5920868 | | | | NOX1 | Height | 5.51E-05 | | | | | |
| | | rs17323346 | -7349 | 0.95 | | | | 1.78E-01 | 3.78E-02 | 4.04E-02 | 1.21E-01 | 4.79E-02 |
| | rs12010175 | | 0 | 1.00 | FAM58A | T2D | 2.00E-09 | 6.06E-02 | 9.52E-03 | 4.88E-03 | 1.99E-02 | 6.08E-03 |
| | | rs137887354 | 5666 | 1.00 | | | | 3.85E-02 | 1.27E-02 | 4.21E-03 | 2.55E-02 | 5.13E-03 |
| | | rs3817718 | -2181 | 1.00 | | | | 9.33E-02 | 1.44E-02 | 1.02E-02 | 3.24E-02 | 1.28E-02 |
| | rs1316982 | | | | NR | BMI | 3.00E-09 | | | | | |
| | | rs2254498 | -31632 | 1.00 | | | | 4.65E-02 | 4.75E-01 | 1.06E-01 | 4.24E-02 | 9.16E-02 |
| | | rs759147 | 58790 | 0.85 | | | | 2.17E-02 | 9.49E-01 | 1.00E-01 | 4.01E-02 | 6.03E-02 |
| | rs1586315 | | | | EDA2R | T2D | 9.28E-04 | | | | | |
| | | rs1385699 | 16391 | 0.84 | | | | 3.01E-01 | 4.27E-02 | 6.87E-02 | 7.43E-02 | 8.59E-02 |
| | | rs5965182 | -201902 | 0.80 | | | | 1.70E-01 | 6.67E-02 | 6.20E-02 | 3.37E-02 | 7.43E-02 |
| | rs570268 | | | | TBX22 | Height | 7.00E-04 | | | | | |
| | | rs113375614 | 6937 | 1.00 | | | | 3.73E-02 | 7.65E-01 | 1.30E-01 | 2.28E-01 | 1.01E-01 |
| | | rs589183 | -37688 | 1.00 | | | | 3.34E-02 | 6.90E-01 | 1.10E-01 | 2.24E-01 | 9.01E-02 |
| | | rs112732910 | -79200 | 0.84 | | | | 1.93E-02 | 2.55E-01 | 3.11E-02 | 1.30E-01 | 4.37E-02 |
| | rs1474563 | | 0 | 1.00 | ITM2A | Height | 3.00E-06 | 1.41E-02 | **2.51E-04** | **4.79E-05** | **5.87E-06** | **2.14E-05** |
| | rs1586315 | | | | EDA2R | T2D | 9.28E-04 | | | | | |
| | | rs5919159 | 1479 | 1.00 | | | | 4.52E-01 | 2.06E-02 | 5.28E-02 | 3.92E-02 | 7.34E-02 |
| | | rs5919120 | -109901 | 1.00 | | | | 3.81E-01 | 3.82E-02 | 7.61E-02 | 5.50E-02 | 1.01E-01 |
| | | rs1385699 | 16391 | 0.84 | | | | 3.48E-01 | 1.04E-02 | 2.40E-02 | 1.97E-02 | 3.40E-02 |
| | | rs5965182 | -201902 | 0.80 | | | | 7.17E-01 | 1.95E-02 | 7.37E-02 | 4.80E-02 | 1.06E-01 |
| **Height** | rs5922023 | | | | HDX | BMI | 2.20E-05 | | | | | |
| | | rs5922934 | -3795 | 1.00 | | | | 9.01E-01 | 4.66E-02 | 1.75E-01 | 1.34E-01 | 2.44E-01 |
| | rs6529684 | | | | HSD17B10, HUWE1 | BMI | 3.00E-08 | | | | | |
| | | rs1858002 | 107385 | 0.97 | | | | 1.40E-02 | 3.15E-01 | 2.84E-02 | 1.17E-01 | 2.29E-02 |
| | | rs57928918 | -53303 | 0.80 | | | | 2.87E-02 | 3.62E-01 | 5.79E-02 | 7.45E-02 | 4.74E-02 |

**S8 Table. Estimated genomic control lambda value ($\lambda_{GC}$) and estimated proportion of truly associated SNPs ($\pi_1$) using data from UKB and MESA.**

| | *Study* | *Trait* | *Female* | *Male* | *Fisher* | *wM3V3.2* | *wM3VNA3.3* |
|---|---|---|---|---|---|---|---|
| $\lambda_{GC}$ | UKB | Height* | 1.033 ($p < 0.1$) | 1.091 ($p < 0.1$) | 1.095 ($p < 0.1$) | 1.117 ($p < 0.1$) | 1.100 ($p < 0.1$) |
| | | Hip | 1.053 | 1.006 | 1.057 | 1.082 | 1.039 |
| | | BMI | 1.032 | 1.056 | 1.040 | 1.062 | 1.018 |
| | | Waist | 1.042 | 1.034 | 1.065 | 1.064 | 1.084 |
| | MESA | Height** | 1.128 ($p = 0.08$) | 1.060 ($p = 0.17$) | 1.183 ($p = 0.01$) | 1.192 ($p = 0.01$) | 1.171 ($p = 0.01$) |
| | | Hip | 1.063 | 1.032 | 1.045 | 1.045 | 1.027 |
| | | BMI | 1.002 | 1.028 | 1.024 | 1.005 | 0.997 |
| | | Waist | 0.975 | 1.139 | 1.063 | 1.143 | 1.081 |

| | *Study* | *Trait* | *Female* | *Male* | *Fisher* | *wM3V3.2* | *wM3VNA3.3* |
|---|---|---|---|---|---|---|---|
| $\pi_1$ | UKB | Height* | 0.044 ($p = 0.1$) | 0.068 ($p < 0.1$) | 0.099 ($p < 0.1$) | 0.124 ($p < 0.1$) | 0.091 ($p < 0.1$) |
| | | Hip | 0.038 | 0.024 | 0.048 | 0.044 | 0.026 |
| | | BMI | 0.017 | 0.010 | 0.013 | 0.034 | 0.005 |
| | | Waist | 0.029 | 0.015 | 0.050 | 0.070 | 0.062 |
| | MESA | Height** | 0.065 ($p = 0.17$) | 0.134 ($p < 0.01$) | 0.144 ($p < 0.01$) | 0.082 ($p = 0.13$) | 0.111 ($p = 0.04$) |
| | | Hip | 0.066 | 0.012 | 0.085 | 0.112 | 0.100 |
| | | BMI | 0.000 | 0.000 | 0.026 | 0.017 | 0.028 |
| | | Waist | 0.000 | 0.074 | 0.070 | 0.132 | 0.069 |

* The empirical *p*-values are based on a permutation study by sampling each quantitative trait without replacement within each sex in the UKB genetic data released in May 2017 ($n$ = 327,393), independently, **10** times.

**The empirical *p*-values are based on a permutation study by sampling each quantitative trait without replacement within each sex in the full MESA data ($n$ = 2,073), independently, **100** times.

### Supplementary Methods

#### 1 A statistical model for quantitative trait variance

Consider a normally distributed quantitative trait $Y$ measured on $N_f$ females and $N_m$ males with $N_f + N_m = n$. We assume that the three possible genotypes of a biallelic single nucleotide polymorphism (SNP) on the female X-chromosome (XCHR) are coded as $AA$, $Aa$ and $aa$, and the two male alleles are $A$ and $a$, with corresponding group sample sizes denoted by $n_{AA}$, $n_{Aa}$, $n_{aa}$ and $n_A$, $n_a$. Without loss of generality, $a$ is assumed to be the minor allele.

For autosomal SNPs, the genotype $G$ is coded additively, taking possible value of 0, 1, 2 according to the number of minor alleles without any ambiguity. For an XCHR SNP, we have the choice to denote the genotype (or the number of minor alleles) by $G = 0, 1, 2$ in females and $G = 0, 1$ in males, i.e. the additive genotype coding assuming no X-inactivation [1]. On the other hand, if the XCHR SNP were under complete X-inactivation, then the number of minor alleles is denoted by $G = 0, 1, 2$ in females and $G = 0, 2$ in males [1]. Note that although $G$ could be coded assuming X-inactivation or no X-inactivation, the two types of coding are generally highly correlated leading to similar association results [2]. Any theoretical derivations in the current manuscript focus on the no X-inactivation coding, as results for X-inactivation coding can be produced similarly.

Suppose the underlying true model of $Y$ has a linear relationship with main effects, pairwise, and three-way interaction effects with the genotype $G$, an environmental covariate $E$ of interest (either continuous or categorical, and additional covariates can be similarly included in the model), and the sex covariate ($S$, traditionally coded as 0 for female and 1 for male). Let true regression coefficients be denoted by $\beta$ with a subscript for each respective predictor:

$$Y \sim \beta_0 + \beta_G G + \beta_E E + \beta_S S + \beta_{GS} GS + \beta_{SE} SE + \beta_{GE} GE + \beta_{GSE} GSE + \epsilon \qquad (1)$$

We assume the continuous covariate of interest is normally distributed $E \sim \mathcal{N}(0, 1)$ and follows the classical gene-environment (G-E) independence assumption [3]. The error term is conventionally assumed to have a normal distribution $\epsilon \sim \mathcal{N}(0, 1)$, and is independent of $G$, $S$ and $E$. Note hat a statistical gene-environment interaction refers to the departure from a model that is linear in main effects. Under this model, we can write the variance and conditional variance of $Y$ as a function of the true regression coefficients. But first, we revisit variance heterogeneity of quantitative traits in autosomal SNPs and then move onto the more challenging scenario of XCHR SNPs.

##### 1.1 Variance heterogeneity: theoretical results for autosomal SNPs

For autosomal SNPs, it has been shown previously that the presence of an interaction between $G$ and $E$, under the standard assumption of G-E independence, could induce a difference in variance of subgroups stratified by $G$ if $Y$ were generated under a reduced linear model [4]:

$$Y \sim \beta_0 + \beta_G G + \beta_E E + \beta_S S + \beta_{GE} GE + \epsilon \qquad (2)$$

Essentially, the conditional distribution of $Y|G$ is expected to differ in mean if $\beta_G \neq 0$, and in variance if $\beta_{GE}$ were non-zero (SFig 1). For an autosomal SNP, the conditional variance can be shown to be a function of the marginal effects that do not involve $G$ (e.g. $\beta_E$) and the interaction with $G$ (e.g. $\beta_{GE}$). Thus, a non-zero interaction would create a variance difference between levels of $G$:

$$E(Y|G=g) = \beta_0 + \beta_G \Pr(G=g) \tag{3}$$

$$\mathrm{Var}(Y|G=g) = (\beta_E + \beta_{GE}\mathrm{g})^2 + 1. \tag{4}$$

The above expression implies that if $\beta_{GE} = 0$, the conditional variance would not vary with the genotypes, or the contrapositive, if the conditional variance varied with the genotype $G$, then $\beta_{GE}$ would be non-zero. This contrapositive statement provides the basis to leverage an observed variance heterogeneity so we could identify candidates SNPs that might be involved in genetic interactions. Note that another important assumption underlying this result is that $E(Y|G=g)$ and $\mathrm{Var}(Y|G=g)$ should be uncorrelated under the null hypothesis of $\beta_G = 0$ and $\beta_{GE} = 0$, i.e. the conditional mean should not be proportional to the conditional variance. Otherwise, a large $\beta_G$ could produce differences in variance that are not due to a non-zero $\beta_{GE}$ as has been cautioned in the literature [4, 5].

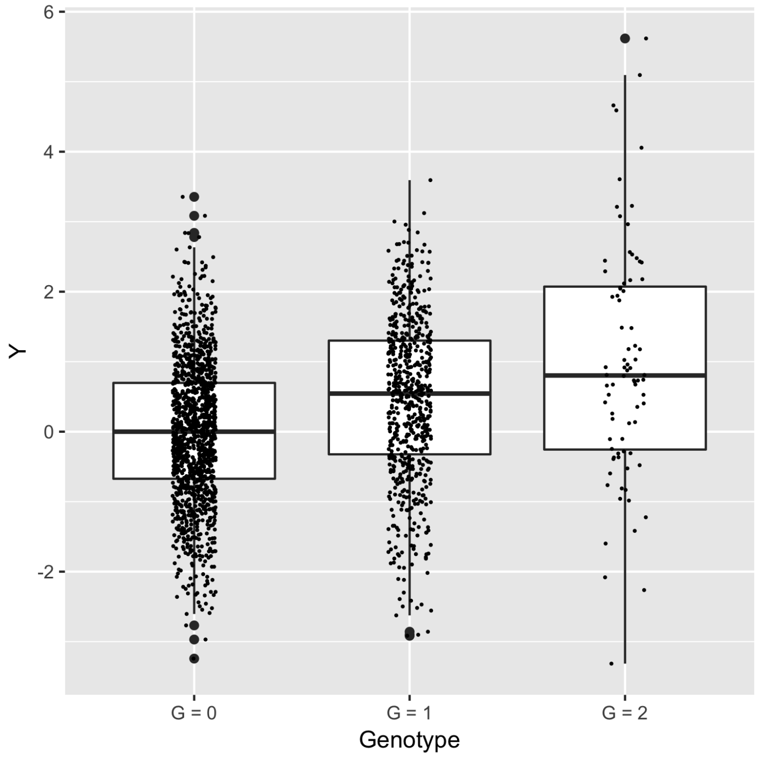

SFig 1: **Distribution of a quantitative trait stratified by genotypes of a biallelic SNP when both $\beta_G$ and $\beta_{GE}$ are non-zero.** The quantitative trait was simulated according to model 2 with a minor allel frequency of 0.2 and sample size of 2,000.

Indeed, Paré *et al.* [4] suggested to find potentially interacting SNPs by examining the variance heterogeneity of a quantitative trait stratified by $G$ for all SNPs without the need to directly measure any interacting covariates. The null hypothesis of variance homogeneity can be easily tested using the classical Levene's test [6]. Then, any SNPs with marginally significant Levene's test $p$-value are selected as likely candidates for statistical interactions with an environmental covariate or another SNP. Further testing of $G$x$E$ or $G$x$G$ interactions can be done in either the same study or followed up in a separate study where the observed covariates are available. Finally, by incorporating additional information, an optimal $p$-value threshold can be determined to declare a marginally significant variance heterogeneity that maximizes the power for interaction testing in follow-up analyses [7].

#### 1.2 Sex as a potential confounder on XCHR SNPs $G$-$Y$ associations

However, the same approach does not extend directly to XCHR SNPs despite the choices of coding for $G$ with respect to X-inactivation [1, 8]. Intuitively, knowing the sex of an individual gives information about the possible values of $G$, implying the possibility of $S$, or any other covariate that could be correlated with $S$, being a potential confounder on the association between $G$ and $\text{Var}(Y|G)$ whenever $Y$ depends on $S$.

In other words, here we observe the classic case of confounding in epidemiology between $G$ and $Y$ via $S$ through either the mean or variance of $Y$, as it is well known in human genetics that many complex traits exhibit sexual dimorphism, whereby the sex-stratified distributions of quantitative trait can be quite different (see Figure 1 of the manuscript). Since a quantitative trait can always be inverse-normally transformed, without loss of generality, we assume the trait is normally distributed such that the difference in distribution is mediated through either a mean (location) or variance (scale) difference. In this subsection, we show how correlation between $G$-$S$ and $S$-$Y$ can create spurious variance heterogeneity signals that are unrelated to the putative $G$x$E$ interaction.

##### 1.2.1 Correlation between $G$ and $S$

The sampling distribution of a genotype is conditional on the genetic sex of an individual for XCHR SNPs outside of the pseudo-autosomal region:

$$(G|S=0) \overset{D}{=} \text{rbinom}(p_{\text{F}}, 2) \quad \text{and} \quad (G|S=1) \overset{D}{=} \text{rbinom}(p_{\text{M}}, 1).$$

This condition necessarily creates a linear correlation between the observed genotype and sex. Let $n_{\text{F}}$ and $n_{\text{M}}$ denote the number of females and males, respectively, and $n = n_{\text{F}} + n_{\text{M}}$ be the combined sample size. Further, let the minor allele frequency (MAF) of an XCHR SNP be $p_{\text{F}}$ in females and $p_{\text{M}}$ in males. Let $r = \frac{n_{\text{M}}}{n}$ denote the proportion of males in the sample. The sample covariance between $G$ and $S$ can be simplified to a function of $r$, $p_{\text{M}}$, and $p_{\text{F}}$:

$$\begin{aligned}
\text{s}(G,S) &= \frac{\sum_{i=1}^{n} G_i S_i}{n} - \frac{\sum_{i=1}^{n} G_i}{n} \frac{\sum_{i=1}^{n} G_i}{n} \\
&= p_{\text{M}} r - \frac{2p_{\text{F}} n_{\text{F}} + p_{\text{M}} n_{\text{M}}}{N} r \\
&= (1-r) r p_{\text{M}} - 2(1-r) r p_{\text{F}} \\
&= (1-r) r (p_{\text{M}} - 2p_{\text{F}}),
\end{aligned} \tag{5}$$

where it is negative when $p_{\text{M}} = p_{\text{F}}$, and the maximum strength is achieved when $r = \frac{1}{2}$ (equal sample sizes in females and males) for any given MAF.

Since a scaled linear measure of dependence between $G$ and $S$ also depends on their variances, we give the variance of $S$ and $G$ in terms of $r$, $p_{\text{M}}$, and $p_{\text{F}}$:

$$s_{\text{S}}^2 = \frac{n_{\text{F}} n_{\text{M}}}{n^2} = (1-r)r \tag{6}$$

and

$$s_{\text{G}}^2 = \frac{n_{\text{M}}}{n} p_{\text{M}}(1-p_{\text{M}}) + 2\frac{n_{\text{F}}}{n} p_{\text{F}}(1-p_{\text{F}}) \tag{7}$$

Suppose further we have $p_{\text{F}} = p_{\text{M}} = p$, the correlation can be simplified to a function of $p$ and $r$ alone:

$$\begin{aligned} \text{cor}(G,S) &= \frac{(1-r)r(-p)}{\sqrt{(1-r)r}\sqrt{6p^2r+3pr-p^2r^2-4p^2}} \\ &= -\frac{\sqrt{(1-r)r}}{\sqrt{6r+3r/p-r^2-4}} \end{aligned} \tag{8}$$

The derivation results indicate that for XCHR SNPs, a non-zero linear correlation exist between the variables $G$ and $S$ whenever $p_{\text{F}} = p_{\text{M}}$. Surprisingly, the correlation between $G$ and $S$ is not symmetric about $r = \frac{1}{2}$ the same way for covariance. But a closer inspection reveals that the variance of $G$ also depends on $r$, so a small simulation check shows that correlation is negative and achieves its largest size when $p = 0.49$ and in the interval $r \in (0.58, 0.59)$. It also follows that the more unbalanced the sample sizes in females and males, the more variation around the correlation between $G$ and $S$ as the correlation depends on variance of both $G$ and $S$.

###### 1.2.2 Correlation between $Y$ and $S$: mean and variance of $Y$ stratified by $S$

The sampling distribution of quantitative trait $Y$ could be subjected to the genetic sex of an individual, a consequence of sexual dimorphism, which creates a linear dependence between the observed phenotype and sex. For many common complex traits, the sex-stratified distribution can be classified into one of the four categories provided that the trait had been normalized in the entire sample so only differences in the first two moments are of interest. For example, using complex traits collected in the multi-ethnic study of atherosclerosis (MESA) [9], the four categories corresponds to body mass index (BMI) with both mean and variance differences, height with a mean difference only, hip circumference with a variance difference only, and triglycerides with no clear signs of sexual dimorphism (SFig 2).

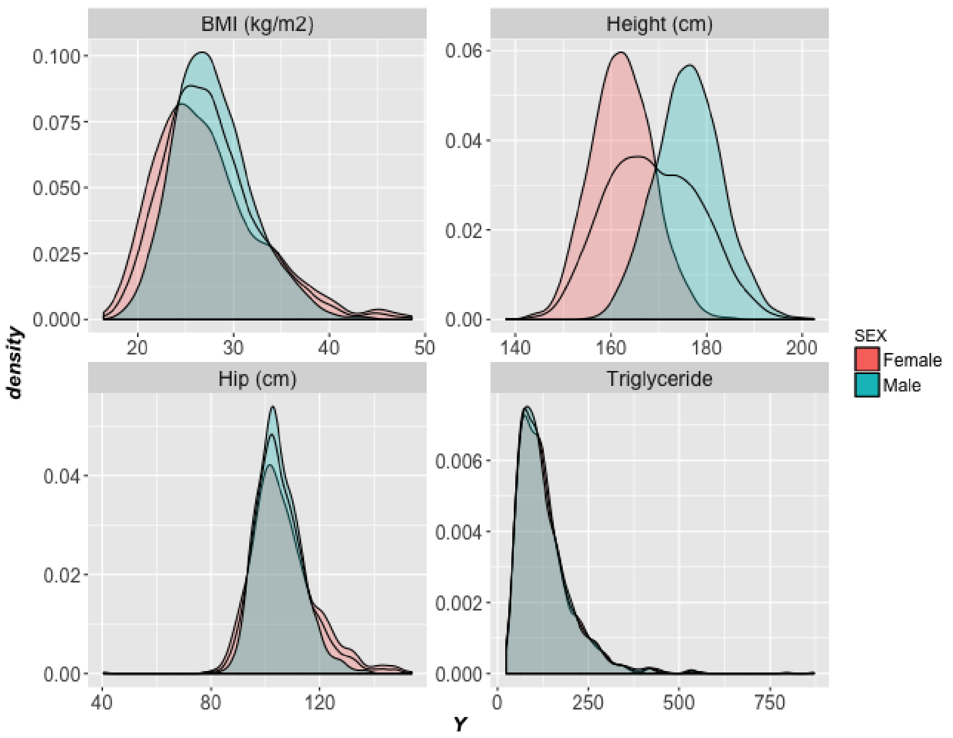

SFig 2: Sex-stratified distributions of complex traits using data from multi-ethnic study of atherosclerosis

Following the true model defined in Equation 1, we can write the conditional mean and variance of $Y$ on $S$:

$$\begin{aligned}\mathrm{E}(Y|S=s) &= \beta_0 + \beta_G \mathrm{E}(G|S=s) + \beta_E \mathrm{E}(E|S=s) + \beta_S s + \beta_{GS} \mathrm{E}(G|S=s)s \\ &+ \beta_{SE} \mathrm{E}(E|S=s)s + \beta_{GE} \mathrm{E}(GE|S=s) + \beta_{GES} \mathrm{E}(GE|S=s)s + \mathrm{E}(\epsilon|S=s) \\ &= \beta_0 + \beta_G \mathrm{E}(G|S=s) + \beta_S s + \beta_{GS} \mathrm{E}(G|S=s)s \\ &+ \beta_{GE} \mathrm{E}(GE|S=s) + \beta_{GES} \mathrm{E}(GE|S=s)s \\ &= \beta_0 + \beta_S s + (\beta_G + \beta_{GS} s)\mathrm{E}(G|S=s) \end{aligned} \tag{9a}$$

$$\begin{aligned}\mathrm{Var}(Y|S=s) &= \mathrm{Var}((\beta_G + \beta_{GS} s)G|S + (\beta_E + \beta_{SE} s)E|S + (\beta_{GE} + \beta_{GES} s)GE|S) \\ &= (\beta_E + \beta_{SE} s)^2 + \Big((\beta_G + \beta_{GS} s) + (\beta_{GE} + \beta_{GES} s)\Big)^2 \mathrm{Var}(G|S) \end{aligned} \tag{9b}$$

The true regression coefficient contributing directly to the difference in sex-stratified means of $Y$ is $\beta_S$, and indirectly through $\beta_G$ and $\beta_{GS}$ by a factor of the difference between $\mathrm{E}(G|S=0)$ and $\mathrm{E}(G|S=1)$. The true regression coefficients directly impacting the difference in sex-stratified variance between females and males are $\beta_{SE}$ and $\beta_E$, and indirectly through $\beta_G$, $\beta_{GS}$, $\beta_{GE}$ and $\beta_{GES}$, while the size of the differences is modulated through the conditional variance of the genotype

Var($G|S$). The implication of this result is that, in the absence of any genetic interaction effects ($\beta_{GE}$, $\beta_{GS}$, and $\beta_{GES}$), a spurious variance signal could arise due to a non-zero $\beta_{SE}$ or a large enough $\beta_G$. Notice that the variance signal due to a non-zero $\beta_G$ has been suggested by Ma *et al.*, [10], whenever random X-inactivation is expected as Var($G|S$) becomes a mixture of variance under incomplete dosage compensation.

The effect of sex on mean and variance of $Y$, when genetic effect is absent, i.e. $\beta_G = \beta_{GE} = \beta_{GS} = \beta_{GES} = 0$, can be quantified by examining the conditional mean and variance:

$$\mathrm{E}(Y|S=s) = \beta_0 + \beta_S s, \tag{10a}$$

$$\mathrm{Var}(Y|S=s) = (\beta_E + \beta_{SE}s)^2 + 1. \tag{10b}$$

These show that under the simple model without any genetic effects, $\beta_S$ fully accounts for the difference in mean, and $\beta_{SE}^2 + 2\beta_{SE}\beta E$ fully accounts for the difference in variance between sexes. In other words, the effect of $S$ on mean and variance could be adjusted by properly accounting for these two parameters. Indeed, including potential confounders in linear regression models as covariates is a common approach to reduce the effect of confounding on associations.

##### 1.2.3 Correlation between $Y$ and $G$: mean and variance of $Y$ stratified by $G$

The correlation between $G$ and $Y$ in terms of the linear model (Equation 1) can manifest as differences in either conditional mean, conditional variance or both when $Y$ is stratified by $G$. A stratified mean difference is of interest in the context of an XCHR-wide association for marginal effects, while a stratified variance difference is of interest, potentially, for an XCHR-wide variance heterogeneity analysis.

Without assuming $E-S$ independence, and further if we followed the coding scheme that $S=0$ is for male and $S=1$ for female (for mathematical convenience), we have the following expressions:

$$\begin{aligned}
\mathrm{E}(ES|G=g) &= \sum_s \mathrm{E}(E|G=g,S=s)\mathrm{Pr}(S=s|G=g) \\
&= \mathrm{E}(E|G=g,S=1)\mathrm{Pr}(S=1|G=g), && (11a)\\
\mathrm{Var}(ES|G=g) &= \mathrm{E}(E^2|G=g,S=1)\mathrm{Pr}(S=1|G=g) \\
&\quad - \mathrm{E}(E|G=g,S=1)^2\mathrm{Pr}(S=1|G=g)^2, && (11b)\\
\mathrm{Cov}(S,ES|G=g) &= \mathrm{E}(E|G=g,S=1)\mathrm{Pr}(S=1|G=g)\mathrm{Pr}(S=0|G=g) && (11c)\\
\mathrm{Cov}(E,ES|G=g) &= \mathrm{Pr}(S=1|G=g)[\mathrm{E}(E^2|G=g,S=1) \\
&\quad - \mathrm{E}(E|G=g,S=1)\mathrm{E}(S|G=g)] && (11d)\\
\mathrm{Cov}(E,S|G=g) &= \mathrm{Pr}(S=1|G=g)[\mathrm{E}(E|G=g,S=1) - \mathrm{E}(E|G=g)]. && (11e)
\end{aligned}$$

We can similarly write out the conditional mean and variance of $Y$ given $G$ using regression coefficients in Equation 1:

$$
\begin{aligned}
\mathrm{E}(Y|G=g) &= \beta_0 + \beta_G g + \beta_E \mathrm{E}(E|G=g) + \beta_S \mathrm{E}(S|G=g) + \beta_{GS} g \mathrm{E}(S|G=g) \\
&\quad + \beta_{SE}\mathrm{E}(ES|G=g) + \beta_{GE}\mathrm{E}(E|G=g)g + \beta_{GES}\mathrm{E}(ES|G=g)g + \mathrm{E}(\epsilon|G=g) \\
&= \beta_0 + \beta_G g + \mathrm{E}(E|G=g)(\beta_E + \beta_{GE} g) + \mathrm{E}(S|G=g)(\beta_S + \beta_{GS} g) \\
&\quad + \mathrm{E}(E|G=g, S=1)\mathrm{Pr}(S=1|G=g)(\beta_{SE} + \beta_{GES} g) \qquad (12a) \\
\mathrm{Var}(Y|G=g) &= \mathrm{Var}(\beta_G g + \beta_E(E|G=g) + \beta_S(S|G=g) + \beta_{GS} g(S|G=g) + \beta_{SE} \\
&\quad (ES|G=g) + \beta_{GE} g(E|G=g) + \beta_{GES} g(ES|G=g) + \epsilon|G=g) \\
&= \mathrm{Var}[(\beta_E + \beta_{GE} g)(E|G=g) + (\beta_S + \beta_{GS} g)(S|G=g) \\
&\quad + (\beta_{SE} + \beta_{GES} g)(ES|G=g) + \epsilon|G=g] \\
&= (\beta_E + \beta_{GE} g)^2 \mathrm{Var}(E|G=g) + (\beta_S + \beta_{GS} g)^2 \mathrm{Var}(S|G=g) \\
&\quad + (\beta_{SE} + \beta_{GES} g)^2 \mathrm{Var}(ES|G=g) + \mathrm{Var}(\epsilon|G=g) \\
&\quad + 2(\beta_E + \beta_{GE} g)(\beta_S + \beta_{GS} g)\mathrm{Cov}(E, S|G=g) \\
&\quad + 2(\beta_E + \beta_{GE} g)(\beta_{SE} + \beta_{GES} g)\mathrm{Cov}(E, ES|G=g) \\
&\quad + 2(\beta_S + \beta_{GS} g)(\beta_{SE} + \beta_{GES} g)\mathrm{Cov}(S, ES|G=g) \qquad (12b)
\end{aligned}
$$

Under the $G$-$E$ independence assumption, we can rearrange the above to

$$
\begin{aligned}
\mathrm{E}(Y|G=g) &= \beta_0 + \beta_G g + \mathrm{Pr}(S=1|G=g)(\beta_S + \beta_{GS} g) \\
&\quad + \mathrm{E}(E|S=1)\mathrm{Pr}(S=1|G=g)(\beta_{SE} + \beta_{GES} g) \qquad (13a) \\
\mathrm{Var}(Y|G=g) &= (\beta_E + \beta_{GE} g)^2 + 1 + (\beta_S + \beta_{GS} g)^2 \mathrm{Var}(S|G=g) \\
&\quad + (\beta_{SE} + \beta_{GES} g)^2 [\mathrm{E}(E^2|S=1)\mathrm{Pr}(S=1|G=g) \\
&\quad - \mathrm{E}(E|S=1)^2 \mathrm{Pr}(S=1|G=g)^2] \\
&\quad + 2(\beta_E + \beta_{GE} g)(\beta_S + \beta_{GS} g)[\mathrm{E}(E|S=1)\mathrm{Pr}(S=1|G=g)] + \\
&\quad + 2(\beta_E + \beta_{GE} g)(\beta_{SE} + \beta_{GES} g)[\mathrm{E}(E^2|S=1) \\
&\quad - \mathrm{E}(E|S=1)\mathrm{Pr}(S=1|G=g)]\mathrm{Pr}(S=1|G=g) \\
&\quad + 2(\beta_S + \beta_{GS} g)(\beta_{SE} + \beta_{GES} g)[\mathrm{E}(E|S=1)\mathrm{Pr}(S=1|G=g)\mathrm{Pr}(S=0|G=g)] \\
&\qquad (13b)
\end{aligned}
$$

Further, in the absence of any genetic effects, i.e. $\beta_G = \beta_{GE} = \beta_{GES} = \beta_{GS} = 0$, the conditional mean and variance can be simplified to

$$
\begin{aligned}
\mathrm{E}(Y|G=g) &= \beta_0 + \mathrm{Pr}(S=1|G=g)[\beta_S + \mathrm{E}(E|S=1)\beta_{SE}] \qquad (14a) \\
\mathrm{Var}(Y|G=g) &= \beta_E^2 + \beta_S^2 \mathrm{Var}(S|G=g) \\
&\quad + \beta_{SE}^2[\mathrm{E}(E^2|S=1)\mathrm{Pr}(S=1|G=g) - \mathrm{E}(E|S=1)^2\mathrm{Pr}(S=1|G=g)^2] \\
&\quad + 2\beta_E\beta_S \mathrm{E}(E|S=1)\mathrm{Pr}(S=1|G=g) + \\
&\quad + 2\beta_E\beta_{SE}[\mathrm{E}(E^2|S=1) - \mathrm{E}(E|S=1)\mathrm{Pr}(S=1|G=g)]\mathrm{Pr}(S=1|G=g) \\
&\quad + 2\beta_S\beta_{SE}\mathrm{E}(E|S=1)\mathrm{Pr}(S=1|G=g)\mathrm{Pr}(S=0|G=g) + 1 \qquad (14b)
\end{aligned}
$$

From the above expressions, both the conditional mean and variance are subjected to differ whenever $S$ and $G$ are correlated. Thus, without taking into account the $G$-$S$ correlation, we could be capturing a spurious signal due to a non-zero $\beta_S$ or $\beta_{SE}$ in either the marginal association test (mean) or the variance heterogeneity test (variance). To see this more clearly, we can simplify Equations 13a and 13b under the further assumption of $E-S$ independence conditional on $G$:

$$\mathrm{E}(Y|G=g) = \beta_0 + \beta_G g + \Pr(S=1|G=g)(\beta_S + \beta_{GS} g) \tag{15a}$$

$$\begin{aligned}\mathrm{Var}(Y|G=g) = {} & (\beta_E + \beta_{GE} g)^2 + 1 \\ & + \Pr(S=1|G=g)[(\beta_S + \beta_{GS} g)^2 \Pr(S=0|G=g) + \\ & + (\beta_{SE} + \beta_{GES} g)^2 + 2(\beta_E + \beta_{GE} g)(\beta_{SE} + \beta_{GES} g)].\end{aligned} \tag{15b}$$

Similarly, if we remove the genetic effects, the above can be simplified to

$$\mathrm{E}(Y|G=g) = \beta_0 + \Pr(S=1|G=g)\beta_S \tag{16a}$$

$$\mathrm{Var}(Y|G=g) = \beta_E^2 + \beta_S^2 \Pr(S=0|G=g) + (\beta_S + \beta_{SE})^2 \Pr(S=1|G=g) + 1. \tag{16b}$$

Under these strict assumptions, the conditional mean and conditional variance could vary across the possible genotypes to achieve substantial variance heterogeneity through $\beta_S$, $\beta_{SE}$, and $\beta_E$ as a result of the linear dependence between $G$ and $S$. Similar to the simplified variance stratified by sex, we make the conclusion that the effect of sex can not be simply adjusted away due to the fact that $\mathrm{E}(S|G=g)$ and $\mathrm{Var}(S|G=g)$ are not constant across genotypes. The amount of confounding due to the $S$-$Y$ association is controlled by the sizes of $\beta_S$ and $\beta_{SE}$, which then influence the expected conditional mean and variance of the quantitative trait when stratified by $G$.

##### 1.2.4 Correlation between $Y$ and $G \times S$: mean and variance of $Y$ stratified by $G \times S$

We have shown the effect of sex on the conditional variance of $Y|G$, implying that the stratified variance of $Y|G$ could be confounded by the effect of sex through either $\beta_S$ or $\beta_{SE}$. To further investigate the use of quantitative trait variance as a valid instrument for measuring unobserved genetic interactions, we examined the mean and variance of subgroups stratified by $G \times S$ under the full linear model (Equation 1). We further assume the independence between $S$ and $E$ conditional on $G$ as $E$ will be a confounder through $S$ otherwise.

$$\begin{aligned}\mathrm{E}(Y|G=g,S=s) &= \beta_0 + \beta_G g + \beta_E \mathrm{E}(E|G=g,S=s) + \beta_S s \\ &\quad + \beta_{GS} gs + \beta_{SE} s + \beta_{GE} \mathrm{E}(E|G=g,S=s) g \\ &\quad + \beta_{GES} gs \mathrm{E}(E|G=g,S=s) + \mathrm{E}(\epsilon|G=g,S=s) \\ &= \beta_0 + \beta_G g + \beta_S s + \beta_{GS} gs \end{aligned} \tag{17a}$$

$$\begin{aligned}\mathrm{Var}(Y|G=g,S=s) &= \mathrm{Var}(\beta_G g + \beta_E (E|G=g,S=s) + \beta_S s + \beta_{GS} gs + \\ &\quad + \beta_{SE} s(E|G=g,S=s) + \beta_{GE} g(E|G=g,S=s) \\ &\quad + \beta_{GES} gs(E|G=g,S=s) + \epsilon|G=g,S=s) \\ &= \mathrm{Var}((\beta_E + \beta_{SE} s + \beta_{GE} g \\ &\quad + \beta_{GES} gs)(E|G=g,S=s) + \epsilon|G=g,S=s) \\ &= \Big(\beta_E + \beta_{SE} s + \beta_{GE} g + \beta_{GES} gs\Big)^2 \mathrm{Var}(E|G=g,S=s) \\ &\quad + \mathrm{Var}(\epsilon|G=g,S=s) \\ &= \Big(\beta_E + \beta_{SE} s + \beta_{GE} g + \beta_{GES} gs\Big)^2 + 1 \end{aligned} \tag{17b}$$

The conditional variance is still not free from confounding of sex due to the presence of $\beta_{SE}$ as can be shown under the null hypothesis of no genetic interactions, i.e. $\beta_{GE} = \beta_{GES} = 0$:

$$\mathrm{Var}(Y|G=0,S=0) = \beta_E^2 + 1 \tag{18a}$$
$$\mathrm{Var}(Y|G=1,S=0) = \beta_E^2 + 1 \tag{18b}$$
$$\mathrm{Var}(Y|G=2,S=0) = \beta_E^2 + 1 \tag{18c}$$
$$\mathrm{Var}(Y|G=0,S=1) = (\beta_E + \beta_{SE})^2 + 1 \tag{18d}$$
$$\mathrm{Var}(Y|G=1,S=1) = (\beta_E + \beta_{SE})^2 + 1 \tag{18e}$$

We can observe from the conditional variance that there is a systematic difference in variance between male and female due to a non-zero $\beta_{SE}$. On the other hand, the $G$-$S$ interaction is completely absorbed in the conditional mean and does not appear in the conditional variance.

### 2 Strategies to test variance heterogeneity in X-chromosomal SNPs

For XCHR SNPs, the distribution of the observed genotype is conditional on $S$, thus deciding whether the maximum number of copies of an allele is either 1 for male or 2 for female. This potentially introduces confounding on the location (mean) or scale (variance) association of $Y$ and $G$ through $S$ as we have shown in the previous section. The confounding effects are realized through the presence of non-constant values of $\mathrm{E}(S|G=g)$ for mean and $\mathrm{Var}(S|G=g)$ for variance among the three possible levels of $G=g$ in the above formulae. Thus, variance heterogeneity of $Y|G$ or $Y|G,S$ in XCHR SNPs does not necessarily indicate the presence of a $G$-$E$ or $G$-$E$-$S$ interaction.

Below, we outline the two main approaches to test for variance heterogeneity considered in this manuscript and propose tailored strategies using these approaches for the unique settings of XCHR SNPs with sexual dimorphic traits.

#### 2.1 A brief review of Levene's test

We describe Levene's test as an example of the ANOVA type variance test for its flexibility to incorporate different grouping schemes, error variance structure, and existing methodology to perform meta-analysis [11]. The original Levene's test performs ANOVA on $\|y_i - \bar{y_i}\|$, the absolute deviation of each observation $y_{ij}$ from the $i$th sample group mean ($\bar{y_i}$). A variation of the Levene's test is Brown–Forsythe test [12], where the group median ($\tilde{y_i}$) is used instead of the group mean. The median option is recommended for its robustness to non-normally distributed traits and unbalanced group sizes, and thus is adopted here in all analyses using the ANOVA type Levene's test.

Let $k$ denote the number of subgroups ($k = 3$ for the three possible genotypes of an autosomal SNP), and the parameters are $\sigma^2_{G=0}$, $\sigma^2_{G=1}$, and $\sigma^2_{G=2}$. Levene's test assesses the null hypothesis:

$$H_o : \sigma^2_{G=0} = \sigma^2_{G=1} = \sigma^2_{G=2} = \sigma^2,$$

against the alternative hypothesis:

$$H_1 : \text{at least one of } \sigma^2_{G=0}, \sigma^2_{G=1}, \sigma^2_{G=2} \text{ does not equal to } \sigma^2.$$

Under the null hypothesis of variance homogeneity, Levene's test statistics (Equation 19) follows an $F$-distribution with $k-1$ and $N-k$ degrees of freedom:

$$L = \frac{(N-k)\sum_{i=0}^{k-1} n_i(\bar{Z}_i - \bar{\bar{Z}})^2}{(k-1)\sum_{i=0}^{k-1}\sum_{j=1}^{n_i}(z_{ij} - \bar{Z}_i)^2} \tag{19}$$

where $z_{ij} = \|y_{ij} - \bar{Y}_i\|$, $y_{ij}$ is the $j^{th}$ observation in the $i^{th}$ subgroup and $\bar{Y}_i$ the median of $Y_{ij}$. $\bar{Z}_i$ is the group mean of $Z_{ij}$, and $\bar{\bar{Z}}$ the overall mean of $Z_{ij}$.

#### 2.2 A generalized Levene's test under a two-stage regression framework

Alternatively, Levene [6] and others [13–15] had considered the generalized Levene's test. The regression-type Levene's test provides a flexible two-stage testing framework, whereby stage 1 produces the residuals by removing the location effects either due to any confounding variables; and stage 2 tests the absolute value of the residuals, a measure of spread from the location parameters, with a score that represents the grouping assignment in a linear regression. Thus, the correct type I error rates can be maintained by properly accounting for any confounder in stage 1 and stage 2. This framework can be easily adapted to accommodate sample correlation and group uncertainty if necessary [15].

Stage 1 can be seen to be equivalent to the mean/median adjustment in the original Levene's test without accounting for additional covariates. In stage 1, $Y$ is first adjusted for a shift in location within each group. The residuals can be calculated via the ordinary least squared (OLS) method or least absolute deviations (LAD) method. Since the group sample size might be different, it is recommended to use LAD when unbalanced sample sizes are expected [15].

Formally, let $C$ be a categorical variable indicating the grouping assignment (or a weighted score of it) and $C_1, C_2, \ldots, C_m$ are $m$ ($m < n$) covariates to be adjusted for. The mean/median stage consists of calculating the residual $r$:

$$r = \text{resid}(y \sim \alpha_0 + \beta_C C + \beta_{C_1} C_1 + \cdots + \beta_{C_m} C_m) \tag{20}$$

In stage two, the response is the absolutely residual from stage one estimated using LAD. We assess variance heterogeneity by testing the null hypothesis $H_o : \gamma_C = 0$:

$$\|r\| \sim \gamma_0 + \gamma_C \text{C} \tag{21}$$

#### 2.3 Variance heterogeneity testing strategies using original Levene's test and generalized Levene's test

Here we present statistical tests based on the two approaches for variance heterogeneity of XCHR SNPs and provide theoretical results to support the use of a tailored generalized Levene's test.

##### 2.3.1 Naive Approaches using Levene's test

To test variance heterogeneity using Levene's test on XCHR, a vital step is to determine the number and the assignment of subgroups for the calculation of variation within. A few testing strategies directly using Levene's test can be devised depending on the subgroup assignments.

**Lev3** The most straightforward group assignment is $G = 0, 1, 2$, which has been used for autosomal SNPs. But our theoretical results in Section 1 indicate the possibility of producing inflated results whenever $\beta_G$, $\beta_S$ and $\beta_{SE}$ are non-zero (Equation 16b).

**Lev5** Another option is to use the combined $G \times S$ as a five-level factor, and this can similarly produce an inflated number type I errors whenever $\beta_{SE}$ is non-zero (Equation 18e).

**Fisher** As commonly done in genome-wide analysis, a sex-specific test can be first performed in females and males separately, denoted by **LevM** and **LevF**. That is, we analyze the variance heterogeneity separately in females and males for XCHR SNPs using Levene's test assuming $G$ is the factor with 3-levels in females and 2-levels in males. We then combine the two $p$-values using Fisher's method and obtain an overall $p$-value, denoted by **Fisher**.

##### 2.3.2 Model-based approach using the generalized Levene's test

Suppose we have the full model as described above (Equation 1) with an unobserved environmental covariate $E$. The variable of interest is the genotype $G$ and possible sources of confounding include sex $S$ and its interaction with $G$, as has been shown that $G$x$S$ interaction creates variance differences when the trait is stratified by either $S$ or $G$ even in the absence of $G$x$E$ or $G$x$E$x$S$ interactions (Equation 15b).

Since the sample size might be low in the group with the lowest genotype count, it is recommended to use LAD for XCHR SNPs for unbalanced sample sizes [15]. In addition, the additive coding is assumed for stage 1, as it has been suggested that most genetic variance is additive [16].

In stage 1, the trait is adjusted for a shift in location (mean or median) using one of the three models (M1-M3) below:

**Mean/Median Tests**:

$$\text{M1} : r_1 = \text{resid}(y \sim \alpha_1 + \beta_{G_1} G) \tag{22a}$$

$$\text{M2} : r_2 = \text{resid}(y \sim \alpha_2 + \beta_{G_2} G + \beta_{S_2} S) \tag{22b}$$

$$\text{M3} : r_3 = \text{resid}(y \sim \alpha_3 + \beta_{G_3} G + \beta_{S_3} S + \beta_{GS_3} GS) \tag{22c}$$

In stage 2, the response ($d = \|r\|$) is the absolutely residual from stage one estimated using LAD. If the genotype effect is additive on the variance of the trait, the three models as shown below (V1-V3) can be used to assess variance heterogeneity by testing the regression coefficient $H_o : \gamma_G = 0$; as well as a 2-degree freedom test (V3.2) for $H_o : \gamma_{G_3} = \gamma_{GS_3} = 0$, which could be more powerful if higher order interaction involving $G$ is present.

**Variance Tests**:

$$\text{V1} : d \sim \gamma_1 + \gamma_{G_1}\text{G} \tag{23a}$$

$$\text{V2} : d \sim \gamma_2 + \gamma_{G_2}\text{G} + \gamma_{S_2}\text{S} \tag{23b}$$

$$\text{V3} : d \sim \gamma_3 + \gamma_{G_3}\text{G} + \gamma_{S_3}\text{S} + \gamma_{GS_3}\text{GS} \tag{23c}$$

If we do not restrict ourselves to additive effect on the variance, the following models assuming non-additive genetic effects (NAV1-3) can be deployed to assess the variance heterogeneity by testing whether $H_o : \gamma_{G1} = \gamma_{G2} = 0$ for any departure of group variance from the referenced major allele homozygote/major allele group, where $G1 = \mathbb{1}_{G=1}$ and $G2 = \mathbb{1}_{G=2}$. It is also possible to have a 3-degree freedom test (VNA3.3) for $H_o : \gamma_{G1} = \gamma_{G2} = \gamma_{G1S_3} = 0$ that can be more powerful when a higher-order interaction involving $G$ is present.

$$\text{VNA1} : d \sim \gamma_1 + \gamma_{G1_1}\text{G1} + \gamma_{G2_1}\text{G2} \tag{24a}$$

$$\text{VNA2} : d \sim \gamma_2 + \gamma_{G1_2}\text{G1} + \gamma_{G2_2}\text{G2} + \gamma_{S_2}\text{S} \tag{24b}$$

$$\text{VNA3} : d \sim \gamma_3 + \gamma_{G1_3}\text{G1} + \gamma_{G2_3}\text{G2} + \gamma_{S_3}\text{S} + \gamma_{G1S_3}\text{G1S} \tag{24c}$$

The number of tests resulted from all possible adjustments under a regression-type Levene's test is 14 = 9 (1-degree freedom additive tests) + 3 (2-degree freedom non-additive tests) + 2 (3-degree freedom non-additive and 2-degree freedom additive tests).

Notice that under the model assumptions given by Equation 1, the residuals when standardized by their variances follow a student's t distribution, which means that $d_1$, $d_2$, and $d_3$, under large sample approximation, can be considered as half-normal random variables. One interesting characteristics of a half-normal random variable is the mean ($\mu = \sqrt{\frac{2}{\pi}}\sigma$) and variance ($\sigma^2(1 - \frac{2}{\pi})$) both depend on $\sigma^2$, the variance of the original normal distribution.

This is particularly worrisome in the second stage of the ANOVA or linear regression as homogeneity of variance is assumed, which implies that the variance of $d$ should be constant. An extreme case of heteroscedasticity could be due to the variance of the trait differ greatly between females and males, as sometime is observed in complex traits such as hip and waist circumference. To alleviate this problem, we propose a weighted variance approach instead by using $d_w = d/w$, where $w = \mathbb{1}_F s_F + \mathbb{1}_M s_M$ and $s_F$ and $s_M$ denote the sample standard deviation of $Y$ in females and males, respectively. These tests are denoted by wM1V1-wM3V3.2 and wM1VNA1-wM3VNA3.3, respectively, for the 14 tests described above.

#### 2.4 Discussion on variance testing strategies

It is important to define the null and alternative hypotheses of interest in the context of each variance heterogeneity testing strategy. Our primary interest in variance heterogeneity of XCHR SNPs is to discover potential $G$-$E$ interactions similar done in autosomal SNPs. Our proposed strategy defines the null hypothesis in terms of the variance heterogeneity induced by either a $G$-$E$ interaction, $G$-$E$-$S$ interaction or the presence of both while accounting for the confounding due to possible sexual dimorphism (i.e. association between $Y$ and $S$). The necessity of this adjustment has been demonstrated in the previous section that sex and any linear terms involving sex introduce spurious variance heterogeneity signals. However, we want to clarity that genetic interactions or confounding due to sexual dimorphism are not the only mechanisms that lead to variance heterogeneity in XCHR SNPs. For example, as has been shown elsewhere [10] that it could be explained by a large marginal effect creating a larger variance in the heterozygote group due to random X-inactivation.

The proposed testing strategies and the corresponding null hypothesis are summarized in Table 1.

| Strategies | Shorthand | Model Used | Null hypothesis of the Test |
|---|---|---|---|
| Location Test (LAD) | M1 | $y \sim \alpha_1 + \beta_{G_1} G$ | $H_o$:$\beta_{G_1} = 0$ |
| | M2 | $y \sim \alpha_2 + \beta_{G_2} G + \beta_{S_2} S$ | $H_o$: $\beta_{G_2} = 0$ |
| | M3 | $y \sim \alpha_3 + \beta_{G_3} G + \beta_{S_3} S + \beta_{GS_3} GS$ | $H_o$: $\beta_{G_3} = 0$ |
| Variance Test (Additive Coding) | wM1V1 | $d_{w1} \sim \gamma + \gamma_{G_1}$ G | $H_o$: $\gamma_{G_1} = 0$ |
| | wM1V2 | $d_{w1} \sim \gamma_2 + \gamma_{G_2}$ G $+ \gamma_{S_2}$ S | $H_o$: $\gamma_{G_2} = 0$ |
| | wM1V3 | $d_{w1} \sim \gamma_3 + \gamma_{G_3}$ G $+ \gamma_{S_3}$ S $+ \gamma_{GS_3}$ GS | $H_o$: $\gamma_{G_3} = 0$ |
| | wM1V3.2 | $d_{w1} \sim \gamma_3 + \gamma_{G_3}$ G $+ \gamma_{S_3}$ S $+ \gamma_{GS_3}$ GS | $H_o$: $\gamma_{G_3} = \gamma_{GS_3} = 0$ |
| | wM2V1 | $d_{w2} \sim \gamma + \gamma_{G_1}$ G | $H_o$: $\gamma_{G_1} = 0$ |
| | wM2V2 | $d_{w2} \sim \gamma_2 + \gamma_{G_2}$ G $+ \gamma_{S_2}$ S | $H_o$: $\gamma_{G_2} = 0$ |
| | wM2V3 | $d_{w2} \sim \gamma_3 + \gamma_{G_3}$ G $+ \gamma_{S_3}$ S $+ \gamma_{GS_3}$ GS | $H_o$: $\gamma_{G_3} = 0$ |
| | wM2V3.2 | $d_{w2} \sim \gamma_3 + \gamma_{G_3}$ G $+ \gamma_{S_3}$ S $+ \gamma_{GS_3}$ GS | $H_o$: $\gamma_{G_3} = \gamma_{GS_3} = 0$ |
| | wM3V1 | $d_{w3} \sim \gamma + \gamma_{G_1}$ G | $H_o$: $\gamma_{G_1} = 0$ |
| | wM3V2 | $d_{w3} \sim \gamma_2 + \gamma_{G_2}$ G $+ \gamma_{S_2}$ S | $H_o$: $\gamma_{G_2} = 0$ |
| | wM3V3 | $d_{w3} \sim \gamma_3 + \gamma_{G_3}$ G $+ \gamma_{S_3}$ S $+ \gamma_{GS_3}$ GS | $H_o$: $\gamma_{G_3} = 0$ |
| | wM3V3.2 | $d_{w3} \sim \gamma_3 + \gamma_{G_3}$ G $+ \gamma_{S_3}$ S $+ \gamma_{GS_3}$ GS | $H_o$: $\gamma_{G_3} = \gamma_{GS_3} = 0$ |
| Variance Test (non-Additive Coding) | M1VNA1 | $d_{w1} \sim \gamma + \gamma_{G1_1}$ G1 $+ \gamma_{G2_1}$ G2 | $H_o$: $\gamma_{G1_1} = \gamma_{G2_1} = 0$ |
| | wM1VNA2 | $d_{w1} \sim \gamma_2 + \gamma_{G1_2}$ G1 $+ \gamma_{G2_2}$ G2 $+ \gamma_{S_2}$ S | $H_o$: $\gamma_{G1_2} = \gamma_{G2_2} = 0$ |
| | wM1VNA3 | $d_{w1} \sim \gamma_3 + \gamma_{G1_3}$ G1 $+ \gamma_{G2_3}$ G2 $+ \gamma_{S_2}$ S $+ \gamma_{G1S_3}$ G1S | $H_o$: $\gamma_{G1_3} = \gamma_{G2_3} = 0$ |
| | wM1VNA3.2 | $d_{w1} \sim \gamma_3 + \gamma_{G1_3}$ G1 $+ \gamma_{G2_3}$ G2 $+ \gamma_{S_2}$ S $+ \gamma_{G1S_3}$ G1S | $H_o$: $\gamma_{G1_3} = \gamma_{G2_3} = \gamma_{G1S_3} = 0$ |
| | wM2VNA1 | $d_{w2} \sim \gamma + \gamma_{G1_1}$ G1 $+ \gamma_{G2_1}$ G2 | $H_o$: $\gamma_{G1_1} = \gamma_{G2_1} = 0$ |
| | wM2VNA2 | $d_{w2} \sim \gamma_2 + \gamma_{G1_2}$ G1 $+ \gamma_{G2_2}$ G2 $+ \gamma_{S_2}$ S | $H_o$: $\gamma_{G1_2} = \gamma_{G2_2} = 0$ |
| | wM2VNA3 | $d_{w2} \sim \gamma_3 + \gamma_{G1_3}$ G1 $+ \gamma_{G2_3}$ G2 $+ \gamma_{S_3}$ S $+ \gamma_{G1S_3}$ G1S | $H_o$: $\gamma_{G1_3} = \gamma_{G2_3} = 0$ |
| | wM2VNA3.2 | $d_{w2} \sim \gamma_3 + \gamma_{G1_3}$ G1 $+ \gamma_{G2_3}$ G2 $+ \gamma_{S_3}$ S $+ \gamma_{G1S_3}$ G1S | $H_o$: $\gamma_{G1_3} = \gamma_{G2_3} = \gamma_{G1S_3} = 0$ |
| | wM3VNA1 | $d_{w3} \sim \gamma + \gamma_{G1_1}$ G1 $+ \gamma_{G2_1}$ G2 | $H_o$: $\gamma_{G1_1} = \gamma_{G2_1} = 0$ |
| | wM3VNA2 | $d_{w3} \sim \gamma_2 + \gamma_{G1_2}$ G1 $+ \gamma_{G2_2}$ G2 $+ \gamma_{S_2}$ S | $H_o$: $\gamma_{G1_2} = \gamma_{G2_2} = 0$ |
| | wM3VNA3 | $d_{w3} \sim \gamma_3 + \gamma_{G1_3}$ G1 $+ \gamma_{G2_3}$ G2 $+ \gamma_{S_3}$ S $+ \gamma_{G1S_3}$ G1S | $H_o$: $\gamma_{G1_3} = \gamma_{G2_3} = 0$ |
| | wM3VNA3.2 | $d_{w3} \sim \gamma_3 + \gamma_{G1_3}$ G1 $+ \gamma_{G2_3}$ G2 $+ \gamma_{S_3}$ S $+ \gamma_{G1S_3}$ G1S | $H_o$: $\gamma_{G1_3} = \gamma_{G2_3} = \gamma_{G1S_3} = 0$ |
| Levene's Test (median option) | Lev3 | $y \sim \mathbb{1}_{G=1} + \mathbb{1}_{G=2}$ | $H_o$: $\sigma^2_{AA+A} = \sigma^2_{Aa+a} = \sigma^2_{aa}$ |
| | Lev5 | $y \sim \mathbb{1}_{G=1,S=0} + \mathbb{1}_{G=2,S=0} + \mathbb{1}_{G=1,S=1} + \mathbb{1}_{G=0,S=1}$ | $H_o$: $\sigma^2_{AA} = \sigma^2_{Aa} = \sigma^2_{aa} = \sigma^2_{A} = \sigma^2_{a}$ |
| | LevM | $y_{\text{Male}} \sim \mathbb{1}_{G=1}$ | $H_o$: $\sigma^2_{A} = \sigma^2_{a}$ |
| | LevF | $y_{\text{Female}} \sim \mathbb{1}_{G=1} + \mathbb{1}_{G=2}$ | $H_o$: $\sigma^2_{AA} = \sigma^2_{Aa} = \sigma^2_{aa}$ |
| | Fisher | $\chi^2_4 \sim -2(\ln(\text{LevM}) + \ln(\text{LevF}))$ | $H_o$: $\sigma^2_{AA} = \sigma^2_{Aa} = \sigma^2_{aa}$ and $\sigma^2_{A} = \sigma^2_{a}$ |

STab 1: A summary of variance heterogeneity testing strategies for XCHR SNPs

We can draw some theoretical conclusions on the equivalence of some tests under the null hypothesis. For example, **Lev3** is equivalent to **M1VNA1** (without the sex-stratified variance adjustment) with the LAD option corresponding to using median and OLS corresponding to using mean as the group centre for Levene's test.

Similarly, the five subgroup **Lev5** is equivalent to **M3VNA3.3** without the sex-stratified variance adjustment, but testing the null hypothesis of $\gamma_{G1} = \gamma_{G2} = \gamma_{G1S} = \gamma_S = 0$ as supposed to $\gamma_{G1} = \gamma_{G2} = \gamma_{G1S} = 0$.

Finally, the Fisher's method to combine the sex-specific Levene's test $p$-value can be seen as the equivalent of **wM3VNA3.3**. It is clear that wM3 is the only location test that can guard against

1) any sex-difference in mean and variance of the trait, 2) uncertainty of X-inactivation, and 3) possibly allelic heterogeneity between sexes for mean associations.

To see the equivalence between Fisher's method and **wM3VNA3.3**, we first briefly show that the null hypotheses under consideration are equivalent, and then show under the null scenario that the test statistics for the sex-specific case are captured by the test statistic resulted from **M3VNA3.3**. Finally, the sex-stratified weighted approach **wM3VNA3.3** combines test statistics from females and males linearly while accounting for the possible variance differences.

For the non-additive tests, we can express variance in each of the sub-group with respect to the parameters in the null hypothesis:

| | Null Hypothesis | $\sigma^2_{AA}$ | $\sigma^2_{Aa}$ | $\sigma^2_{aa}$ | $\sigma^2_{A}$ | $\sigma^2_{a}$ |
|---|---|---|---|---|---|---|
| **NAV1** | $\gamma_{G_1} = \gamma_{G_2} = 0$ | $\gamma_1$ | $\gamma_1 + \gamma_{G_1}$ | $\gamma_1 + \gamma_{G_2}$ | $\gamma_1$ | $\gamma_1 + \gamma_{G_1}$ |
| **NAV2** | $\gamma_{G_1} = \gamma_{G_2} = 0$ | $\gamma_2 + \gamma_S$ | $\gamma_2 + \gamma_S + \gamma_{G_1}$ | $\gamma_2 + \gamma_S + \gamma_{G_2}$ | $\gamma_2$ | $\gamma_2 + \gamma_{G_1}$ |
| **NAV3** | $\gamma_{G_1} = \gamma_{G_2} = 0$ | $\gamma_3 + \gamma_S$ | $\gamma_3 + \gamma_S + \gamma_{G_1} + \gamma_{G_1 S}$ | $\gamma_3 + \gamma_S + \gamma_{G_2}$ | $\gamma_3$ | $\gamma_3 + \gamma_{G_1}$ |
| **NAV3.3** | $\gamma_{G_1} = \gamma_{G_2} = \gamma_{G_1 S} = 0$ | $\gamma_3 + \gamma_S$ | $\gamma_3 + \gamma_S + \gamma_{G_1} + \gamma_{G_1 S}$ | $\gamma_3 + \gamma_S + \gamma_{G_2}$ | $\gamma_3$ | $\gamma_3 + \gamma_{G_1}$ |

STab 2: A summary of non-additive tests and the subgroup variances

Clearly, the null hypothesis of $\gamma_{G_1} = \gamma_{G_2} = \gamma_{G_1 S} = 0$ can be reduced to the null hypothesis tested by Fisher's method $H_o : \sigma^2_{AA} = \sigma^2_{Aa} = \sigma^2_{aa}$ and $\sigma^2_A = \sigma^2_a$.

To be consistent, we chose to illustrate the equivalence of the test statistics under the mean option for both Fisher's method and M3VNA3.3. In stage 1, the estimated group centres from sex-specific Levene's test are denoted by $\hat{\mu}_{F0}, \hat{\mu}_{F1}, \hat{\mu}_{F2}$ and $\hat{\mu}_{M0}, \hat{\mu}_{M1}$. Let the residuals in females be denoted by $d_F = \|y_F - \hat{\mu}_F\|$, while the residuals per genotype group in females denoted by $d_{F0} = \|y_{F0} - \hat{\mu}_{F0}\|$, $d_{F1} = \|y_{F1} - \hat{\mu}_{F1}\|$ and $d_{F2} = \|y_{F2} - \hat{\mu}_{F2}\|$, respectively. Similarly define $d_M$, $d_{M0}$ and $d_{M1}$ for males. The sum of squares in the combined sample can be decomposed:

$$
\begin{aligned}
\sum_{i=1}^{n}(d_i - \bar{d})^2 &= \sum_{i=1}^{n_{AA}}(d_{F0(i)} - \bar{d})^2 + \sum_{i=1}^{n_{Aa}}(d_{F1(i)} - \bar{d})^2 + \sum_{i=1}^{n_{aa}}(d_{F2(i)} - \bar{d})^2 \\
&\sum_{i=1}^{n_A}(d_{M0(i)} - \bar{d})^2 + \sum_{i=1}^{n_a}(d_{M1(i)} - \bar{d})^2 \\
&= \sum_{i=1}^{n_{AA}}(d_{F0(i)} - \bar{d}_{F0})^2 + + \sum_{i=1}^{n_{Aa}}(d_{F1(i)} - \bar{d}_{F1})^2 + \sum_{i=1}^{n_{aa}}(d_{F2(i)} - \bar{d}_{F2})^2 \\
&\sum_{i=1}^{n_A}(d_{M0(i)} - \bar{d}_{M0})^2 + \sum_{i=1}^{n_a}(d_{M1(i)} - \bar{d}_{M1})^2 + n_{AA}(\bar{d}_{F0} - \bar{d})^2 \\
&+ n_{Aa}(\bar{d}_{F1} - \bar{d})^2 + n_{aa}(\bar{d}_{F2} - \bar{d})^2 + n_A(\bar{d}_{M0} - \bar{d})^2 + n_a(\bar{d}_{M1} - \bar{d})^2 \\
&= \text{RSS}_{\text{Female}} + \text{RSS}_{\text{Male}} + \text{Model}_{\text{Female}} + \text{Model}_{\text{Male}} \qquad (25)
\end{aligned}
$$

For stage 1 of the M3VNA3.3 test, we have the linear model:

$$y \sim \beta_3 + \beta_{G_3} G + \beta_{S_3} S + \beta_{GS_3} GS. \qquad (26)$$

The variances are not equal among the five groups defined by the combinations of $G$ and $S$ under the linear model (Equation 1) according to (Equation 18e). Note that we must have

$\mathrm{Var}(r_3|G = g, S = s) = \mathrm{Var}(y|G = g, S = s)$, which means the OLS estimates though are still unbiased, but the true variance of the estimates might be underestimated. However, since our main objective in the first stage is not inference, but rather estimating an accurate location shift for each subgroup, this does not invalidate our procedure. If we used the dummy coding for sex, i.e. $S = 1$ indicates the male gender while $S = 0$ is the female sex, then the location effect for each subgroup is

$$\text{Female AA} : \bar{m}_{F0} = \hat{\beta}_3 \tag{27a}$$
$$\text{Female Aa} : \bar{m}_{F1} = \hat{\beta}_3 + \hat{\beta}_{G_3} \tag{27b}$$
$$\text{Female aa} : \bar{m}_{F2} = \hat{\beta}_3 + 2\hat{\beta}_{G_3} \tag{27c}$$
$$\text{Male A} : \bar{m}_{M0} = \hat{\beta}_3 + \hat{\beta}_{S_3} \tag{27d}$$
$$\text{Male a} : \bar{m}_{M1} = \hat{\beta}_3 + \hat{\beta}_{S_3} + \hat{\beta}_{G_3} + \hat{\beta}_{GS_3} \tag{27e}$$

or equivalently, if we used the coding scheme of $S = 2$ for females, the above location effect estimates become:

$$\text{Female AA} : \bar{m}_{F0} = \hat{\beta}_3 + 2\hat{\beta}_{S_3} \tag{28a}$$
$$\text{Female Aa} : \bar{m}_{F1} = \hat{\beta}_3 + 2\hat{\beta}_{S_3} + \hat{\beta}_{G_3} \tag{28b}$$
$$\text{Female aa} : \bar{m}_{F2} = \hat{\beta}_3 + 2\hat{\beta}_{S_3} + 2\hat{\beta}_{G_3} \tag{28c}$$
$$\text{Male A} : \bar{m}_{M0} = \hat{\beta}_3 + \hat{\beta}_{S_3} \tag{28d}$$
$$\text{Male a} : \bar{m}_{M1} = \hat{\beta}_3 + \hat{\beta}_{S_3} + \hat{\beta}_{G_3} + \hat{\beta}_{GS_3} \tag{28e}$$

Taking the residuals from stage 1, the models in stage 2 aim to test the association between the absolute residuals with G either without any adjustments or while accounting for S or GS. The variance of residuals from M3 can be expressed through the law of total variance:

$$\mathrm{Var}(r_3) = \mathrm{Var}(\mathrm{E}(r_3|G, S)) + \mathrm{E}(\mathrm{Var}(r_3|G, S)) = \mathrm{E}(\mathrm{Var}(r_3|G, S)), \tag{29}$$

and thus it is sufficient to look at the conditional variance. Since the $S$-$E$ interaction could create a variance difference between females and males, a problem is present in the second stage as $r_{3_{(i)}}$ will have a variance difference in sex. In this case, the error variance $\sigma^2$ depends on the group each observation falls into. Let $\sigma^2(G, S)$ denote the error variance and $\hat{\sigma}^2(G, S)$ denote the estimated error variance in each group defined by combinations of $G$ and $S$, where

$$\hat{\sigma}^2(G = g, S = s) = \frac{\sum_{i=1}^{n} r_{3_{(i)}}^2 \mathbb{1}_{G_i=g} \mathbb{1}_{S_i=s}}{\sum_{i=1}^{n} \mathbb{1}_{G_i=g} \mathbb{1}_{S_i=s} - 1}$$

.

The studentized residual

$$\frac{r_{3_{(i)}}}{\hat{\sigma}(G = g, S = s)\sqrt{1 - h_{ii}}}$$

has a t-distribution with degrees of freedom $n - 5$ (5 is the number of parameters), where $h_{ii}$ is the $i$th diagonal entry of the hat matrix $H = X(X^T X)^{-1} X^T$ and $X$ is the design matrix. As all predictors in this model are categorical variables, there are only 6 distinct values in the hat matrix (one for the intercept and one for each of the five groups).

Asymptotically, the studentized residual has a normal distribution with mean 0 and variance 1 as the sample size is usually in the thousands or tens of thousands for large genome-wide studies. This then implies that the absolute value of the studentized residual,

$$\frac{d_{3_{(i)}}}{\hat{\sigma}(G=g,S=s)\sqrt{1-h_{ii}}},$$

has a half-normal distribution with mean $\sqrt{2/\pi}$ and variance $(1-\frac{2}{\pi})$. Clearly, the mean and variance are correlated and the estimates will be biased if we use $d_{3_{(i)}}$ as the response without accounting for the systematic variance difference in $\hat{\sigma}(G=g,S=s)$ due to sex. Under the null hypothesis of no genetic effects, the error variance $\sigma^2(G,S)$ really only depends on the size of $\beta_{SE}$. In other words, any adjustment that goes beyond the sex-stratified variances will lead to an under-estimate of the variance heterogeneity, but no adjustments for sex-stratified variances will bias the results.

To see how it will impact the validity of test, we look at the model without weights first. For stage 2 of the test, we have the linear regression model based on $d_{3(i)}$,

$$d_3 \sim \gamma_3 + \gamma_{G1_3}\mathrm{G1} + \gamma_{G2_3}\mathrm{G2} + \gamma_{S_3}\mathrm{S} + \gamma_{G1S_3}\mathrm{G1S}, \tag{30}$$

to test the null hypothesis of $\gamma_{G1_3} = \gamma_{G2_3} = \gamma_{G1S_3} = 0$.

Let $\hat{d}_{3(i)} = \hat{\gamma}_3 + \hat{\gamma}_{G1_3}\mathrm{G1}_i + \hat{\gamma}_{G2_3}\mathrm{G2}_i + \hat{\gamma}_{S_3}\mathrm{S}_i + \hat{\gamma}_{G1S_3}\mathrm{G1}_i\mathrm{S}_i$ denote the fitted absolute residual. Further, define $d_{3(i)FO}$, $d_{3(i)F1}$, $d_{3(i)F2}$, $d_{3(i)MO}$ and $d_{3(i)M1}$ to be the $i$th absolute residual in female $A/A$, $A/a$, $a/a$, male $A$, and male $a$ genotype/allele group. The sum of squares in the combined sample can be decomposed:

$$\begin{aligned}
\sum_{i=1}^{n}(d_{3(i)} - \hat{d}_{3(i)})^2 &= \sum_{i=1}^{n_{AA}}(d_{3(i)FO} - \hat{d}_{3(i)})^2 + \sum_{i=1}^{n_{Aa}}(d_{3(i)F1} - \hat{d}_{3(i)})^2 + \sum_{i=1}^{n_{aa}}(d_{3(i)F2} - \hat{d}_{3(i)})^2 \\
&+ \sum_{i=1}^{n_{A}}(d_{3(i)MO} - \hat{d}_{3(i)})^2 + \sum_{i=1}^{n_{a}}(d_{3(i)M1} - \hat{d}_{3(i)})^2 \\
&= \sum_{i=1}^{n_{AA}}(d_{3(i)FO} - \bar{m}_{F0})^2 + \sum_{i=1}^{n_{Aa}}(d_{3(i)F1} - \bar{m}_{F1})^2 + \sum_{i=1}^{n_{aa}}(d_{3(i)F2} - \bar{m}_{F2})^2 \\
&+ \sum_{i=1}^{n_{A}}(d_{3(i)MO} - \bar{m}_{M0})^2 + \sum_{i=1}^{n_{a}}(d_{3(i)M1} - \bar{m}_{M1})^2 + n_{AA}(\bar{m}_{F0} - \hat{d}_{3(i)})^2 \\
&+ n_{Aa}(\bar{m}_{F1} - \hat{d}_{3(i)})^2 + n_{aa}(\bar{m}_{F2} - \hat{d}_{3(i)})^2 + n_{A}(\bar{m}_{M0} - \hat{d}_{3(i)})^2 \\
&+ n_{a}(\bar{m}_{M1} - \hat{d}_{3(i)})^2 \\
&= \mathrm{RSS}_{\mathrm{Female}} + \mathrm{RSS}_{\mathrm{Male}} + \mathrm{Model}_{\mathrm{Female}} + \mathrm{Model}_{\mathrm{Male}}
\end{aligned} \tag{31}$$

For stage 2 of the test, we can also construct a linear regression model based on $d_{w3(i)} = d_{3(i)}/w_i$, where $w_i = \mathbb{1}_{S_i \neq 1}\hat{\sigma}_F + \mathbb{1}_{S_i=1}\hat{\sigma}_M$:

$$d_{w3} \sim \gamma_3 + \gamma_{G1_3}\mathrm{G1} + \gamma_{G2_3}\mathrm{G2} + \gamma_{S_3}\mathrm{S} + \gamma_{G1S_3}\mathrm{G1S} \tag{32}$$

to test the null hypothesis of $\gamma_{G1_3} = \gamma_{G2_3} = \gamma_{G1S_3} = 0$.

The sum of squares in the combined sample can be decomposed:

$$\sum_{i=1}^{n}(d_{w3(i)} - \hat{d}_{w3(i)})^2 = \frac{1}{\hat{\sigma}_F^2}\Big(\sum_{i=1}^{n_{AA}}(d_{3(i)FO} - \hat{d}_{3(i)})^2 + \sum_{i=1}^{n_{Aa}}(d_{3(i)F1} - \hat{d}_{3(i)})^2 + \sum_{i=1}^{n_{aa}}(d_{3(i)F2} - \hat{d}_{3(i)})^2\Big)$$
$$+ \frac{1}{\hat{\sigma}_M^2}\Big(\sum_{i=1}^{n_A}(d_{3(i)MO} - \hat{d}_{3(i)})^2 + \sum_{i=1}^{n_a}(d_{3(i)M1} - \hat{d}_{3(i)})^2\Big)$$
$$= \frac{1}{\hat{\sigma}_F^2}\Big(\sum_{i=1}^{n_{AA}}(d_{3(i)FO} - \bar{m}_{F0})^2 + \sum_{i=1}^{n_{Aa}}(d_{3(i)F1} - \bar{m}_{F1})^2 + \sum_{i=1}^{n_{aa}}(d_{3(i)F2} - \bar{m}_{F2})^2\Big)$$
$$+ \frac{1}{\hat{\sigma}_M^2}\Big(\sum_{i=1}^{n_A}(d_{3(i)MO} - \bar{m}_{M0})^2 + \sum_{i=1}^{n_a}(d_{3(i)M1} - \bar{m}_{M1})^2\Big)$$
$$+ \frac{1}{\hat{\sigma}_F^2}\Big(n_{AA}(\bar{m}_{F0} - \hat{d}_{3(i)})^2 + n_{Aa}(\bar{m}_{F1} - \hat{d}_{3(i)})^2 + n_{aa}(\bar{m}_{F2} - \hat{d}_{3(i)})^2\Big)$$
$$+ \frac{1}{\hat{\sigma}_M^2}\Big(n_A(\bar{m}_{M0} - \hat{d}_{3(i)})^2 + n_a(\bar{m}_{M1} - \hat{d}_{3(i)})^2\Big)$$
$$= \frac{1}{\hat{\sigma}_F^2}\mathrm{RSS}_{\mathrm{Female}} + \frac{1}{\hat{\sigma}_M^2}\mathrm{RSS}_{\mathrm{Male}} + \frac{1}{\hat{\sigma}_F^2}\mathrm{Model}_{\mathrm{Female}} + \frac{1}{\hat{\sigma}_M^2}\mathrm{Model}_{\mathrm{Male}} \quad (33)$$

Clearly, the location estimates will be equivalent to the estimated group centres for sex-specific mean when the genetic effects, $\beta_G = \beta_{GS_3} = 0$. However, when they are not zero, the difference will be small as most genetic effects contribute additively. The five group centres estimates can be adequately approximated by the four parameters assuming an additive coding of $G$ through $\beta_G$. Thus, the location adjustment of sex-specific Levene's tests and that of the M3 models are equivalent.

In the second stage, without any weights adjustment, the residual sum of squares from the M3VNA3.3 model is simply the sum of the residual sum of squares from the sex-specific Levene's test, while the model sum of squares are simply the sum of the model sum of squares from the sex-specific Levene's test. Thus, the evidence from the combined sex-specific Fisher's method and the M3VNA3.3 differs only in the way the $p$-values are calculated according to the null distribution. The advantages of the Fisher's method is that sex-specific variance does not influence the $p$-values, while the disadvantage is that it is observed to be consistently underpowered than wM3VNA3.3 in simulations under plausible scenarios.
